# A geometric and dynamical theory of latent computations in biological neural networks

**DOI:** 10.64898/2026.07.10.737763

**Authors:** Fatih Dinc, Marta Blanco-Pozo, David Klindt, Francisco Acosta, Cameron Sylber, Yiqi Jiang, Sadegh Ebrahimi, Adam Shai, Hidenori Tanaka, Peng Yuan, Nina Miolane, Mark J. Schnitzer

## Abstract

Many neural recordings have revealed low-dimensional sets of behaviorally relevant variables encoded within large-scale neural activity patterns. However, dimensionality reduction analyses alone cannot yield causal explanations for how networks stably implement computations that are resilient to the substantial variability of single neuron dynamics. Further, existing methods for dimensionality reduction often rely on simplifying assumptions about network structure that limit their applicability and explanatory power. To provide a theoretical framework describing the dynamics of low-dimensional computation in high-dimensional neural networks, here we introduce the concept of latent processing units (LPUs), which are architecture-agnostic computational elements operating within biological neural circuitry. Six theorems governing coding and computation by LPUs collectively provide explanations for a range of common biological findings: low-dimensional sets of coding variables can generate high-dimensional neural dynamics; many neurons have activity patterns that represent behaviorally relevant variables but exert little influence on downstream circuits; linear readouts of neural population activity commonly permit near-optimal decoding; the drift of neural representations is often substantial even while network computations remain intact. Overall, our treatment of LPUs, as enacted in network dynamics, unifies the geometric and dynamical views of neural computation under a joint framework and provides systems neuroscience with a causal account of how the brain executes reliable computations.

## INTRODUCTION

To understand neural computation, neuroscientists need theoretical frameworks that explain how large populations of noisy, heterogeneous neurons can collectively and reliably implement a wide range of complex computations^1–3^. Unlike in human-engineered circuits, in biological circuits the individual neurons often exhibit continual variability in their tuning properties^4–6^, a phenomenon termed ‘representational drift’. Although this drift in single-cell coding appears to be widespread across many brain regions and animal species^7^, the general statistics of neural ensemble coding and the resulting ability to decode behaviorally relevant variables often remain stable over timescales longer than those characterizing the drift in cellular coding^4,8–10^. This dichotomy between neural ensemble-level stability and cellular variability raises the fundamental question of how neural circuits can maintain stable computations in the face of ongoing representational drift.

One potential solution to this question focuses on the computations performed via the low-dimensional (or latent) population variables encoded within high-dimensional neural dynamics. In analyses of large-scale neural recordings, identification of such low-dimensional latent variables typically involves a dimensional reduction of the set of neural activity traces^3,11,12^. Research to date on low-dimensional population codes can be broadly divided into studies that employed one of two successive generations of dimensionality reduction strategies. The first-generation approaches applied static dimensionality reduction methods; the second-generation techniques modeled the dynamics of large-scale neural populations within low-dimensional subspaces.

Specifically, the first-generation methods used static reduction techniques, such as those based on a principal or tensor component analysis^13^, to extract a low-dimensional set of descriptors of population activity^1,13–22^. Such methods (also termed manifold learning^23^), successfully identified low-dimensional subspaces of neural ensemble activity encoding behaviorally relevant variables^13,14^. Studies of these subspaces revealed key geometric properties of neural activity patterns^24,25^, such as that they are often constrained to evolve within specific manifolds^26^ (*e.g*., toroidal manifolds^27^). However, while these methods have provided important advances in our understanding of neural computation from a geometrical perspective, they generally do not describe neural dynamics from a mechanistic perspective or provide biologically interpretable causal explanations.

To address these shortcomings, second-generation techniques for analyzing large-scale neural dynamics explicitly inferred low-dimensional dynamical variables directly from neural recordings. Notable examples of such methods include linear dynamical systems^28,29^, recurrent switching linear dynamical systems^30–32^ (rSLDS), and low-rank recurrent neural networks^11,33–36^. These approaches yielded key advances by providing causal interpretations and identifying specific dynamical structures, *e.g*., line attractors^31,37^, but often relied on restrictive modeling assumptions and specific architectures^11,19,33–36^ that usually do not fully capture the rich geometry of biological computation^26^. Consequently, these dynamical methods also do not fit well with several empirical observations regarding the redundancy of neural coding^8^, the diversity and nonlinearity of single-cell tuning^38^, the high-dimensionality of neural activity manifolds^39,40^, and the robustness of biological neural ensemble coding to drift in cellular tuning properties^7,41^ (**Table S1**).

Recognizing the above limitations, to achieve a causal description of neural computation we propose a third-generation approach that unifies the respective geometric and dynamical perspectives of the first- and second-generation methods and is centered around a biological unit of neural computation termed the ‘Latent Processing Unit’ (LPU). In our framework, LPUs formalize the relationship between low-dimensional latent dynamics and high-dimensional neural ensemble activity while relying on reasonable, minimal assumptions about neuronal physiology. As in earlier studies of artificial neural networks^42,43^, our work on LPUs assumes that at individual synapses the pre- and post-synaptic activity levels are linearly related. Further, our framework uses smooth, nonlinear transfer functions relating a neuron’s synaptic current inputs to its somatic voltage. The framework allows for the possibility of multiple synaptic compartments, such as those that can underlie nonlinear dendritic computations. Beyond these basic ingredients, we make no assumptions about network architecture, implying that our theoretical results hold for a very broad class of biologically plausible networks. For this broad class, we prove the existence of LPUs and extend existing theories of neural computation by linking latent variable and neural activity dynamics through a set of dynamical equations and six mathematical theorems (**Methods**; **Supplementary Notes 1** and **2)**. Strikingly, as detailed below, our treatment yields predictions and explanations for multiple biological observations that were not well explained by prior frameworks^11,20,30–35,44^. Each of the six theorems, explained via illustrative examples in **Figures 1–6**, respectively, has substantial biological implications and allows the LPU framework to fill a conceptual gap in theoretical systems neuroscience:

**Figure 1.**
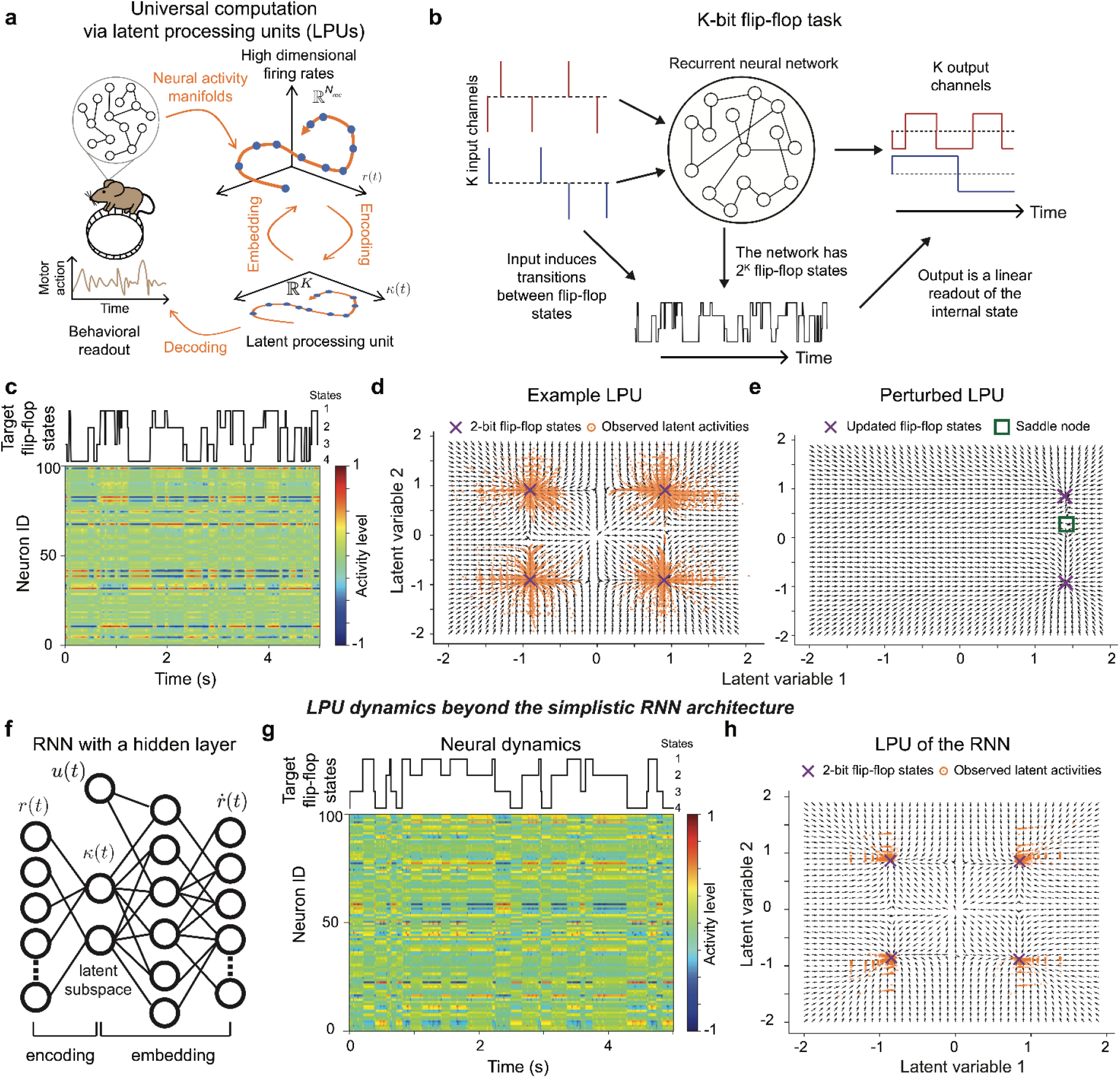
(Theorem 1, Universal computing). Latent processing units (LPUs) are low-dimensional dynamical systems that have universal computing capabilities. **(a)** Schematic overview of latent processing units (LPUs), showing the relationships between animal behavior, a high-dimensional set of neural activity traces, and neural ensemble coding, which often has much lower dimensionality. An LPU is a mathematical construct that describes neural computation and involves a set of latent variables, **κ**(*t*) ∈ ℝ^*K*^, and deterministic equations defining their time evolution. The latent variables are not directly measurable in neural recordings but instead are encoded in and can be inferred by taking weighted averages of individual cells’ activity levels, ***r***(*t*) ∈ ℝ^*N*^. We assume *K* ≪ *N*, such that the latent dynamics, 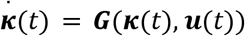, are termed low-dimensional. Then, the dynamics relevant for neural computation are parametrized solely by network parameters describing neural connectivity and input-output functions. **Theorem 1** about the universal computing capabilities of LPUs establishes that, with suitable synaptic weights, the latent dynamics can approximate any smooth and continuous trajectory as *N* → ∞. Cells’ activity traces constitute nonlinear dynamical embeddings of the latent variables, allowing neural trajectories to lie on extrinsically high-dimensional manifolds. Yet, parameters describing animal behavior can be inferred via linear decoding of the latent variables. Given this conceptual abstraction, **Theorem 1** guarantees that any set of smooth, behaviorally relevant variables can be realized in an LPU. **(b)** As an illustrative example of how computations enacted via high-dimensional (*N*) neural activity traces can be interpreted in terms of their LPU dynamics, we simulated recurrent neural networks (RNNs) that perform a *K*-bit flip-flop task, a well-studied short-term memory task^60^, in which the network transitions between 2^*K*^ different dynamical states. The network has *K* independent ternary inputs (0, –1 or 1) and *K* continuously valued outputs. Input values are mainly zeros but include pulses of ± 1 that signal transitions between different memory states. Output values convey the RNN’s dynamical state, and in each output channel the value should approximate that of the most recent pulse in the corresponding input channel. A well-trained RNN for this task should have 2^*K*^ attractive fixed-points that maintain internal states corresponding to the 2^*K*^ different flip-flop configurations, for which a *K*-dimensional LPU is needed (one latent variable per flip-flop; see **c–e**). That the fixed-points are attractive ensures the RNN can store these configurations long after the corresponding input pulses have ended. The output from the latent state is computed as a linear combination of neural activities. Here, the latent variable obeys the equation, 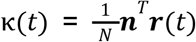, where the vector ***n*** is an encoding weight, illustrating that latent variables themselves can serve as outputs (**d, e**). **(c–e)** To illustrate that computations that seem high-dimensional in multineuronal recordings can often be explained using low-dimensional, identifiable LPUs, we examined an example RNN with 100 neurons, whose activity patterns were governed by a two-dimensional dynamical system. For this example, we trained a rank-2 RNN on a 2-bit flip-flop task using a two-step learning paradigm^33^. First, we trained the unconstrained, full-rank RNN using backpropagation through time (BPTT^99^), updating network parameters to minimize the mean squared error between the RNN readouts and target outputs. Second, we trained a rank-2 RNN, again using BPTT, to reproduce the dynamics of the full-rank RNN. This two-step training process captured the function of the unconstrained, full-rank network using a low-rank RNN, which then exhibited a readily interpretable two-dimensional LPU. **(c)** Raster plot showing the time-dependent activity of 100 neurons in an example RNN performing the 2-bit flip-flop task. The solid black line shows the target states the RNN should instantiate, as signaled by the transient input pulses illustrated in **b.** Driven by these pulses, the well-trained RNN transitions between its 4 different attractor states. Although many individual neurons exhibit state-dependent activity patterns, cellular tuning patterns are insufficient by themselves to explain the RNN dynamics. The individual activity patterns in the raster plot appear complex, but owing to **Theorem 1** and the low-dimensionality of learned fixed-point attractors, their collective embedding must enact an underlying low-dimensional dynamical system, the LPU, executing the computation. **(d, e)** Two-dimensional phase portraits of the latent dynamics that determine the tuning properties of the neurons in **c**. Panel **d** shows a phase portrait of the latent dynamics in the absence of external input to the RNN. The 4 attractive fixed-points are marked with purple **x**’s and correspond to the 4 different flip-flop states, which we refer to with a numeric pair, (*S1, S2*), in which each of *S1* and *S2* can have values of ± 1. Each orange dot marks an individual state that is traversed by the RNN’s dynamical trajectories across the 50s interval (beyond the 5s shown in **c**), during which the RNN performs the 2-bit flip-flop task. Black arrows denote flow vectors, all of unity length, that show the directions of the system’s time-dependent evolution in the phase portrait, as governed by the latent dynamics, 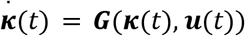. For the phase portrait of **e**, the RNN receives a constant input in its first channel with a magnitude of 10% of that of the binary pulses received during normal operation. This external perturbation leads to a bifurcation, such that two of the four original flip-flop states, (−1,−1) and (−1,1), no longer exist. Here, purple **x**’s mark the two remaining fixed-points, (1,1) and (1,-1), and the green square marks the saddle node between them. As shown by the flow vectors, the network will be attracted to either the (1,-1) or (1,1) states, as specified by the most recent value received in the RNN’s second input channel. Overall, the dynamics of the 100-neuron RNN can be well understood by analyzing the two-dimensional LPU (**d, e**), whereas a comparable interpretation is obscured within the set of neural activity traces (**c**). **(f)** Our formulation of RNNs as a combination of encoding-embedding maps enables the identification of latent variables in a broad class of architectures. In panels (**c-e**), we studied a basic RNN whose embedding map consisted of a generalized linear model. This panel shows a schematic of a slightly more complicated network with an LPU, which we refer to as a gain-modulated RNN. In this network class, the embedding map is modeled by a hidden layer with a tanh nonlinearity and the encoding map is once again linear. (**g, h**) For this example, we trained a rank-2 gain-modulated RNN on a 2-bit flip-flop task (100 neurons). We used an explicit rank constraint on the synaptic connectivity matrix, whereas the amplitude matrix was unconstrained (**Methods**). The plots in these panels mirror those in (**c,d**), which were computed using the neural and latent dynamics of the RNN with a hidden layer from panel (**f**).

### (1) Universal computing

A central challenge has been to explain how biological networks, composed of cells with properties constrained by biophysics, flexibly implement a vast range of computations. **Theorem 1** offers a resolution to this puzzle by showing that when neurons are connected within a network with LPUs, the LPUs can universally approximate any arbitrary dynamical system with the same dimensionality as the number of latent variables. Here, each latent variable is a linear combination of the activity levels of many neurons, rather than that of a single cell. This powerful mathematical result about the universality of computation shows how the brain could enact a broad array of computations and that, moreover, to perform a novel computation, the brain need not develop or evolve new types of neurons with distinctive biophysical or physiological properties. Instead, our treatment offers a theoretical guarantee that the neural dynamics needed to perform novel computations can be enacted by establishing new patterns of neural connectivity.

### (2) Computing over extended timescales

Neural spikes generally have millisecond-scale durations. Yet, cognition and animal behavior typically unfold over seconds to minutes, leading to the conundrum of how biological neural networks bridge these two sets of phenomena, of which the characteristic timescales differ by up to four orders-of-magnitude. **Theorem 2** resolves this question by showing that LPUs can sustain computations over durations far longer than the timescales of individual neural spikes, provided that the neural networks are sufficiently large. This result explains how neural populations collectively subserve long-lasting cognitive operations, despite the millisecond-scale time constants of single neurons. Owing to the resulting long timescales characterizing neural computation, neural manifolds^26^ naturally emerge, since the long-timescale dynamics are constrained to evolve within specific dynamical subspaces.

### (3) Optimality of linear LPU readouts

Another longstanding challenge has been to understand how brain areas that receive high-dimensional, noisy signals, *e.g*., those sent from upstream brain regions, faithfully extract task-relevant information from these unreliable signal sets. Although important ideas regarding linear decoding and communication subspaces have been previously considered^8,38,45,46^, **Theorem 3** provides a formal proof that linear readouts can serve as the optimal output of computations done by LPUs, guaranteeing the simple and robust extraction of information. This implies that biologically simple mechanisms, such as weighted summation, can serve as efficient readouts of population activity.

### (4) Intrinsic low-dimensional dynamics with extrinsic high-dimensional embeddings

A recent, significant claim in systems neuroscience has been that the high-dimensional nature of neural population activity may be a key signature of high-dimensional neural population coding^39,40^. **Theorem 4** challenges this claim by proving that high-dimensional neural activity can be extrinsic, *i.e*., it is not a property of a high-dimensional, population-level computation but instead arises from the nonlinear transformations that occur within individual neurons. In this view, the intrinsic dimensionality of computation can remain low and organized around the latent manifolds, even when the dimensionality of neural population activity is much higher (see also Ref. 47). This relationship, between high-dimensional neural activity and a low-dimensional computation enacted by these dynamics, shows that neural activity can seem highly complex^39,40^ yet still support compact, low-dimensional (and potentially simple) computations that drive animal behavior.

### (5) Neural coding without causal influence on behavior

Across several decades of past research, neuroscientists have often focused on neurons with activity tuned to specific stimuli or behavioral conditions. Well-known examples include hippocampal place cells^48^ and orientation-selective neurons^49^ in primary visual cortex. Although it has often been assumed that the activity levels of such neurons are causally related to animal perception, cognition, or behavior, recent causal experiments suggest that perturbations to the dynamics of stimulus-coding neurons can greatly alter the activity levels of individual or subsets of cells without major changes in neural population activity or animal behavior^50–52^. **Theorem 5** reconciles these distinct perspectives by proving that LPUs preserve their low-dimensional latent dynamics even when large subsets of neurons (of which the activity levels may be correlated with animal behavior) are perturbed. The key insight is that the dynamical subspaces driving computation are causally distinct from the parameters that define single-cell tuning, making the former resilient to fluctuations in single cell activity.

### (6) Computational robustness to representational drift

Synaptic connections in the brain undergo constant modification due to plasticity, spine turnover, or consolidation processes, raising the question of how computations can remain stable and reliable despite continual change. Empirical studies suggest that population codes are often robust but not fully invariant to representational drift^7,53^. **Theorem 6** provides insight into this phenomenon by proving that the nonlinearity of the embedding that grants LPUs their capacity to serve as universal approximators also establishes the existence of drift-robust (but not invariant) dimensions that enable continual adaptation of readouts without loss of computational power. This result provides a principled framework for theoretical studies of representational drift in biological networks and shows that robustness is not incidental but emerges systematically at large scales from the structural properties of LPUs.

Each of the above theoretical results yields experimentally testable predictions. Key perturbations influencing behavior should occur in low-dimensional linear subspaces corresponding to the latent dynamics (**Theorems 1–3**). Neural manifolds, which arise naturally in our framework from the task-driven slow timescales of latent dynamics, should exhibit extrinsic curvature (**Theorem 4**). Finally, neural populations should maintain coding subspaces that are robust to both random and structured forms of representational drift (**Theorems 5, 6**). Future empirical tests of these predictions will help refine models of neural computation and lead to an integrated understanding of low-dimensional computation, neural activity manifolds, and robustness to drift and perturbations.

Overall, LPUs provide a principled, biologically inspired, and causal framework to examine how reliable computations can underlie complex forms of animal behavior. While latent dynamics^11,31–34,44^ and linear communication subspaces^8,45^ between brain regions are regularly studied in the field, our treatment identifies a unified network mechanism to causally store, manipulate, and communicate information across brain regions. Further, by directly linking latent variables to the geometry of neural dynamics, we provide a mechanistic account of how neural manifolds and associated constraints could arise from latent variables, unifying two lines of research that were previously largely disjoint. These strides derive from the conceptual advance of viewing latent computations through an encoding-embedding relationship that constrains neural dynamics via two independent maps, the embedding and encoding maps. This differs crucially from AI-inspired, deep learning models of latent variables, in which the two maps are designed to invert one another^14,17^, highlighting the potential richness of computation in biological neural networks.

## RESULTS

Our goal in this section is to develop a theoretical account of neural computation that explains how reliable, low-dimensional computations are embedded within high-dimensional neural activity. Starting from four basic principles, we link neural dynamics to population-level latent variables and introduce our central object of study, the LPU. We characterize its properties through a set of six theorems that collectively address the biological puzzles raised above.

### Latent processing units: From constrained elements to universal computation

In a population-coding framework^54^, neurons constitute physical substrates that encode a set of abstract coding variables:

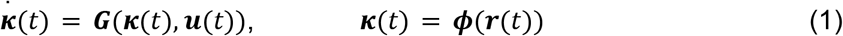

where ***r***(*t*) ∈ ℝ^*N*^ represents the activities of *N* neurons at time *t*, ***κ***(*t*) ∈ ℝ^*K*^ denotes the *latent* (*i.e*., directly unobservable) coding variables with *K* ≪ *N*, 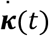 refers to the time derivative, ***G***(·) denotes the latent vector field, 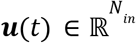 represents task-relevant inputs (*e.g*., contextual cues), and ***ϕ***: ℝ^*N*^ → ℝ^*K*^ is an *encoding* function transforming neural activities into latent variables. Existing methods for modeling latent variables differ primarily in how they define the encoding map and its inverse, ***κ***(*t*) → ***r***(*t*), in which the latter often is subject to additional assumptions (*e.g*., linearity^33^), largely due to analytical or computational convenience^3,11,20,33,34^. As a result, this often leads to overly simplified neural activity traces with one-to-one mappings between ***r***(*t*) and ***κ***(*t*) values, creating a disjunction between the dynamical^54^ and geometrical^26^ views of neural computation.

The key methodological advance of our work is to supplement Eq. (1) with an *embedding* map, which does not perform an inversion (***κ***(*t*) → ***r***(*t*)) but instead maps the latent variables to the neural dynamics 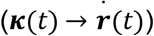. This conceptual change allows us to introduce latent dynamical systems with theoretically and biologically desirable properties across a broad class of neural networks. We rely on four basic principles that make minimal assumptions about the nature of neural computation. The first three principles govern the mathematical properties of the latent variables, whereas the last one connects these abstract coding variables to biological reality.

The first two principles, **P1** and **P2**, connect latent variables to neural activity. The first principle defines latent variables as linear combinations of neural activity traces, whereas the second one connects the neural dynamics to the latent variables:

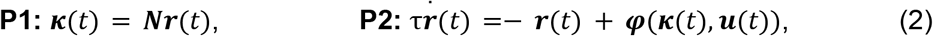

where ***N*** ∈ ℝ^*K*×*N*^ is referred to as the encoding matrix, ***φ***(·) is a dynamical (nonlinear) embedding map, and τ > 0 is the neuronal time constant. This construction enforces a linear encoding map, which may seem computationally restrictive at first blush but in fact is not (see **Theorem 1** below) and has several desirable properties (**Table S2**). Notably, Eq. (2) can be used to capture commonly studied network architectures in computational neuroscience^11,14,20,30–35,37,44^ (see **Supplementary Note 1** for details). This difference between the linearity of encoding and the nonlinearity of embedding is also at the core of compressed sensing theory^55^, which has previously been applied to study sparse neural population codes^56,57^.

The third principle (**P3**) posits that behaviorally relevant computations are driven by a low-dimensional latent dynamical system, mathematically described by a vector field ***G*** as in Eq. (1). Conceptually, this principle states that the latent (population-coding) variables, and not the noisy activities of individual neurons, process information and drive the animal’s behavior. Below, we show that these three abstract principles **P1–P3** can explain diverse empirical aspects of the biological neural computation (see **Fig. 1a** for a visual summary).

To connect these abstract principles to biology, we note that neural networks communicate through synapses and induce presynaptic currents at each node (*e.g*., neurons), and that distinct architectures differ in how they turn these presynaptic currents into neural activity traces ***r***(*t*). To remain agnostic to these architecture-specific processes, we consider an abstract neuron with presynaptic inputs that are described by the equation ***z***(*t*) = ***Wr***(*t*) + ***W***_*in*_ ***u***(*t*) + ***b*** ∈ ℝ^*N*^ but do not specify how these are transformed at individual neurons. Here, ***W*** ∈ ℝ^*N*×*N*^ are synaptic weights, 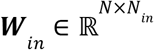 are input weights, and ***b*** ∈ ℝ^*N*^ are neuronal biases. Thus, in a broad, biologically plausible class of networks, each neuron can have multiple presynaptic inputs:

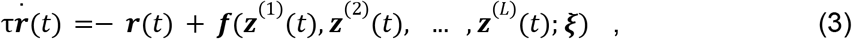

where 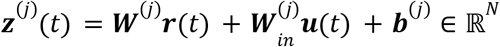 is the net induced current in the *j*th synaptic compartment. The term synaptic compartment reflects the biophysical observation that distinct branches of a neuron’s dendritic arbor can independently integrate synaptic inputs, with their currents combining nonlinearly at the soma. This process is modeled here through distinct sets of parameters 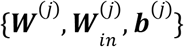 across *j* = 1, …, *L* compartments. We use ***ξ*** (of unspecified dimension) to denote any additional parameters not directly affecting the presynaptic inputs ***z***^(*j*)^ (*t*). ***f***: = {*f*_1_ (.·), …, *f*_*N*_ (·)} is a collection of element-wise nonlinearities with *f*_*i*_ : ℝ^*L*^ → ℝ. To specify an architecture, one must define the nonlinearities ***f***, the parameters ***ξ***, the number of synaptic compartments, and how the post-synaptic currents combine within each neuron. For the remainder of this work, unless otherwise stated, we drop the superscript (*j*) for notational convenience.

To introduce the final principle, we leverage the empirical observation that most biological networks effectively exhibit low-rank structure^58^ and write the synaptic weights as ***W*** ≈ ***MN***, where ***N*** ∈ ℝ^*K*×*N*^ is the encoding matrix defined above, and we refer to ***M*** ∈ ℝ^*N*×*K*^ as the embedding matrix. Then, the final principle (**P4**) supplies the encoding matrix of **P1** from biology, so that ***MNr***(*t*) = ***Mκ***(*t*), and thereby establishes how latent variables shape the presynaptic inputs:

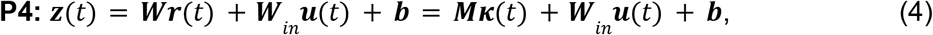

which is a map from a low-dimensional (*N*_*in*_ + *K*) subspace to a high-dimensional (*N*) one. The encoding matrix (***N***) defines the latent variables (via **P1**) and the embedding matrix (***M***) transforms the latent variables into arguments of ***φ***(·) (cf. **P2**). With this definition, Eq. (3) becomes a biologically constrained version of Eq. (2), where all abstract parameters of the latent dynamical system (*i.e*., ***N, φ***(·), and behavioral decoders; see **Fig. 1a**) are defined by the physical parameters of the network (**P4**), such as synaptic connections between neurons.

To formalize latent computations in such networks, we first note that the latent dynamics of networks defined by Eq. (2) do not depend on ***r***(*t*) explicitly, only implicitly via ***κ***(*t*) = ***Nr***(*t*):

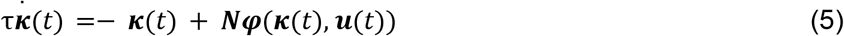

owing to the linear encoding principle (**P1**). This group includes those described by Eq. (3). We refer to the latent dynamical system in Eq. (5) as an LPU if it also satisfies a universal approximation property. In the following theorem, we show that one realistic class of networks, which makes use of a finite repertoire of cell types, achieves this property.

#### Theorem 1

*(Universal approximation theorem). Let a neural network follow the time dynamics in Eq. (3) and fix the latent dimension K. Suppose each element-wise nonlinearity f*_*i*_ (·) *is sampled independently from a finite set of functions. Assume this distribution assigns positive probability to at least one function that is locally bounded, piecewise continuous, and not equal almost everywhere to a polynomial in one of its arguments. Then, with probability one as N* → ∞, *the induced latent dynamics can approximate any continuous K-dimensional vector field* ***G***(***κ***(*t*), ***u***(*t*)) *on compact subsets of* (***κ, u***)*-space with arbitrarily small uniform error*.

This theorem (proof in **Methods**) extends classical universal approximation results for recurrent neural networks^35,42^ to a biologically relevant setting with heterogeneous element-wise nonlinearities (representing distinct cell types) and dendritic computations beyond the standard linear-nonlinear form. Moreover, the probabilistic formulation is in line with biological neural networks having distinct cell types, in which the universal approximation property can be achieved even if some neurons have linear or other simple (polynomial) transfer functions. Finally, as a consequence of **Theorem 1**, when Eq. (5) describes the dynamics of an LPU, the second term of the equation, ***Nφ***(***κ***(*t*), ***u***(*t*)), is promoted implicitly to a universal approximator in the limit *N* → ∞.

### An illustration of computation through LPUs in distinct architectures

Earlier work studying low-rank RNNs has shown that computations carried out by high-dimensional neural dynamics can often be described via low-dimensional latent dynamical systems^11,33–36^. Here, we extend this line of work and illustrate how LPUs carry out neural computations in diverse RNN architectures. To this end, we focus on two key architectures whose dynamics can be written as:

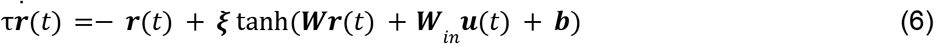

with ***ξ*** ∈ ℝ^*N*×*N*^. When ***ξ*** is an identity matrix, Eq. (6) corresponds to a basic RNN in which each rate-coding neuron applies its nonlinearity locally to its own presynaptic input. Thus, the computation is a linear summation of inputs followed by an element-wise nonlinearity. For the general case, Eq. (6) describes a gain-modulated RNN, in which the gain parameters ***ξ*** mix the nonlinear responses, tanh(***z***(*t*)), across all neurons. This architecture captures nonlinear, dendrite-like computations at individual neurons by defining the embedding map as a feedforward neural network with a single hidden layer (**Methods**). As we show in **Methods**, both architectures are members of the class described by Eq. (3) and satisfy the requirements of **Theorem 1**, *i.e*., they both support LPUs. However, only gain-modulated RNNs are capable of universally modeling ***φ*** (**Methods**). Hence, while basic RNNs have commonly been used as models of neural computation^59,60^, especially with low-rank constraints^11,33–35^, they impose specific restrictions on the embedding map that gain-modulated RNNs do not. This distinction motivates the analyses below, in which we use the LPU framework to study one instance of preserved latent computations beyond the basic RNN architectures considered in prior work^11,33–35^.

We first trained full-rank basic RNNs to perform K-bit flip-flop tasks (**Figs. 1** and **S1**). These tasks require trained RNNs to maintain 2^*K*^ distinct internal states representing possible K-bit combinations (*e.g*., ({-1,-1}, {-1,1}, {1,-1}, {11}) for *K* = 2) and transition between them in response to transient inputs (**Fig. 1b**). For an illustrative case with 2 bits, a well-trained RNN on this task developed 2^2^ = 4 attractive fixed-points in the neural dynamics (**Fig. 1c-e**). The network’s computational mechanism could be well approximated by a rank-2 RNN after a rank distillation process^33^ (**Fig. 1b**). The distilled RNN exhibited state-dependent persistent activity at the single-neuron level (**Fig. 1c**) and formed four attractive fixed-points in its two-dimensional LPU (**Fig. 1d**). Transient inputs caused the latent state to jump between attractors, updating the network’s behavioral output.

LPUs enable analyses of neural computation beyond the identification of equilibrium states, as has been commonly done before^60^. They give direct access to latent trajectories outside the point attractors (**Fig. 1d**) and make it possible to quantify how latent dynamics transform under external inputs (**Fig. 1e**) or synaptic perturbations. For example, introducing an input to the first channel induced a bifurcation in the flow field (**Fig. 1e**), resulting in the disappearance of two of the four fixed-point attractors. The flow along each output direction remained largely independent. Input to one channel perturbed only its associated latent variable, while the other continued to evolve according to its own dynamics (**Fig. 1e**). Overall, training the high-dimensional RNN parameters effectively led to the identification of a low-dimensional solution^61^, in which each flip-flop channel operates through a dedicated latent dimension.

Next, we trained a gain-modulated RNN on the same 2-bit flip-flop task (**Fig. 1f**). To showcase an alternative training paradigm, we enforced a rank constraint (*K* = 2) throughout training. Similar to the RNN in **Fig. 1c-e**, neural dynamics in this architecture also showed preferential tuning towards flip-flop states (**Fig. 1g**) and produced qualitatively the same phase portrait in its LPU (**Fig. 1h**). Overall, two very different architectures (basic and gain-modulated RNNs) learned the same LPU with four point attractors despite their distinct single-neuron computations and expressive power. This commonality supports the notion that learned solutions reflect the demands, structure, and dimensionality of the task^62^, which can transcend the specific network architectures and be extracted via low-dimensional LPUs.

### Latent computations across extended timescales are enabled by universal approximation

How can neural networks stabilize and sustain latent computations over timescales much longer than those of individual neurons^63^? To answer this question, we first illustrate how latent variables can perform computations through their dynamical evolution with seconds-long effective timescales.

To build intuition, consider the ideal case of a network with infinitely many neurons (*N* → ∞) and no inputs (***u*** = 0). For an LPU with universal dynamics, Eq. (5) can be written as:

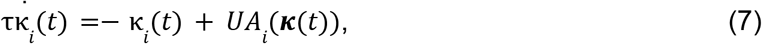

where *UA*_*i*_ (***κ***(*t*)) refers to a universal approximator governing the dynamics of the *i*th latent variable. To see how latent computations can be extended beyond the neuronal time constant τ ~ *O*(*ms*), consider a universal approximator, 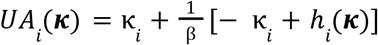. Here, β = τ_*eff*_ /τ is the timescale separation between the neural (τ) and behavioral (τ) timescales and *h*_*i*_ (***κ***) is a continuous vector field subserving the target computation. For large β, the universal approximator is fine-tuned around the identity (*UA*_*i*_ (***κ***) ≈ *κ*_*i*_), and the small deviations are precisely what encode the target computation enabled by an arbitrarily selected β-independent ***h***(***κ***). (This process is similar to the one in linear networks, in which the eigenvalues of the Jacobian matrix deviate only slightly from the edge of stability to enable diverse network timescales^64^.) After fine-tuning of the universal approximator ***UA***(***κ***), the network dynamics follow an effective equation:

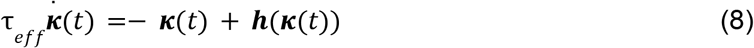

Since the right hand side of the dynamics does not depend on the ratio β ≫ 1, rescaling time *t* → *t*/τ_*eff*_ renders the dynamics β-independent. ***h***(***κ***) defines the computation across an independently picked τ_*eff*_ timescale regardless of τ. This is what we refer to as the separation of neural and latent timescales. In the limit *N* → ∞, as long as *h*_*i*_(***κ***(*t*)) is continuous and β finite, **Theorem 1** guarantees that Eq. (8) can be achieved. Next, we consider the effects of having a finite *N*.

### Separation of neural and latent timescales requires large-scale networks

In this section, we study the limits imposed by finite-size errors on how well networks can implement a target ***h***(***κ***). We show that inherent variability of network parameters (*e.g*., day-to-day changes in neural tuning properties) limits how long latent computations driven by a particular ***h***(***κ***) can be sustained, but this limit can be mitigated by an increased network size *N*.

As discussed above, **Theorem 1** guarantees the existence of an approximator ***UA***(***κ***; *N*) = ***Nφ***(***κ***(*t*), ***u***(*t*)) that becomes universal (***UA***(***κ***)) in the limit *N* → ∞. However, there is no prescribed procedure for choosing ***N*** and ***φ***(·) to achieve this. To make theoretical progress in the large *N* limit, we make two assumptions (see **Methods** for formal statements). First, we assume that ***UA***(***κ***; *N*) is self-averaging^65^, *i.e*., for large *N*, the latent dynamics depend on the distribution of synaptic parameters, but not on the exact pattern of sampled synapses. Second, we assume that the error between ***UA***(***κ***; *N*) and ***UA***(***κ***) is dominated by the inherent variability of the synapses (*e.g*., synaptic failures) and therefore is independent of the specific computation or timescale separation β. Then, we can prove the following theorem:

#### Theorem 2

*(Computation over extended timescales). Let an LPU implement the function in Eq. (8) for a finite N using an approximator* ***UA***(***κ***; *N*) *that is self-averaging. Further, let the synaptic parameters have inherent variability that is independent of* β. *Then, the estimator* ***h***(***κ***; *N*) = β(***UA***(***κ***; *N*) − ***κ***) + ***κ*** *has an error scaling as* 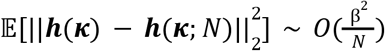, *where the expectation is taken over the synaptic parameter distribution for a given* ***κ***.

This theorem implies that if 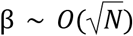, the latent vector field ***h***(***κ***) cannot be reliably implemented. This observation offers an alternative explanation for why biological neural networks may engage millions of neurons to solve tasks that can be performed with hundreds of artificial neurons^66^. One classical explanation is that a large number of neurons is required to suppress noise in downstream readouts^67^. **Theorem 2** introduces a complementary explanation: inherent variability of synaptic parameters (*e.g*., due to ongoing plasticity^9,53,68^) requires increased network size to accommodate seconds-long effective timescales. As a simple scaling estimate, roughly a million neurons are needed to extend the effective timescale from *O*(*ms*) to *O*(*s*).

### Designing bistable dynamical systems with distinct neural and latent timescales

We now illustrate the consequences of **Theorem 2** with bistable dynamical systems capable of storing flip-flop states. Such systems are commonly studied as models of binary decision-making^54,60^, and they provide a concrete, biologically relevant case in which **Theorem 2**’s finite-size error scaling can be observed directly.

To study bistable dynamics, one could attempt to train RNNs performing K-bit flip-flop tasks and study the emergent dynamics with 2^*K*^ attractors, as we have done in **Fig. 1**. However, training networks with a large number of neurons becomes prohibitively difficult (**Fig. S1a-b**). Instead, we study this problem through a sampling-based approach (**Fig. S1c, Supplementary Note 2**; also see Refs.34,35). To design a one-dimensional LPU with bistable dynamics, we draw the encoding and embedding weights of each neuron from a specifically designed probability distribution (**Fig. 2a**; **Methods**). The distribution is designed such that the LPU realizes a target latent timescale (τ_*eff*_, see **Fig. 2b**) for its linearized latent dynamics around the origin, which it does despite fixed neural time constants τ = 1 *ms* in the limit *N* → ∞. For finite *N*, synaptic variability can introduce errors (**Fig. 2b**). As we show next, **Theorem 2** quantitatively predicts these errors.

**Figure 2.**
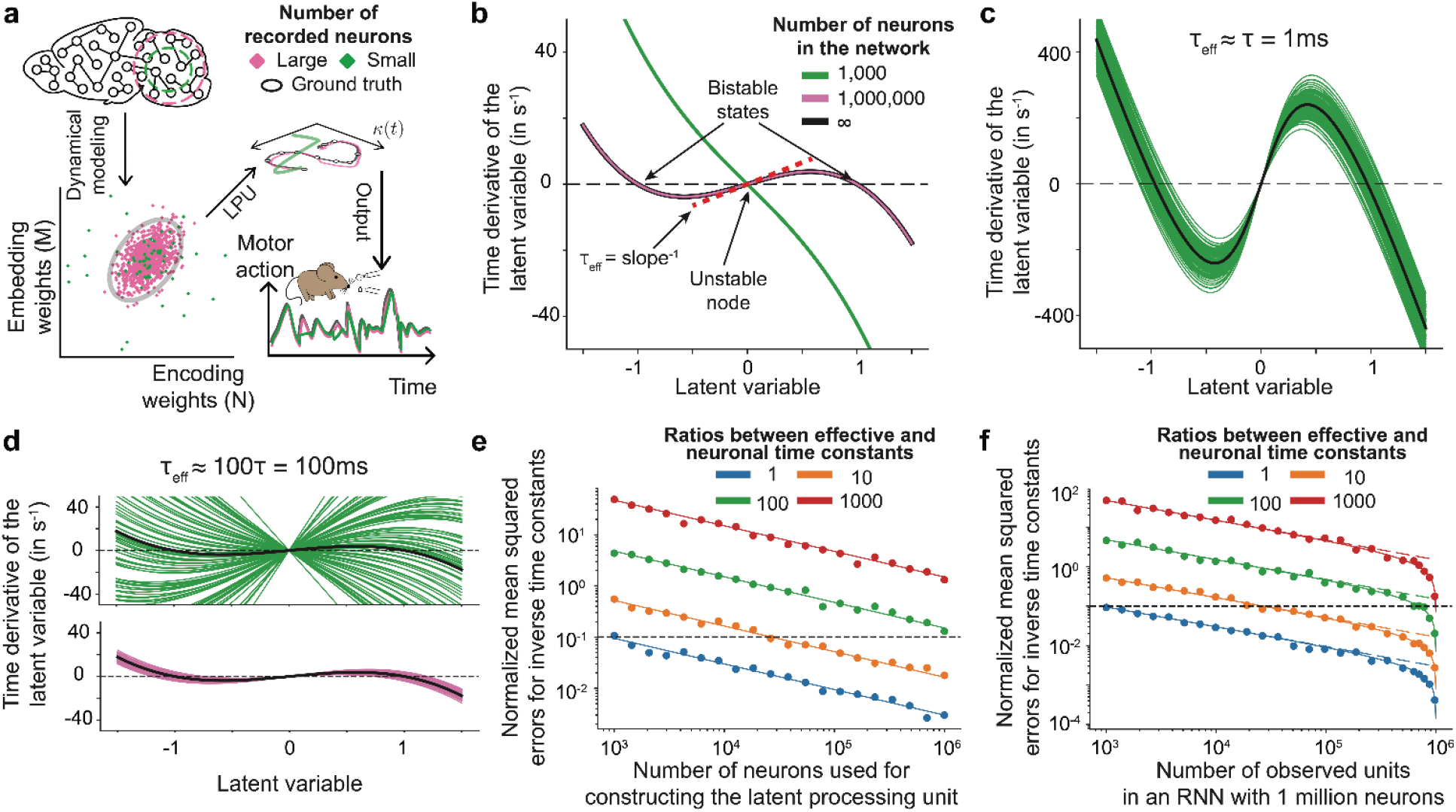
(Theorem 2, Computing over extended timescales). Embedding LPUs in large networks allows reliable computations over timescales far slower than cellular time constants. Using LPUs, neural populations can perform computations over timescales much longer than those governing the dynamics of individual neurons (**Theorem 2**). LPU dynamics are constructed by learning sets of encoding and embedding weights, as instantiated in neural synapses, suited to perform a specific computation of interest. The total set of synaptic connections, rather than any specific synaptic weight, determines the computation that the network performs. This perspective, in turn, suggests that systems neuroscientists should focus on latent variables as the core elements of network computation. But, computations at extended timescales come at the cost of scale requirements: Large numbers of neurons are needed to support such latent computations, and, conversely, large neural recordings may be necessary to extract the dynamical properties of these networks. **(a)** Schematic illustrating the dual role of scale and subsampling in LPUs. Embedding LPUs in large networks stabilizes latent computations against biological noise and extends their processing capabilities to durations far longer than the fast timescales characteristic of individual neurons (**Theorem 2**). In contrast, a much smaller set of cells suffices for accurate decoding of behaviorally relevant variables. In an example case, recordings sample only a small portion of the cells participating in the relevant computation (*top left*). Recordings from small (green) or large (pink) subpopulations of cells permit approximations of the underlying weight distribution with varying levels of accuracy (*bottom left*, the ellipse outlines the 95% C.I. of the distribution of embedding and encoding weights, within which 95% of the neural weights reside). Notably, the weight distribution defines the latent dynamical system enacted by the full neural population (*top right*); accurate estimation of these dynamics requires recordings from large sets of individual cells (pink). On the other hand, low-dimensional projections of neural dynamics, *e.g*., the subsequent behavioral readouts (*bottom right*), may be decoded from small subsets of cells (green). **(b–d)** To illustrate how large networks are needed to support latent computations over timescales of seconds, we examined RNNs designed to perform a discrimination task. Using our theoretical framework, we designed a sampling process for setting the encoding and embedding weights and thus the synaptic connections between neurons (see **Supplementary Note 2**), such that an RNN with infinitely many cells contains a one-dimensional LPU with two attractive fixed-points at *κ* ≈± 1 and an unstable node at *κ* = 0. Plots in **b–d** show the time derivative of the latent variable (*y*-axis) as a function of the latent variable itself (*x*-axis). The black solid lines were obtained by sampling the encoding and embedding weights from a predetermined, two-dimensional Gaussian distribution that yields the desired LPU as *N* → ∞ (**Supplementary Note 2**). While the theoretical design is exact in the limit *N* → ∞, we created RNNs with finite numbers of cells that approximate this limit, albeit with variability due to sampling noise that diminishes as *N* → ∞. Each colored solid line shows a latent dynamical system belonging to a network with either 1,000 (green curves) or 1,000,000 (purple curves) neurons. In these RNNs, the individual cells had membrane time constants of 1 ms, but the latent systems were designed such that their latent variables, when initialized near *κ* ≈ 0, could evolve towards one of the two attractors over intervals lasting up to seconds. **(b)** Each RNN’s response to the discrimination task was determined according to which one of the two attractors (*κ* ≈± 1) its latent variable approached. This allows an evaluation of how well the RNNs constructed with a finite number of neurons capture the desired LPU properties. For instance, here, sampling a network with 1,000 neurons led to a monostable LPU, *i.e*., a qualitative error in realizing the targeted bistable dynamics. In contrast, a network with 1,000,000 neurons had minimal deviations from the target LPU. To quantify the error incurred during network construction in panels **e** and **f** below, we consider the associated effective time constant obtained by linearizing the bistable latent dynamical system around the origin (red dashed line). The inverse of the effective time constant τ_*eff*_ is given by the slope of the line 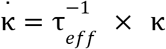 passing through the origin. **(c, d)** Accuracy of reconstruction depends on both the scale of the network and the separation of the network’s effective timescale from that of the individual cells. **(c)** When the latent bistable dynamical system was designed to have an effective timescale τ ≈ τ_*eff*_ = 1*ms*, all networks with as few as 1,000 cells accurately reconstructed the qualitative bistable dynamics. In **c** and **d**, each colored line characterizes the LPU dynamics of such a constructed network, whereas black solid lines show the analytically designed LPU dynamics. Each of the 100 different green lines shows the LPU dynamics of an individual sampled RNN with 1,000 cells. **(d)** For τ_*eff*_ ≈ 100τ = 100 *ms*, networks with 1,000 cells had substantial variability in their LPU dynamics (*top*), and far more poorly approximated the dynamics of the large network limit than networks with 1,000,000 cells (*bottom*). The black solid line shows the analytically designed LPU dynamics. Each of the 100 different green lines corresponds to networks with 1,000 cells, whereas each of the 100 pink lines is for networks with 1,000,000 cells. **(e, f)** We quantified how finite-size effects induce errors in the estimated effective time constants for LPUs implementing bistable latent dynamics (**Supplementary Note 2**). In a first scenario **(e)**, the theoretical disagreement between the designed and the empirically sampled LPU diminishes as the number of cells *N* used to embed the LPU increases (**Supplementary Note 2**). In a second scenario **(f)**, when a finite network realizes a particular LPU, LPU reconstruction with subsampled cells from this network grows more accurate as *N*_*obs*_ → *N*_*obs*_, where *N* is the number of observed neurons. **(e)** Theoretical analysis suggests that if *N* neurons are used to construct an LPU with a target effective time constant, the errors between the target and sampled (inverse) time constants 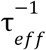 decline as 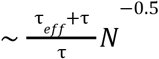. Plots show the normalized root mean squared errors between theoretically designed 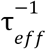 (limit *N* → ∞) and those calculated from the sampled networks, normalized by the theoretical value. Each data point corresponds to an error value computed from 30 sampled networks; solid lines denote theoretically predicted errors; the horizontal dashed line corresponds to 10% estimation errors. Lines of different colors show results from networks with different τ_*eff*_ values. The agreement between the theory and simulations suggests that, *e.g*., to be able to design a bistable dynamical system with τ_*eff*_ ten times larger than τ, one needs a hundred times more neurons to achieve comparable errors for producing the same qualitative dynamics as a network with τ_*eff*_ = τ. Overall, this panel illustrates the necessity of using large networks to construct LPUs that support reliable computations over long timescales. **(f)** In this panel, we study an experimentally relevant question: how to faithfully reconstruct the dynamical properties of a sampled network using partially observed neurons. If LPU dynamics are reconstructed using partially recorded neurons from a network, the errors in estimated effective time constants decline as 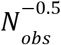 for *N*_*obs*_ ≪ *N*, but rapidly decline as 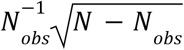 once *N*_*obs*_ ~ *N*. To test this result, we first designed LPUs with bistable dynamics and varying effective time constants using the analytical limit *N* → ∞ (black lines in earlier panels). Then, we sampled RNNs with a million neurons each from the corresponding weight distributions. Finally, using randomly subsampled neurons from each of these constructed networks, we estimated the corresponding LPU, specifically, estimated the 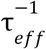 values. Plots show the errors between 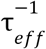 values of sampled (ground truth) and those of reconstructed (from partial observations) networks normalized by the analytical target in the limit *N* → ∞. Each data point corresponds to an error value computed from 30 sampled networks; solid lines denote theoretically predicted errors; dotted lines correspond to the asymptotic 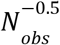 scaling when *N*_*obs*_ ≪ *N*; and horizontal dashed line corresponds to 10% estimation errors. Note that up until the large-scale recording regime, *i.e*., *N*_*obs*_ ~ *N*, the benefit of adding more neurons for the accurate reconstruction is minimal. Hence, to accurately estimate the effective timescales of an actual network implementing the bistable dynamics while remaining faithful to the underlying weight distributions, most neurons need to be observed and used for the reconstruction. Overall, while neural populations can perform computations over timescales much longer than those governing the dynamics of individual cells (**Theorem 2**), this comes at the expense of requiring large numbers of cells in the network implementing the LPU dynamics. This in turn motivates large-scale recordings to correctly estimate both the dynamical (*e.g*., τ_*eff*_) and structural (*e.g*., encoding and embedding weights) properties of the underlying network.

As anticipated, we found that LPUs with effective timescales close to the intrinsic timescales of the individual neurons could be reliably constructed with as few as a thousand neurons (**Fig. 2c**), in line with prior work showing that small networks are sufficient for solving common behavioral tasks^34,66^. However, the same number of neurons failed to support LPUs operating at seconds-long timescales (**Fig. 2d**, *top*). When β ≈ 100, the failures could even be qualitative, with thousands of neurons spuriously generating monostable dynamics (**Fig. 2b** and **2d**, *top*). In contrast, one million neurons generated bistable dynamics reliably (**Fig. 2d**, *bottom*). In **Fig. 2e**, we further confirmed the quantitative agreement between numerical simulations and the theoretical predictions of **Theorem 2**.

**Theorem 2** characterizes the finite-size error incurred when an LPU is designed to implement a target computation using all neurons in the network. A complementary question arises when recording the whole population is not an option, *e.g*., as is typical in recordings from mammalian brains. Can the LPU of a neural network be reconstructed using partial observations, *i.e*., without having recorded every neuron? A thorough analytical treatment of this question is challenging^69–71^. Instead, in **Supplementary Note 2**, we perform exact analytical calculations on the bistable dynamical system using an idealized estimation procedure. As illustrated in **Fig. 2f**, we find that errors governing the reconstruction of a particular LPU scale as 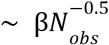 when observing only a fraction of neurons (*N*_*obs*_ ≪ *N*). In contrast, in the limit *N*_*obs*_ → *N*, errors decline much faster 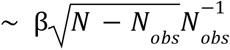. Together with prior findings that a few thousand neurons can suffice for optimal linear decoding of population codes^72,73^, this result highlights the advantage of large-scale recordings for inferring latent neural dynamics.

### Identifiable linear readouts can subserve complex behavior

A standard concern with latent variable models is identifiability, *i.e*., whether the latent variables are unique given the observed neural activities. The state of the art is to extract latent variables that are identifiable up to linear transformations^14^. LPUs possess this property by design (**Supplementary Note 1**), going a step further to remove the need for identification altogether. To see how, consider a practical question. To achieve optimal behavioral readouts ***Ψ***(***κ***(*t*)), which linear combinations of ***κ***(*t*), if any, would need to be tracked within a biological network? If the linear information (*i.e*., what downstream neurons can access through direct synaptic projections^74^) is sufficient to support behavior, the network does not need to track the latent variables at all. Instead, LPU computations can be read out directly from ***r***(*t*), bypassing the need to explicitly infer the latent variables that drive the internal computation (**Theorem 3** below).

While linear readouts have long been viewed as biologically plausible and efficient ^8,38,45,75^, whether they are sufficient and optimal has remained unresolved. This claim (that linear readouts suffice) might appear counterintuitive, since nonlinear decoders are more expressive by design. In our framework, the trainable nonlinear embedding governs computational complexity (**P2, Theorem 1**), whereas the linear encoding (**P1**) ensures identifiability:

#### Theorem 3

*(Optimality of linear LPU readouts). Let* ***r***(*t*) ∈ ℝ^*N*^ *represent neural activities of an LPU. Let the latent variables* ***κ*** ∈ ℝ^*K*^ *evolve following a continuous vector field* ***G***(·). *Then, for a fixed K* ≪ *N, any continuously differentiable function of the latent variables can be read out linearly from* ***r***(*t*) *with vanishing approximation errors in the limit N* → ∞.

**Theorem 3** implies that with sufficient neurons (*N* ≫ *K*), biological networks can use linear readouts to approximate nonlinear ones with vanishing approximation errors. This makes linear readouts ideal communication tools, as restricting to them incurs no loss in expressive power.

### Testing the optimality of linear readouts

Using a few hundred neurons, earlier work has shown that categorical variables can be optimally decoded from cortical neural activities using linear decoders^38,46^. Here, we extend this optimality result to a large dataset of thousands of neurons. Specifically, we re-analyzed published mesoscopic recordings of layer 2/3 pyramidal neurons across up to eight neocortical regions while mice performed a visual discrimination task^8^ (**Figs. 3, S2**). Mice saw either a Go or NoGo stimulus, followed by a brief delay, then indicated their decision via licking or withholding a lick (**Fig. 3a**). During this task, the recordings captured on average 3595 ± 989 neurons while mice performed 1296 ± 379 correct trials within 5 ± 1 imaging sessions (all quantities in mean ± s.d.; **Fig. 3b**). Using the extracted neural activities, we trained linear and nonlinear decoders to differentiate between correctly and incorrectly performed trials. Although decoding does not imply causal involvement, the correct trial identity correlates with both cue identity and response. Hence, this information is likely to be actively communicated across regions^8^, which makes trial identity a suitable test case for the optimality of linear decoders under the assumptions of **Theorem 3**.

**Figure 3.**
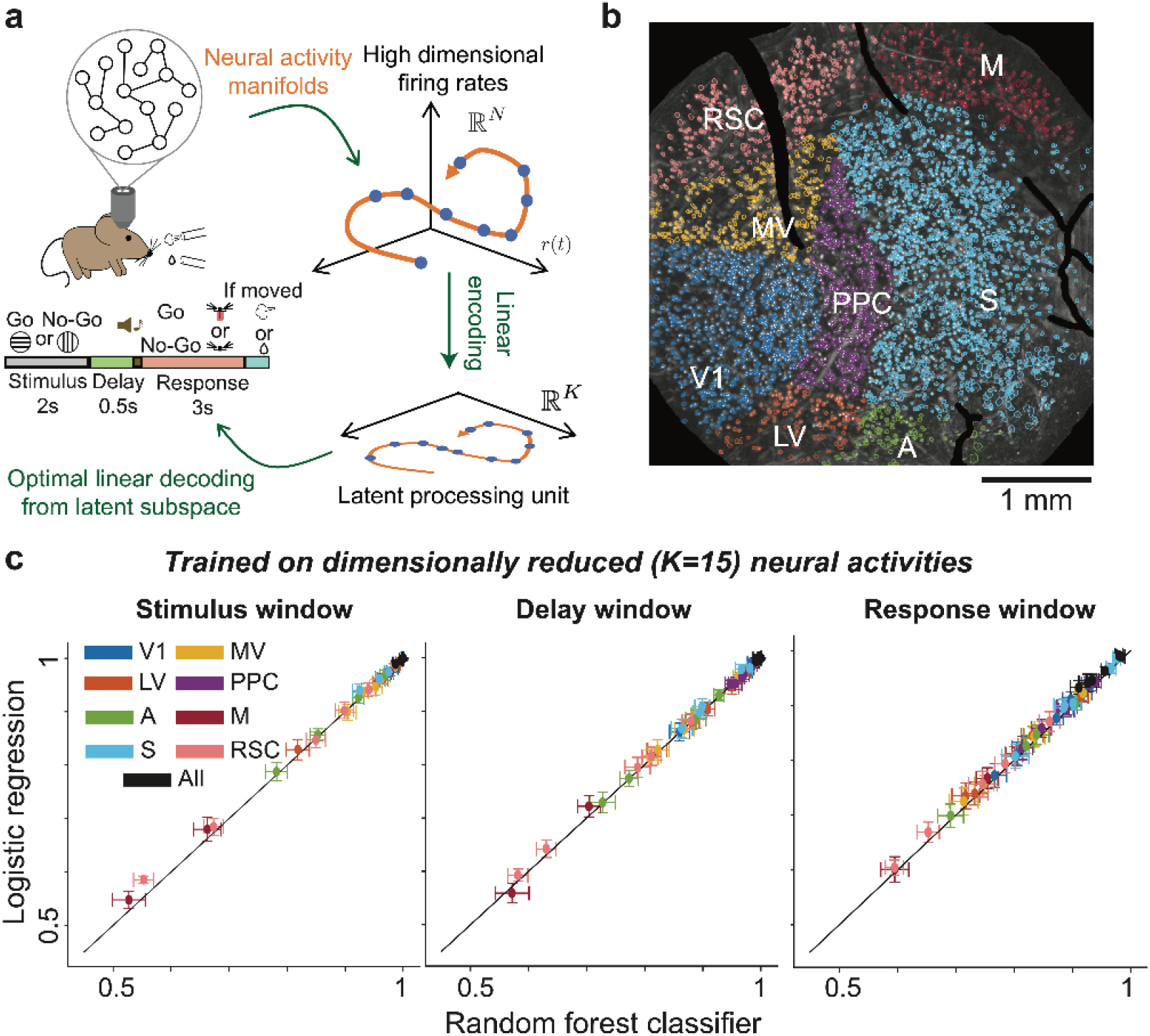
(Theorem 3, Optimality of linear LPU readouts). Linear decoders of behaviorally relevant, latent dynamical variables attain performances comparable to nonlinear decoders. **(a)** In principle, the linear encoding of behaviorally relevant variables by LPUs allows neuroscientists to decode these variables of interest directly from sets of neural activity traces, without explicit identification of the underlying latent variables and dynamics. Notably, although neuroscientists commonly constrain decoders of neural population activity to linear forms, in the LPU framework this linearity does not diminish their expressivity when decoding is constrained to latent subspaces, as formalized in our universal decoding theorem (**Theorem 3**). To showcase this theorem, here we study and compare the accuracies of linear and nonlinear decoders trained on presumed latent subspaces. Specifically, we re-analyzed recent recordings of the concurrent dynamics of thousands of neurons, sampled across the entire visual cortex, plus surrounding neocortical regions, in mice performing a visual discrimination task^8^. In this binary visual task with Go-NoGo structure, mice were trained to lick a spout to receive a reward after the presentation of one visual stimulus (horizontal gratings), but to avoid licking after the presentation of another stimulus (vertical gratings). Each trial had a 2s period of stimulus presentation, followed by a 0.5s delay period and a 3s response period during which the mouse had to make its response. **(b)** Example maximum projection of a Ca^2+^ video of the mouse cortex, from which we extracted 5,292 neurons. Overlaid are spatial profiles of the individual extracted neurons, color-coded by cortical region. Brain areas: V1 (primary visual cortex), LV (lateral visual cortex), MV (medial visual cortex), PPC (posterior parietal cortex), A (auditory cortex), S (somatosensory cortex), M (motor cortex), and RSC (retrosplenial cortex). **(c)** If behavioral signals are communicated between brain regions using latent variables, a linear combination of latent variables can allow optimal information transfer (**Theorem 3**). However, the LPU framework does not preclude the possibility that nonlinear decoders of high-dimensional neural ensemble activity might yield superior decoding performance, as is sometimes observed in practice^15^. For instance, this superiority of nonlinear decoders can occur when there is redundant information in the high-dimensional set of traces that is unavailable in the low-dimensional set of latent variables. Hence, one concrete prediction of our LPU framework is that when decoders are trained on latent variables, linear and nonlinear decoders should perform equally well. To test these ideas, for the experiment of **(a)** we trained linear and nonlinear decoders on the set of extracted latent variables to classify the visual stimulus based on the neural dynamics evoked within individual or all 8 brain areas. The activity traces were from trials on which the mouse performed the task correctly and were averaged over one of three different intervals within the trial structure (stimulus period, [0.5 s, 2 s]; delay period, [2 s, 2.5 s]; response period, [2.5 s, 5.5 s]). Given the presumption that latent variables are low-dimensional and linearly encoded in the neural activity traces, we first performed a linear dimensionality reduction of the set of neural activity traces using a PLS analysis. Using these low-dimensional variables, we trained logistic regression (linear) and random forest (nonlinear) decoders of the trial-type. The accuracies of both decoder types saturated when the presumed latent subspace had as few as 15 dimensions (**Fig. S2)**, in line with prior work using linear decoders^8^. Across brain regions, the accuracies of linear decoders matched those of nonlinear decoders trained on these dimensionally reduced latent variables. Each datum denotes the mean test accuracy for cortical neurons from an individual brain area in one animal (error bars: s.d. across 100 different randomly chosen training and testing splits of the data, which are from 6 mice and a total of 30 imaging sessions). Solid lines: diagonals passing through the origin with unity slope. Overall, **Theorem 3** concerns the ability of linear decoders to optimally extract information encoded in the low-dimensional set of latent variables, as illustrated in **c.** LPUs provide a principled explanation for the effectiveness and efficiency of low-dimensional codes using linear communication subspaces, as found in multiple past experimental studies of animal brains^8,45^.

We compared the performance of linear (logistic regression, LDA) and nonlinear (random forest, QDA) decoders trained on the activity traces from individual brain regions (**Figs. 3c** and **S2**). All decoders were regularized using dimensionality bottlenecks via partial least-squares, following the procedures outlined in Ref. 8 and summarized in **Methods**. Across regions and task epochs (stimulus, delay, response, post-response), decoder performance increased most dramatically within the first 15 latent dimensions (**Fig. S2**), in which linear decoders either matched or slightly outperformed nonlinear ones (**Fig. 3c**). As dimensionality increased, performance converged across decoder types (**Fig. S2**). These results align with earlier findings in the primary visual cortex, in which class covariances across cues were nearly equal, making linear and quadratic discriminants equally effective^72^. Here, we generalize these observations to thousands of cells sampled across the cortex. For behaviorally relevant information that must be transmitted across regions, linear subspaces are both sufficient and biologically plausible, consistent with **Theorem 3**.

### Emergence of neural manifolds from the separation of timescales

We now illustrate how low-dimensional latent variables can subserve high-dimensional neural activity. As we formalize in **Theorem 4** below, the mathematics underlying this also naturally give rise to what are commonly studied as neural manifolds in systems neuroscience^1,3,26^.

For LPUs to have complex, nonlinear temporal dynamics, at least one of the encoding or embedding maps must be nonlinear (**Supplementary Note 1**; **Theorem 1**). Linear encoding is needed for the autonomous dynamics of latent variables without imposing restrictive conditions (**Table S2**) and enables optimal linear readouts directly from neural activity (**Theorem 3**). This observation leaves the nonlinearity to the embedding (**Fig. 4a**), giving rise to neural manifolds:

**Figure 4.**
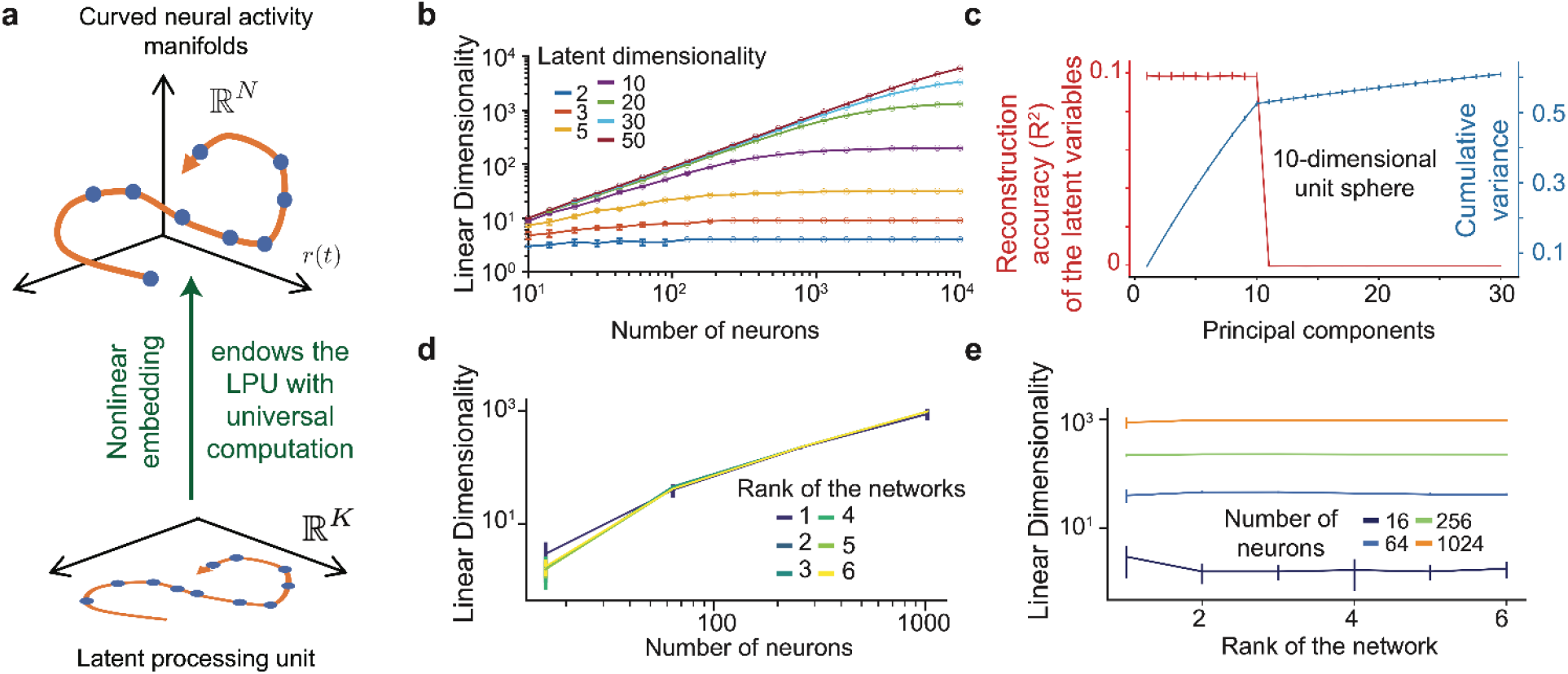
(Theorem 4, Intrinsic low-dimensional dynamics with extrinsic high-dimensional embeddings). Nonlinear embedding of LPUs with low-dimensional dynamics can lead to extrinsic high-dimensional representations. **(a)** Given linear encoding maps, which endow LPUs with their interpretability and desirable properties (*e.g*., the potential optimality of linear decoding; **Figure 3**), the property of universality positing that LPUs can perform complex computations implies the embedding maps must be nonlinear (**Theorem 1**). The diagram illustrates that the embedding map takes the low-dimensional latent variables and maps them to a set of neural activity traces with a high linear dimensionality. (Here, the linear dimensionality can be quantified as the number of principal components needed to explain 99% of the cumulative variance in the neural ensemble dynamics.) Using these definitions and a nonlinear mapping as shown in the plot, one can construct a plausible scenario, in which even one-dimensional LPUs can lead to a linear dimensionality that scales linearly with the number of cells in the network. This illustrates a key prediction of our framework about the dimensionality of coding. Namely, the set of neural activity traces can have a high linear dimensionality that originates from nonlinear coding manifolds but that can be spanned by a low-dimensional set of latent dynamical variables (**Theorem**). **(b–e)** While **Theorem 4** proves that the linear dimensionality of the set of neural activity traces can scale linearly with the number of cells in the network even when the latent dimensionality is fixed, the nonlinear embedding illustrated in panel **a** can achieve this in several distinct ways. In these panels, we illustrate two empirically important ways this can be plausibly achieved (see **Figure S5** for another relevant example with randomly connected LPUs). **(b)** As a starting point, to illustrate that nonlinear embedding maps can increase linear dimensionality even when there are no dynamics associated with the latent variables, we performed a static and noiseless embedding of an LPU, in which the latent variables are sampled uniformly from the surface of a *K*-dimensional unit sphere, with neural activities given by 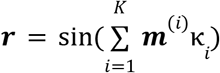. Here, κ_*i*_ is the *i*th latent variable describing the unit sphere, ***m***^(*i*)^ ∈ ℝ^*N*^ denotes its embedding weights onto *N* neurons, and sin(·) is applied elementwise. The plot, with logarithmic axis scales, shows the linear dimensionality of the neural activities (*y*-axis) as a function of the number of cells (*x*-axis). Increasing the number of cells leads to an orders-of-magnitude rise in the linear dimensionality of their activities, requiring up to thousands of principal components to explain 99% of the variance in neural activities even for *K* ~ *O*(10). Data are mean ± s.e.m. values over 5 random initializations. Solid lines: linear interpolations between data points. **(c)** The simulations in panel **b** do far more than reveal theoretical points of interest. Rather, they provide an alternative plausible explanation for a recent experimental study^40^ arguing that about half the variance in neural ensemble dynamics is in principal dimensions that encode behaviorally irrelevant aspects of neural computation in the context of a fixed experimental task. Our illustration shows that these dimensions do not necessarily correspond to new coding variables, rather they can be the signatures of curvature caused by the nonlinear embedding of latent variables subserving behavior (**Theorem 4**). Similar to **(b)**, we embedded a 10-dimensional latent sphere onto a network with 1,000 neurons. We computed the correlations between the top principal components of the neural dynamics and the latent variables, as well as the cumulative variance in these components. Here, the embedding of a 10-dimensional unit sphere results in about half the variance being in neural activity dimensions that are uncorrelated with latent variables, even though no additional coding variables exist. Data are mean ± s.e.m. values over 100 random initializations. Solid lines: linear interpolations between data points. **(d, e)** To show that latent variables can lead to a spuriously high linear dimensionality in task-performing noisy RNNs, and that the behavioral task, not the rank of the network, is expected to determine the intrinsic dimensionality of the LPU, we studied the dimensionality of neural dynamics in RNNs trained to perform a sequence sorting task. In this task, the networks are given a set of *T* numeric values, which the RNNs must sequentially output in order of ascending magnitude. RNNs performing this task (here for *T* = 2, see **Fig. S4** for longer sequences) had increased linear dimensionality with added neurons **(d)** despite having ranks of 1–6. Notably, the rise in the linear dimensionality cannot be explained by the rise in network rank **(e).** Data are mean ± s.e.m. values across 10 runs. Solid lines: linear interpolations between data points.

#### Theorem 4

*(Neural manifolds). Let* ***r***(*t*) ∈ ℝ^*N*^ *represent neural activities of an LPU, which evolve via* 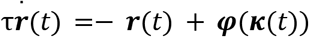 *with* ∥***φ***(***κ***) − ***φ***(***κ***′)∥_2_ ≤ *L*∥***κ*** − ***κ***′∥_2_ *for some L* > 0 *and* ***φ***(***κ***) *is bounded. If the latent variables evolve slowly relative to neural dynamics and* β = τ_*eff*_ /τ ≫ 1, *neural activities lie on average near the intrinsically low-dimensional structure* ***r***(***κ***) = ***φ***(***κ***) + *O*(β^−1^) *whose extrinsic dimensionality can attain the upper bound N*.

**Theorem 4** implies that neural manifolds^3,26^ naturally emerge from the separation of latent and neural timescales, whereas high linear dimensionality in large-scale recordings^39,40^ can originate from low-dimensional LPUs with nonlinear embeddings. Specifically, as long as ***φ*** satisfies a mild regularity condition (satisfied by common nonlinearities such as tanh or clipped ReLU; and expected as neural firing rates are intrinsically bounded by biological constraints), neural activities lie near a low-dimensional structure parameterized by ***κ***. Whether this structure constitutes a formal mathematical manifold depends on further properties of ***φ*** (**Methods**), though neuroscientists broadly refer to such structures as neural manifolds^3,26^. Furthermore, noisy dynamics in real datasets (*e.g*., trial-to-trial variability in neural activities, stochasticity in ***φ*** due to neuronal noise, or short-term drift in neural tuning properties) mean that this confinement should be considered an average tendency, rather than an exact constraint on the neural activities.

While neural manifolds can technically be defined in the dynamical systems view of neural computation^11,33,34^, constrained embeddings in these models (**Supplementary Note 1**) confine activity to linear subspaces whose dimensionality is at most *K*. **Theorem 4** overcomes this limitation by lifting these architectural assumptions, in which even a one-dimensional LPU can have an extrinsic dimensionality that is equal to the neuron count (**Methods**).

### Nonlinear embedding can lead to extrinsically high-dimensional dynamics

A curved embedding, ***r***(***κ***) = ***φ***(***κ***), can increase the extrinsic dimensionality. Below, we show three distinct mechanisms through which this occurs.

As a first experiment, we performed a static noiseless embedding of a *K*-dimensional unit (latent) sphere following the formula 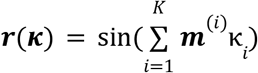, where ***m***^(*i*)^ ∈ ℝ^*N*^ are embedding weights drawn from a normal distribution with zero mean (**Methods**). This can be considered the limit β → ∞, in which activities instantaneously decay onto the neural manifold. We quantified extrinsic dimensionality via linear dimensionality, *i.e*., the minimum number of principal components needed to explain 99% of the variance in neural activities. With as few as 50 latent dimensions, the linear dimensionality grew linearly with neuron count up to 10,000 neurons (**Fig. 4b**). This occurred even with 10 latent dimensions with a different nonlinearity (**Fig. S3a**). Overall, the non-zero embedding curvature induced by the nonlinearity was sufficient to increase the extrinsic dimensionality by three orders of magnitude.

This setup also provides an alternative explanation for a recent observation in Ref. 40, in which about half the variance in neural activities remains uncorrelated with behavior, whereas the top 10-20 principal components (PCs) correlate with the animal’s behavior. While this observation could reflect the high dimensionality of latent variables, some of which are simply not correlated with behavior, the same scenario can be reproduced with the embedding of a 10-dimensional latent sphere (**Fig. 4c**). Here, only the first 10 PCs had non-negligible correlations with the latent variables (*i.e*., the proxy for behaviorally relevant directions), whereas about half of the variance was in PCs that were not correlated with the latent sphere (**Fig. 4c**). We confirmed that this result was not an artifact of non-smoothness of the manifold, as the configurations used in this experiment preserved the distances in the embedded unit sphere (**Fig. S3b**). Behaviorally irrelevant principal dimensions can thus arise without high-dimensional latent variables, and may instead reflect geometric consequences of nonlinear embedding (*i.e*., curved neural manifolds) rather than behaviorally irrelevant internal signals of neural computation^40^.

In the second experiment, we trained noisy low-rank RNNs on a sequence sorting task, in which the network is presented with a set of numbers and must output them in sorted order (**Fig. 4d-e**). This task is low-dimensional and temporally well-structured, and it starts from zero-initialized activity. Thus, any extrinsic dimensionality that emerges must stem from the interaction of the learned LPU dynamics with injected noise. Consistent with the first experiment, linear dimensionality scaled with neuron count (**Fig. 4d**) but remained constant across RNN rank (**Fig. 4e**) for fixed *N*. These patterns held across varying sequence lengths (**Fig. S4**). Consistent with prior research on task dimensionality^62,76^, this suggests that the inherent LPUs, rather than the full rank permitted by the architecture, drive the scaling.

In the third experiment, we went beyond task-trained LPUs. Instead, we studied the extrinsic dimensionality of randomly initialized rank-deficient RNNs, which can become chaotic^77^ in the limit *K, N* → ∞. In these networks, linear dimensionality grew in proportion to the number of neurons in the RNNs despite fixed rank (**Fig. S5**). Hence, high extrinsic dimensionality is not necessarily a task-specific phenomenon but could be a generic property of nonlinear LPUs initialized with random connections.

Together, these three experiments (static nonlinear embedding, interplay of structured LPUs with noise, and randomly initialized chaotic LPUs) provide converging evidence consistent with **Theorem 4**. Across settings differing in task structure, noise, and LPU architecture, nonlinear embedding maps produced high extrinsic dimensionality despite fixed latent dimensionality.

### Separation of causal and acausal coding dimensions

Now, we show that the independence of encoding (**P1**) and embedding (**P2**) maps enables LPUs to separate neural activity that causally drives the computation from neural activity that is merely correlated with the latent variables.

To formally introduce causality into our framework, we define two types of dimensions in population activity: (i) causal coding dimensions, spanned by the row subspace of the encoding matrix ***N***, and (ii) acausal coding dimensions, defined as the remaining neural activity directions that have non-zero overlap with the image of the embedding map ***φ***. The former originate from the linear encoding ***κ***(*t*) = ***Nr***(*t*), specifying which neural directions feed back into, and causally influence, the LPU dynamics. The latter are described by the low-dimensional neural manifold ***r***(***κ***) ≈ ***φ***(***κ***) (**Theorem 4**) and specify how latent variables are expressed in neural activity.

In the most general scenario, acausal coding dimensions exist for the networks defined in Eq. (3). In one analytically convenient case, a linearity constraint ***r***(*t*) = ***Aκ***(*t*) can be imposed for some ***A*** ∈ ℝ^*N*×*K*^. This simplifying assumption underlies many existing models, including rSLDS^20,30–32,37,44^, low-rank RNNs^11,33–35^, and the latest work on latent circuits^36^. In such models, the coding dimensions coincide with the span of the embedding map. However, this choice forces the richness of biological activity into an artificial affine model in which neural activities lie on affine subspaces^3,26^, in contrast to our discussion of manifold dimensionality above. As we now formalize, the separation between causal and acausal coding dimensions provides a natural explanation for representational redundancy in biological neural networks:

#### Theorem 5

*(Neural coding without causal influence on behavior). Let* ***r***(*t*) *denote the neural activity supporting an LPU. Changes in* ***r***(*t*) *orthogonal to the causal coding directions, even along those correlated with the latent variables, do not alter the LPU dynamics. Any difference in neural activity caused by such perturbations decays exponentially over time*.

This theorem shows that networks described by Eq. (2) can code for behavior without exerting causal influence. It also suggests one trivial case of redundancy in neural population codes. For instance, a neuron *i* may have zero weights in the *i*th column of the encoding matrix ***N***, but a non-zero φ_*i*_ (***κ***) value. Such a neuron would have no causal influence on the LPU but would be tuned to the latent variables. In biological neural networks, such acausally coding neurons (**Fig. 5a**) may play a communication role, relaying internal states to downstream regions.

**Figure 5.**
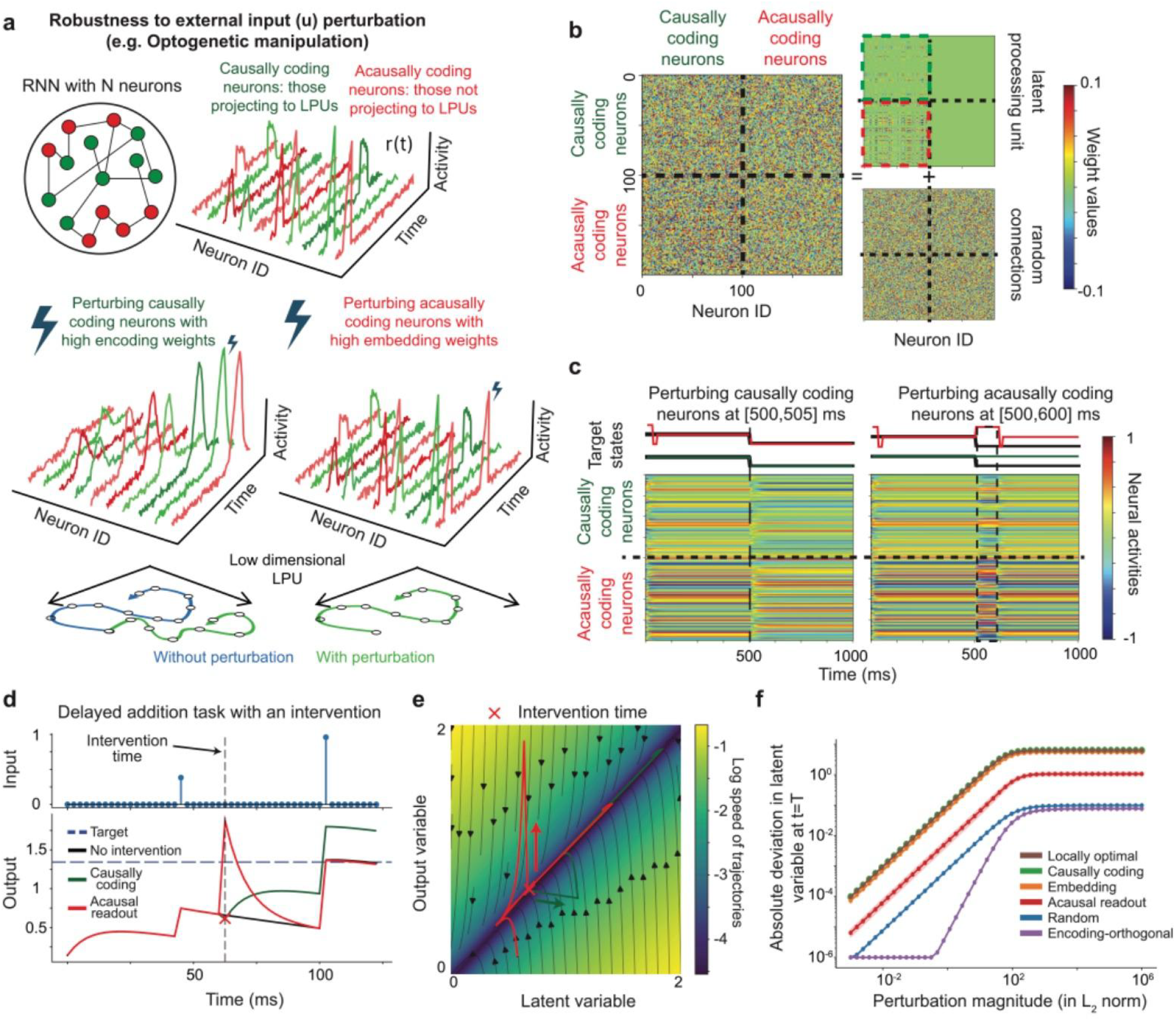
(Theorem 5, Neural coding without causal influence on behavior). Neurons with encoding weights allowing re-entrant influence on LPU dynamics causally shape neural computations, whereas cells lacking this property can merely code for behavioral variables. The LPU framework allows a clear distinction between the tuning properties of individual cells and their causal contributions to the neural computation (**Theorem 5**). While intuition suggests that synaptic connections in the network should subserve a causal link to neural computation, random excitatory and inhibitory connections can mutually cancel; instead, components of synaptic connections that re-enter through the linear encoding map, *i.e*., *re-entrant connections*, are the causal drivers of the latent computation (**Theorem 5**). To illustrate this distinction and how structured encoding weights, not necessarily the full set of synaptic connections, define the causal contribution of individual cells to LPU computation, we present here an *in silico* perturbation experiment. **(a)** A key advantage of our framework compared to earlier work is that the parameters defining the tuning properties of individual cells (*i.e*., embedding weights) are independent of the re-entrant connections through which neurons encode the LPU and causally shape its dynamics. To illustrate this, we designed a perturbation experiment with the RNN of **Fig. 1c–e.** For this network, we called cells ‘causally coding’ if they had non-zero re-entrant connections to at least one of the latent dynamical variables. We called cells ‘acausally coding’ if they had non-zero synaptic weights but lacked re-entrant causal influence on the latent variables. Crucially, causal perturbations applied to the dynamics of acausally coding cells should not affect network operations even though their activities are correlated with the latent variables (**Theorem 5**). **(b, c)** To illustrate that individual cells’ causal contributions to LPUs are defined by their encoding properties (**Theorem 5**), we performed a simulation using the example RNN from **Fig. 1c–e** with 100 neurons, which can transition between fixed-point attractors. We first augmented this RNN by adding 100 acausally coding neurons, with zero encoding but non-zero embedding weights. While the encoding weights of individual cells are not observable, the embedding weights can be inferred via the tuning curves. Using this approach, we designed a realistic perturbation experiment based on the observable tuning properties of the individual cells, in which we activated a group of acausally coding cells over a fixed time window. This perturbative activation was delivered to cells tuned to the specific flip-flop state into which we wished to induce a transition. For comparison, to highlight that activating causally coding cells *can* enable these transitions, we also performed a study in which we selectively activated groups of causally coding cells. **(b)** Plots showing the synaptic connections between neurons in an augmented network, in which we added 100 acausally coding cells that did not re-enter the LPU but still coded for the flip-flop states via non-zero embedding weights. *Left*: Matrix of synaptic weights belonging to the augmented RNN after addition of the acausally coding neurons. To compute this, we first expanded the connections between the original 100 causally coding neurons, whose connections are shown in the matrix top right with a green rectangle, and then padded the connectivity matrix with zeros to incorporate an additional 100 neurons. *Right*: Respecting the low-rank structure of the full matrix, we added random connections from the causally coding neurons to the acausally coding neurons (red rectangle) but not *vice versa*. This created an acausally coding group of neurons that received structured information from the LPU dynamics but did not causally affect these dynamics. To mask the resulting stereotypical structure and bands of zero connectivities, from which acausally coding neurons could be extracted trivially, we also introduced random projections between all neurons, with roughly twice the s.d. of the structured connections. In the end, the full connectivity matrix of the augmented network is the sum of the structured and random components with all-to-all connections in the full network. **(c)** Raster plots of neural activity from an *in silico* perturbation experiment. *Top*: Black lines show the desired flip-flop states, green lines represent states read out from causally coding neurons, and red lines show the states that are readout from acausally coding neurons. *Left*. Perturbing causally coding neurons led to the desired state change, decodable from both types of neurons. *Right*. Perturbing acausally coding neurons did not alter the flip-flop state, though the readout (incorrectly) changed for acausally coding neurons only for the duration of the perturbation. This experiment suggests that tuning properties or synaptic connections alone do not necessarily indicate a neuron’s causal contribution to the LPU (**Theorem 5**). Instead, to have causal influence, neurons must have a re-entrant influence on the LPU dynamics through the encoding weights. **(d-f)** To illustrate that **Theorem 5** provides an effective strategy for perturbing and controlling large networks in a regression scenario, we studied various types of interventions in rank-one RNNs trained to perform a delayed addition task, in which two scalar pulses presented at times *t* = *t*_1/2_ must be integrated and their sum must be reported as the output at the end of a trial. We trained 51 rank-one RNNs in total, each with 10,000 neurons, with each trial lasting 125 ms. **(d)** We designed perturbation experiments on the fully trained RNNs, in which we delivered perturbation pulses at the midway point of each trial (dashed line) along two directions: the encoding direction, which re-enters the LPU and constitutes a causal intervention, and the readout direction, which correlates with the latent variable but does not re-enter the LPU dynamics, constituting an acausal intervention (**Methods**). The plot shows an illustrative trial under both perturbation types. Perturbing along the encoding direction (green) permanently displaced the latent variable and drove the network to an incorrect output. In contrast, perturbing along the readout direction (red) only transiently shifted the output, but the network output (taken from the acausal readout) recovered to the correct target by the end of the trial. The perturbation strength is measured via the L2 norm of the perturbation pulse, which is one for both types in this example. **(e)** The phase portrait of the reduced dynamical system (consisting of the latent variable and the readout variable) provides a geometrical explanation for the asymmetric results in panel **d**. The flow field, colored by log-speed in the natural basis, reveals a slow manifold along the diagonal on which integration unfolds, flanked by fast transverse relaxation along the y-axis. Note the approximate nature of the line attractor, in which both latent and readout variables decay towards the origin in the absence of external inputs consistent with **d**. By **Theorem 5**, the latent variable is the causal driver of this slow integration, while the readout tracks the latent dynamics as an acausal coding direction. A perturbation along the readout direction (red line) displaces the trajectory in this reduced system off the manifold, but the strong transverse flow rapidly returns the trajectory to its original attractor. In contrast, a causal perturbation (green line) displaces the latent variable directly, shifting the trajectory across a basin boundary and committing the network to a new point in the manifold. Hence, the causal reach of an intervention is governed entirely by its projection onto the causally coding subspace, through which neurons have re-entrant influence on the LPU dynamics. The black lines with arrows denote the vector field that governs the dynamics of the two variable system, colored arrows denote the two types of perturbations. **(f)** The encoding direction becomes approximately aligned with the trial-averaged locally optimal perturbation direction, *i.e*., one that induces maximal local changes when the norm of the perturbation pulse is fixed (**Methods**), and can therefore be used to steer neural population activity. The plot shows the average absolute (solid lines) deviations in the latent variable at the end of the trials when networks are perturbed following procedures similar to panels **d-e**, including several other strategies (see legends, described below). Lines correspond to averages across 51 successfully trained rank-one RNNs (each with 10,000 neurons), with shaded regions indicating standard deviation. Intervening through the encoding (green) direction is approximately equivalent to intervening through the trial-averaged locally optimal (brown) perturbation direction across the full range of perturbation strengths tested, confirming that the causally coding subspace provides a practical and accessible target for neural control. For several orders of perturbation magnitude (10^−3^ −10^1^), the signed deviation in the latent variable scales linearly (log-log slope = 0.999, R^2^ = 0.999), indicating that precise linear control of the latent variable is achievable through the encoding direction. Beyond this linear range, nonlinear effects dominate and the deviation saturates. In contrast, intervening through the readout (red), encoding-orthogonal (purple), or purely random (blue) directions produces substantially smaller deviations. The embedding (orange) direction, which shares substantial overlap with the encoding direction (cosine similarity = 0.740 ± 0.003, mean ± s.d.), produces slightly lower deviations. Since all perturbation directions activate neurons embedded in an all-to-all connected network, these results show that causal influence over latent computation is not determined by connectivity alone. Instead, it depends on the component of neural activity that re-enters the LPU through the encoding map.

### Re-entrant weights, not the connectome, define the causal coding dimensions

**Theorem 5** provides a practical guideline for perturbing LPUs: target neural activity directions aligned with the causal coding dimensions, namely those spanned by the row subspace of ***N***. Here, we show that these dimensions are not necessarily directly visible in the connectome or other observable network properties.

A naive approach to estimating the causal coding directions would be to extract neural tuning relationships. As discussed above, single-cell analysis uncovers the effects of ***φ***. Hence, identifying neurons tuned to the latent variables (or behavior) is not sufficient. One might then ask whether synaptic connectivity, *e.g*., complete knowledge of the connectome, could resolve this ambiguity. Indeed, by **P4**, the encoding weights (***N***) must be realized through synaptic connections. As we show next, connectome information alone is also not sufficient. The relevant quantity is not the mere presence of connections between individual neurons, but the specific re-entrant projections hidden within the full pattern of synaptic weights.

To illustrate this point, we modified the basic RNN from **Fig. 1c-e** by adding 100 acausally coding neurons (no re-entrant influence, non-zero embedding; **Fig. 5b-c**). Without further assumptions or modifications, these neurons are functionally inert and structurally disconnected, and can be clearly distinguished when the synaptic connections are observed (**Fig. 5b**, *top right*). To mask this structure, we added a random weight matrix to the low-rank LPU connectivity, inducing dense, bidirectional synaptic connections between all neurons (**Fig. 5b**, *bottom right*). As a result, the structure in the connectome was masked (**Fig. 5b**, *left*). Despite the addition of a densely connected component, causally coding neurons continued to influence the latent states (**Fig. 5c**, *left*), whereas perturbing acausally coding ones had no lasting effects (**Fig. 5c**, *right*). While acausally coding neurons remained functionally ineffective because they had zero re-entrant contribution to the LPU, sufficiently strong random connectivity eventually dominated the recurrent dynamics and disrupted the LPU computation. (**Fig. S6** shows this for a more complex task).

Overall, introducing random synaptic connections can mask the LPU structure but still preserve stable latent computations (**Fig. 5**). Thus, connectome information alone does not necessarily reveal the causal coding dimensions of a neural network. This observation implies that LPU dynamics are not contingent upon an exact low-rank connectivity structure between neurons. Rather, structured low-rank connections superimposed on potentially high-dimensional (functionally inert) connections can support LPUs.

### Causal coding dimensions provide control directions for continuous computations

The preceding results show that tuning relationships and synaptic connectivity can fail to identify which neurons causally influence an LPU. For this reason, recent work has used dynamical and interpretable models such as rSLDS^20,30–32,37,44^, low-rank RNNs^11,33–35^, or deep neural networks^15,78,79^ to infer the underlying dynamics. While such models can serve as *in silico* surrogates for predicting responses to novel perturbations, the re-entrant encoding weights of LPUs provide a more direct principle for population control. By **Theorem 5**, the lasting effect of a perturbation is determined by its projection onto the causal coding dimensions.

To illustrate the control made possible by LPUs, we turn to a regression task in which the goal is to steer a continuous latent variable. Specifically, we trained rank-one basic RNNs to perform delayed addition tasks, in which two scalar inputs provided to the network at two random time points must be summed and reported at the end of the trial (**Fig. 5d**). As we show below, training on this task yields an approximate line attractor that encodes continuous latent variables and therefore constitutes a continuous control scenario. After training, we delivered perturbation pulses to neurons at the midpoint of each trial and compared interventions along several population directions (**Fig. 5d**). This task provides a natural setting for comparing causal and acausal perturbations, because the causal coding dimension for each network is given by an encoding vector, *i.e*., the non-trivial right eigenvector of the synaptic connectivity matrix (cf. Eq. (4)). Further, the learned output weights were strongly correlated with the latent variables by design and were, in practice, orthogonal to the encoding vectors (cosine similarity = 0.006 ± 0.070, mean ± s.d. over 51 networks). Hence, the output weights constituted acausal coding dimensions.

We show the response of an example network to this perturbation experiment in **Fig. 5d-e**. Perturbing along the encoding direction produced a persistent displacement of the latent state and drove the network toward an incorrect final output (**Fig. 5d**). In contrast, perturbing along the readout direction transiently changed the output but did not produce a lasting change in the latent computation (**Fig. 5d**). The phase portrait of the trained network revealed an approximate line attractor as the primary driver of this computation (**Fig. 5e**). Consistent with this geometry, readout-aligned perturbations pushed trajectories away from the slow manifold but were rapidly corrected by transverse relaxation. Encoding-aligned perturbations displaced the latent variable along the computational manifold itself. Across all trained networks, the encoding direction closely matched the trial-averaged locally optimal perturbation direction (**Methods**) and facilitated linear control of the final latent state over several orders of magnitude in perturbation strength (**Fig. 5f**).

Finally, even after adding random synapses between neurons as in **Fig. 5a-c**, encoding weights remained accurate as control directions (**Fig. S6**). Both task performance and controllability were preserved until the synaptic additions drove the network to the edge of stability, where the LPU-contributing weights accounted for about 0.02% of the total synaptic weight power. Overall, these results support the notions that i) the linear encoding map (**P1**) defines practical causal directions for controlling neural computations, and ii) control was effective even when the networks had high-dimensional connectivity matrices with low-rank connections superimposed.

### Theoretical robustness of latent dynamics to changing neural tuning properties

Neither neural tuning nor synaptic connectivity uniquely identifies the causal coding dimensions. Here, we show that the subspaces whose dynamics are preserved across day-to-day changes in neural tuning offer a principled way to identify them. To do so, we first introduce a dynamical model of representational drift and then prove that the latent computations and the underlying coding subspaces are robust to it.

Earlier work on representational drift^4,7^ has often focused on whether behavioral readouts remain invariant. Here, we ask a stronger dynamical question, namely, whether the latent dynamics themselves remain stable. To this end, we developed a dynamical model of drift in networks defined by Eq. (3). Using **Theorem 4** and after some algebra (**Methods**), we express the neural tuning properties in these networks as

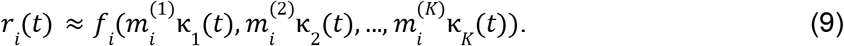

This equation defines the tuning curve of neuron *i* as a function of the latent variables and embedding weights, 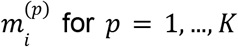. In this notation, the dependence on inputs, biases, and other structural parameters is suppressed. We simulate the drift by modifying the embedding weights associated with the *d*th latent variable, (***m***^(*d*)^ → ***m***^(*d*)^ + Δ***m***). Such a modification generically changes the tuning relationship in Eq. (9). We next ask whether the corresponding latent dynamics remain unchanged. In agreement with experimental findings^7,9^, we find that exact invariance generally does not hold, owing to the nonlinear embedding, but LPUs can remain robust to the first-order changes as we formalize below:

#### Theorem 6

*(Computational robustness to representational drift). Let* ***r***(*t*) *denote the neural activity encoding an LPU with latent variables* ***κ***(*t*). *Let the embedding weights change via* ***m***^(*d*)^ → ***m***^(*d*)^ + Δ***m***. *Under some statistical assumptions (****Methods****), the latent dynamics remain preserved up to second-order corrections* 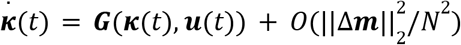 *as long as the drift is purely random or* Δ***m*** *is orthogonal to the causal coding dimensions*.

For any decoder that factors through the latent variables (***Ψ***(***r***(*t*)) = ***Ψ***(***κ***(*t*))), **Theorem 6** implies robustness to representational drift under the same conditions. Given the observation of structured drift in several datasets^53,80^, this robustness may allow time for simple learning mechanisms to compensate for the higher-order (non-vanishing) effects of drift^53^. Practically, this observation also suggests that longitudinal recordings may constrain the encoding directions whose preservation is necessary for stable latent dynamics (**Fig. 6a**).

**Figure 6.**
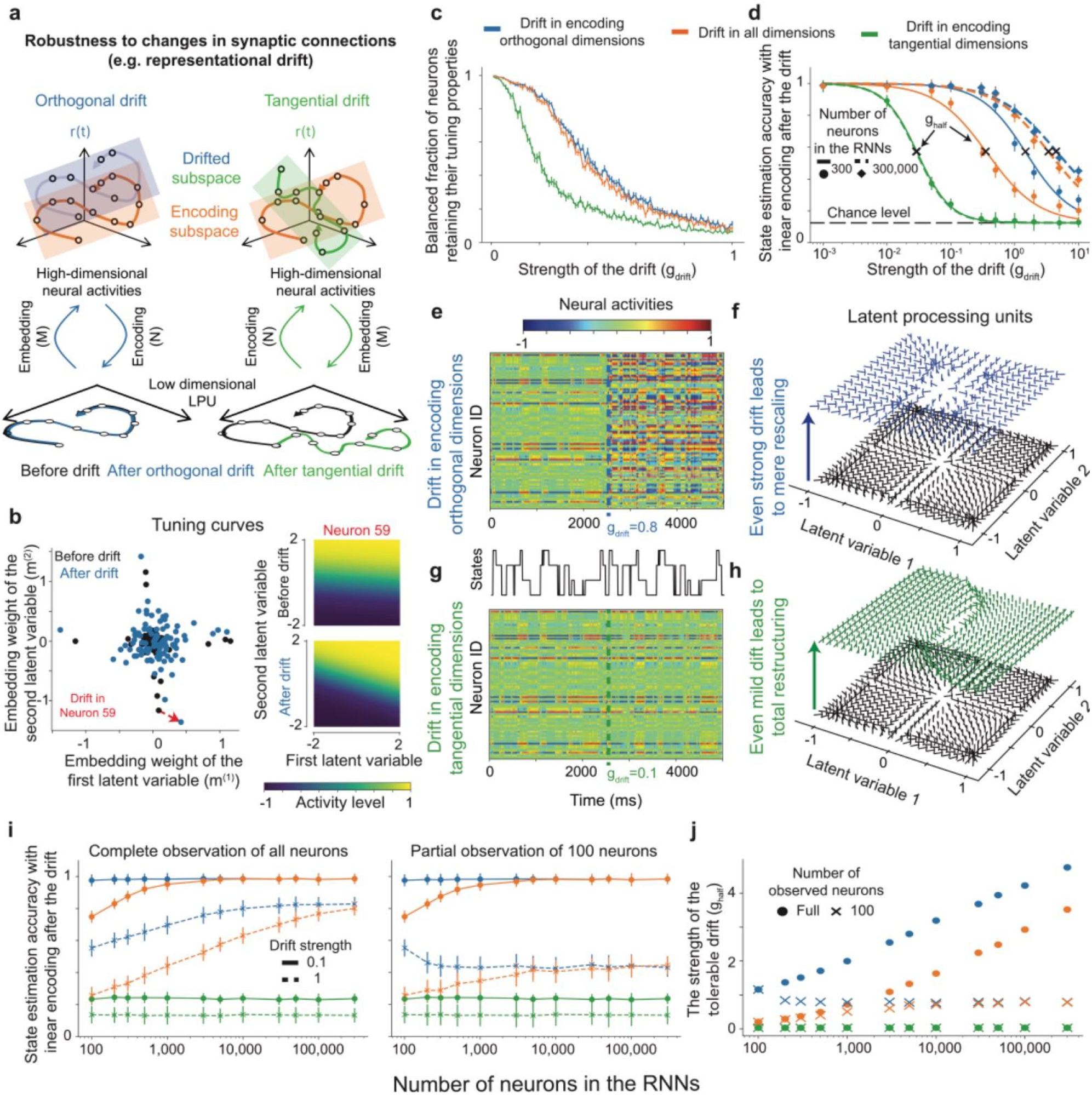
(Theorem 6, Computational robustness to representational drift). LPU dynamics are robust to drift provided the synaptic changes are either purely random or orthogonal to the causally coding subspace. **(a, b)** Linear encoding maps allow LPU dynamics, and not just the behavioral outputs, to be robust to drift in the neural connection weights; this enables RNNs to preserve stable representations even though perturbations to the embedding weights alter neural tuning curves (**Theorem 6**). To illustrate this property, we first noted that the tuning curve of a neuron *i* with respect to the latent variable *j* is defined solely by the embedding weights, 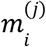, such that 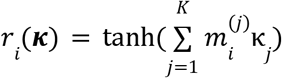 following the definitions in **Figure 4.** Then, simply by shifting the embedding values, we incorporated drift into the neural representations of latent dynamics and studied LPU properties within the augmented network. **(a)** Illustration of representational drift and its potential effects on latent dynamics. To simulate drift, we changed embedding weights either randomly or in a structured way such that the changes were either orthogonal to (termed “orthogonal drift”, labeled blue) or confined to the causally coding subspace (termed “tangential drift”, labeled green). The causally coding subspace is spanned by the re-entrant projections through which individual cells causally impact the LPU dynamics. The hypothesis was that even though the neural activity traces change in both cases (top row), the latent dynamics remain robust only when the drift is unaligned with the causally coding subspace (bottom row, black lines correspond to the representation of the latent dynamics before the drift). Along the trajectories, black-outlined dots denote the progression of the neural and latent dynamics before and after the drift. **(b)** Plot showing the changes in individual cells’ tuning properties as we simulated representational drift as described above. We used the rank-2 RNN from **Fig. 1c–e** that stores 2^2^ = 4 flip-flop states. *Left*: For each neuron (individual data points), we first identified its embedding weights corresponding to each latent variable, providing a pair of outputs (*x-* and *y*-axis values). Black: original network. Blue: after a random perturbation is applied to the embedding weights. *Right*: Plot of the (expected) activity level of an example cell (#59) as a function of the latent variables, *i.e*., its tuning curve. The representational drift shifts its tuning properties. Before the drift, cell 59 selectively codes for the second latent variable; whereas after the drift, its tuning curve shows mixed selectivity for the pair of latent variables. **(c, d)** To quantitatively illustrate LPU robustness to representational drift (**Theorem 6**), we conducted large-scale experiments on low-rank RNNs performing 3-bit flip-flop tasks. We first trained 50 RNNs (100 cells each) to perform the task and then generated larger networks whenever applicable by concatenating and rescaling the learned encoding and embedding weights (**Methods**). We then simulated different forms of instantaneous drift, *i.e*., changes to the embedding weights. To induce a fully random drift, changes were sampled from a zero-mean Gaussian distribution with an s.d. of *g*_*drift*_. To induce targeted drifts, random weights were projected onto directions that were either parallel or orthogonal to the causally coding subspace, with rescaling to maintain constant power. To evaluate the effects of the drift on the computations performed by the RNNs, we defined average state estimation accuracy as the agreement between the target output state and the internal state of the RNN, in which the latter was taken to be the nearest flip-flop state representation in the LPU (*e.g*., the fixed-point attractors) in *L*_2_ norm to the current latent variables. **(c)** To show quantitatively that changing embedding weights leads to representational drift, we plotted the fraction of cells retaining their tuning properties as a function of the drift amplitude, *g*_*drift*_. Since embedding weights define how each cell is activated for a given flip-flop state, we grouped neurons into three distinct groups: positively coding, neutral, and negatively coding. Then, the change in tuning was quantitatively assessed by evaluating the levels of concordance between these groups, before *vs*. after the occurrence of drift, with the assumption that the underlying LPU remains unchanged (*e.g*., assuming acausally coding neurons as in **Figure 5**). Data points: means. Error bars: s.e.m. over 50 simulated networks. **(d)** Average state estimation accuracy, which quantifies the drift in the latent dynamics, after the three types of drift as a function of the drift strength. Linearly encoded LPUs remained robust to drifts in dimensions orthogonal to the causally coding subspace with only a few hundred neurons. Robustness to fully random drifts emerged with large-scale networks. In contrast, tangential drifts along the causally coding subspace consistently resulted in significant decreases in state estimation accuracy from latent variables, regardless of the number of neurons. Here, *g*_*half*_ refers to the drift strength at which the sigmoid fit halves, defined as the “tolerable drift” strength before state estimation accuracies rapidly decline. Notably, in **c**, the drift changes the tuning properties within *g*_*drift*_ ∈ [0. 2, 0. 4], whereas, in this panel, *g*_*half*_ > 1 for the drift-robust network, showing that RNNs were robust to drift even though neural tuning properties changed significantly. Data points: means. Error bars: s.d. over 50 simulated networks. **(e–h)** To study the mechanism underlying the sharp drops in state estimation accuracies following tangential versus orthogonal drifts, we re-analyzed the example RNN with 100 cells from **Fig. 1**, trained to perform the 2-bit flip-flop task. To highlight the contrast, we applied a strong encoding-orthogonal drift with *g*_*drift*_ = 0. 8 and a milder encoding-tangential drift with *g*_*drift*_ = 0. 1. **(e, g)** Similarly to **Fig. 1c**, we show raster plots of activity showing internal neuron activations of an example RNN solving a 2-bit flip-flop task, before and after the drift is applied. Black solid line: the (target) internal flip-flop states of the RNN defined by the inputs. Dashed vertical lines: times at which drifts were applied to the embedding weights. The tuning properties of individual cells changed significantly for strong encoding-orthogonal drift **(e)**, but not for its milder encoding-tangential counterpart **(g). (f, h)** Similar to **Fig. 1e**, we plotted 2D phase portraits of the latent dynamics before and after the drift. Here, arrows show the flows of the latent dynamics, with 4 fixed-points visible before the drift (black phase portraits, the same as in **Fig. 1d**). **(f)** After the drift (blue phase portrait), the LPU structure remained largely intact for the encoding-orthogonal drift despite substantial changes in the tuning properties shown in panel **(e)**, which simply rescaled the latent flow map. **(h)** After the encoding-tangential drift, even though the mild drift caused small changes in the tuning properties of neurons shown in **(g)**, the latent dynamics significantly changed after the drift (green phase portrait). Notably, after the drift, the LPU no longer had the 4 attractive fixed-points necessary to store the flip-flop states. **(i, j)** To quantify the robustness of LPUs to two of the three forms of representational drift, we studied the state estimation accuracies within large-scale RNNs as a function of the number of neurons under full and partial (only 100 out of *N*) observation of neurons. These RNNs were generated using a sampling-based approach from task-trained networks, similar to panels **c-d. (i)** State estimation accuracy, plotted as a function of cell count across different drift amplitudes. While networks grow increasingly resistant to higher levels of drift as neuron numbers rise, this robustness may be masked when only a small fraction of neurons are observed. Data points: means. Error bars: s.d. over 50 simulated networks. **(j)** The plot shows the tolerable drift strength *g*_*half*_, computed from the RNN results shown in panel **i** following the procedure explained in **d**, as a function of the number of cells in the RNN for the three types of drift. Robustness to fully random drift emerged only in networks with more than a few thousand neurons, while encoding-orthogonal drifts could be mitigated with just a handful of neurons. Encoding-tangential drift consistently impaired performance, regardless of the number of neurons. Moreover, when the number of observed neurons was limited to 100, qualitative robustness was still observed in the first two types of drift, though the quantitative values were significantly reduced.

### Robustness of large-scale RNNs to representational drift

To test the predictions of **Theorem 6** in practice and to illustrate our dynamical model of representational drift, we ran simulations of representational drift in large-scale RNNs.

We began with the example RNN from Fig. **1c-e** performing a 2-bit flip-flop task. Before drift, only a few neurons exhibited strong tuning to the flip-flop states, each tuned to a specific input (black dots; **Fig. 6b**). We then modeled representational drift as random perturbations to the embedding weights. After drift, most neurons had altered their tuning (blue dots; **Fig. 6b**). For the rest of this section, we simulated drift in large-scale RNNs in three distinct ways (**Methods**): (i) with fully random Δ***m*** sampled from a Gaussian distribution with zero mean and *g*_*drift*_ s.d., (ii) changes with the same overall magnitude but constrained to remain orthogonal to the causal coding dimensions, and (iii) changes tangential to the causal coding dimensions (**Fig. 6c-j**). By **Theorem 6**, latent dynamics should remain robust to the first two types, but not the third.

To test this prediction, we simulated representational drift by varying *g*_*drift*_ from 0. 001 to 10. Because the original embedding values were ~ 1 (**Fig. S1c**), drift of comparable magnitude substantially changed the tuning properties of individual neurons (**Fig. 6c**). To quantify robustness of latent computations, we estimated the flip-flop states from latent variables (**Methods**) and compared them to the target values (**Methods**). We defined the drift tolerance threshold (*g*_*half*_) as the transition point of a sigmoidal fit to the resulting accuracies (**Fig. 6d**). RNNs with hundreds of neurons were robust to drift constrained to remain orthogonal to the causal coding dimensions, while robustness to fully random changes required thousands of neurons (**Fig. 6d**). By contrast, even networks with 300,000 neurons failed to maintain their latent computations when changes were tangential to the causal coding dimensions (**Fig. 6d**). Together, these observations were consistent with the predictions of **Theorem 6**.

To understand the mechanism for this difference, we revisited the example RNN from **Fig. 1c-e** and applied strong orthogonal (*g*_*drift*_ = 0. 8, **Fig. 6e-f**) and mild tangential (*g*_*drift*_ = 0. 1, **Fig. 6g-h**) drifts. Although strong orthogonal drift produced larger changes in the tuning properties of individual neurons (**Fig. 6e, g**), only the mild tangential drift altered the LPU dynamics qualitatively (**Fig. 6f, h**). Under strong orthogonal drift, the attractive fixed-points shifted toward the origin, but the LPU retained all four internal states (**Fig. 6f)**. Under mild tangential drift, the LPU dynamics were qualitatively altered, and all fixed-point attractors were eliminated through bifurcations (**Fig. 6h**). The persistence of fixed-points under orthogonal drift, despite shifts in their locations, is consistent with the idea that LPUs are robust but not fully invariant (**Theorem 6**).

Finally, we examined how increasing the number of neurons affected tolerance to the three drift types (**Fig. 6i-j**). As expected, even mild tangential changes substantially disrupted latent computations regardless of neuron count (**Fig. 6i**). In contrast, increasing network size improved robustness to orthogonal and fully random drift (**Fig. 6i**). Partial observations from RNNs with up to 300,000 neurons still revealed qualitative properties of the drift tolerance, though not the quantitative values (**Fig. 6j**). Overall, as predicted by **Theorem 6**, purely random changes and those orthogonal to the causal coding dimensions had minimal effects on LPU dynamics, whereas tangential changes disrupted the latent computation.

## DISCUSSION

Biological neural computation presents a series of apparent paradoxes. Computational outputs are reliable, yet their implementation makes use of a constantly changing substrate: neurons whose coding properties drift over time^7,9^. Network outputs, such as motor actions or decisions, are often low-dimensional, yet population activity occupies extrinsically high-dimensional neural manifolds. Many neurons are tuned to behaviorally relevant variables, yet their perturbation can yield no measurable behavioral effect. These observations are difficult to reconcile if neural computation is identified with single-cell tuning^81^, with the geometry of neural activity alone^26^, or with raw synaptic connectivity^82^.

The LPU framework resolves these tensions by treating low-dimensional latent dynamics as the unit of computation (**P3**): its causal variables are set by a linear encoding map (**P1**), expressed through a flexible, nonlinear, drifting embedding (**P2**), and realized in biological synaptic parameters (**P4**). These principles jointly reconcile each paradox in turn, as established across the preceding sections: drift in individual tuning need not destabilize the underlying computation, high extrinsic dimensionality need not imply high-dimensional latent dynamics, and strong tuning need not confer causal influence.

### LPUs generalize low-rank theories of neural computation

The LPUs introduced here build on and extend a recent line of foundational work in computational neuroscience. Previous studies showed that low-rank decomposition can yield latent dynamical systems in specific RNN architectures^11,33–35,77,83^. These studies established important foundations by relating low-rank connectivity between individual neurons to computations in lower-dimensional subspaces, and by proving a universal approximation theorem^35^ analogous to our **Theorem 1**. However, this line of work focused primarily on architectures in which latent dynamical systems were obtained through constraints that impose a linear map from latent variables to neural activities^11,33–35,77,83^. This choice made the theory analytically powerful^34,35,77^, but also constrained neural activity to low-dimensional linear subspaces by construction and imposed corresponding linear restrictions on neural tuning properties.

Low-rank structure, however, is not unique to any particular RNN architecture and is widely observed across physical systems^58^. Motivated by this broader view, we introduced LPUs as a generalization of low-rank RNNs to the larger class of networks described by Eq. (3). Whereas previous studies have proposed separate phenomenological models to explain individual aspects of the observations considered here^3,7^, LPUs provide a unified theoretical framework that derives these properties from a common set of principles. To the best of our knowledge, the specific results formalized in **Theorems 2, 3, 4, 5**, and **6** do not have direct counterparts in the existing low-rank RNN literature. Together, these theorems operationalize low-rank structure as a link between physical circuit parameters and latent computations in biological neural networks.

In line with this discussion, the two architectures described by Eq. (6) also suggest a practical implication for network modeling in systems neuroscience. Basic rate-based RNNs and their low-rank variants can serve as universal approximators of latent computations, but gain-modulated RNNs provide a minimal architecture that can also universally approximate embedding maps, and therefore more flexibly capture neural tuning properties. Thus, when the goal is not only to reproduce behavior or latent dynamics, but also to model neural activity itself, gain-modulated RNNs may provide a more appropriate default than basic rate-based RNNs^84,85^ or more phenomenological models such as rSLDS^31,37^.

### Neural dimensionality need not equal computational dimensionality

A central implication of the LPU framework is that the dimensionality of neural activity and the dimensionality of the computation need not coincide. This distinction bears directly on recent large-scale recordings showing that cortical population activity can exhibit slowly decaying variance spectra, with the variance explained by the *i*th principal component scaling approximately as *i*^−α^ and α ≈ 1 across many cortical areas^39,40^. These spectra have been interpreted as evidence that neural activity is intrinsically high-dimensional. The LPU framework is compatible with this observation but offers a different interpretation of what the high dimensionality reflects. Rather than requiring a correspondingly high-dimensional latent computation, high-dimensional neural activity can arise from the nonlinear embedding of a fixed set of low-dimensional latent variables into a large neural population.

This interpretation follows from the separation between encoding and embedding. The encoding map determines the latent variables that feed back into the computation, whereas the embedding map determines how those variables are expressed in neural activity. If this embedding were linear, neural activity would lie in a (linear) subspace of dimension at most K, and the geometry of the neural population would closely mirror the dimensionality of the latent computation. Nonlinear embeddings can generate curved neural manifolds whose extrinsic linear dimensionality grows with the number of recorded neurons, even when the intrinsic dimensionality of the latent dynamics remains fixed. Thus, high-dimensional activity spectra may reflect the geometry of neural expression, rather than the dimensionality of the computation itself.

This view also reframes the role of neural manifold geometry. Curvature, mixed selectivity, and high-dimensional activity need not be signatures of additional latent variables or hidden behavioral factors. They may instead arise as natural consequences of expressing low-dimensional latent dynamics through heterogeneous nonlinear neurons. Related ideas were suggested in the interpretation of high-dimensional cortical spectra^39^ and later illustrated in a specifically designed RNN model^47^. **Theorem 4** formalizes this observation beyond a particular architecture by showing that nonlinear embeddings can produce extrinsically high-dimensional neural manifolds in a broad class of networks. A key empirical consequence is that estimating the dimensionality of neural computation requires more than measuring the variance spectrum of neural activity. Instead, it requires distinguishing the intrinsic latent dynamics from the extrinsic geometry through which those dynamics are expressed.

### Network scale constrains stable computation over long timescales

The LPU framework also offers a distinct perspective on why biological circuits may rely on very large populations even when smaller artificial networks can solve similar behavioral tasks. Standard RNNs can perform many memory and decision-making tasks with hundreds or thousands of units^34^, whereas biological circuits often engage much larger populations during seemingly simple behaviors^8^. In our framework, network scale is a resource for stabilizing computations over long timescales. **Theorem 2** shows that finite-size variability limits the precision with which a slow latent vector field can be implemented. Because the squared error scales as *O*(β^2^ /*N*), maintaining a fixed level of accuracy requires the timescale separation between neural and latent dynamics to grow no faster than 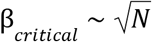. Thus, large populations can be necessary, not because the computation is high-dimensional, but because the same low-dimensional computation must be implemented accurately over behavioral timescales much longer than single-neuron time constants.

This scaling argument complements classical notions of memory capacity. As early as in Hopfield networks^86^, capacity has been characterized by the number of stable states that can be stored in a network, which traditionally scales linearly with network size. Related linear scaling arguments appear more broadly in systems neuroscience, including recent studies of continuous attractor models under constant noise injection^87^ and the number of sequences that can be generated in a recurrent network^88^. Here, the LPU framework predicts a distinct scaling. When there is variability in network parameters, maintaining a stable latent computation over long timescales requires the timescale separation to scale no faster than 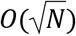. Thus, synaptic variability imposes a stricter sublinear constraint on working-memory computations than classical fixed-parameter capacity arguments. Although neural computation is commonly studied under injected activity noise, this result suggests that intrinsic variability in network parameters can impose separate limits on the stability and duration of latent computations.

Relatedly, this observation is directly relevant to understanding biological neural networks. Since variability of synaptic parameters induces changes in neuronal tuning properties, network scale also affects robustness to representational drift (**Theorem 6**). In our simulations, larger networks became increasingly robust to random changes in embedding weights, whereas drift tangential to the causal coding dimensions disrupted latent computations even in very large networks (**Fig. 6**). Thus, increasing population size can buffer some forms of neural variability but does not make computation generically insensitive to all changes in tuning. Stability depends on the relationship between the direction of drift and the causal coding dimensions. This observation provides a more specific account of robustness than the general idea that larger populations average out noise.

### Experimental tests of computation with latent processing units in biological systems

Our framework makes several experimentally testable predictions concerning the effective perturbation directions in neural networks. As described above, while neural populations often exhibit high-dimensional activity patterns, our theory predicts that effective perturbations, those that meaningfully influence latent dynamics and the subsequent behavior, should be confined to a much lower-dimensional linear subspace (equal to *K* ≪ *N*, **Fig. 5**). This prediction is partially supported by recent experiments^50^, in which perturbations based on tuning properties of individual neurons failed to disrupt behavior, while choosing targets based on their network contributions did.

A direct test of this prediction remains elusive but becomes particularly relevant with recent advances in single-cell manipulation techniques. Using optogenetic manipulations and targeted interventions, one could in principle test whether effective perturbations of neural activity in awake animals are confined to low-dimensional subspaces. Then, by examining how large perturbations decay along orthogonal dimensions, one could determine whether the high linear dimensionality observed in neural recordings arises from nonlinear embedding of low-dimensional latent dynamics as opposed to high-dimensionality of coding variables, some of which may be uncorrelated with behavior. While these experiments could provide direct evidence for our theoretical framework, such a distributed perturbation would require new generation single-cell intervention tools that allow fine-grained optogenetic access to thousands of individual cells.

Beyond these interventional tests that would require sophisticated equipment, more immediately feasible tests exist. Our theory makes specific predictions about neural manifold geometry. Even in seemingly simple tasks, our theory predicts that neural activities will reside in curved manifolds rather than affine subspaces (**Fig. 4**). Preliminary evidence comes from experiments on spatial navigation, in which neural codes of grid cells form toroidal manifolds^27,89^ that are naturally curved. While previous work has characterized the high linear dimensionality of neural activities in passive viewing tasks^39^, manifold curvature in visual coding remains underexplored, partly due to challenges in estimating manifold dimensionality under noisy conditions (though also see Ref. 25).

### Limitations

Several limitations circumscribe the scope of the present framework. First, throughout this work we primarily focused on a weak-scaling regime in which individual encoding weights scale as *N*^−1^. This choice is standard in the low-rank RNN literature^34,35^ and ensures that latent variables remain finite and self-averaging as the number of neurons increases. It is therefore the natural regime for studying stable latent computations implemented by large populations. However, recurrent neural networks are also often studied in a different scaling regime, in which random connectivity scales as *N*^−0.5^ and can generate strongly fluctuating, chaotic, and high-dimensional dynamics^77,90^. While we did consider the example of low-rank chaotic networks (**Fig. S5**), the present theory does not attempt to unify these regimes or explain computation in networks whose dynamics are dominated by such strongly fluctuating activity.

Second, direct identification of LPUs in biological data remains challenging. Our results specify the mathematical objects that define an LPU, including latent variables, nonlinear embeddings, and causal encoding directions, but estimating all of these objects from neural recordings is difficult. This difficulty is especially pronounced under the conditions of partial observation, in which only a small fraction of the relevant neural population is recorded. In such settings, behaviorally relevant variables may be decodable even when the underlying causally coding subspace and latent dynamics remain only partially identifiable (cf. **Theorem 2**).

Third, our model of representational drift isolates one interpretable abstract mechanism. In this model, changes in embedding weights alter neural tuning and either preserve or disrupt the causal coding dimensions. This abstraction yields clear predictions about random, orthogonal, and tangential drift, but it does not specify the biological processes that generate these changes. For example, a mechanistic mapping between embedding parameters and synaptic connections could incorporate mechanisms such as synaptic turnover, sparse activity, homeostatic plasticity, or task-dependent remodeling. We have not attempted to model these processes here. Moreover, our treatment does not address the sources of apparent or genuine representational change. For instance, earlier work has shown that changes in behavioral patterns can be misinterpreted as changes in neural representations^91^, and that local learning rules can induce representational change^92^. Modeling these sources of change within our framework remains an open challenge.

Finally, the present work characterizes the structures that can support robust latent computation but does not provide a complete theory of how those structures are learned. The theorems establish the existence, readout properties, geometry, perturbation sensitivity, and drift robustness of LPUs, but do not specify the biological learning rules by which re-entrant encoding weights and nonlinear embeddings are formed. Prior work that used backpropagation through time suggests that even full-rank networks tend to learn functionally low-rank components^33,61,93^, though connecting these findings to biologically plausible learning processes remains to be addressed.

### Outlook

Although we illustrated LPUs with simple examples here, applying the framework to complex networks could help compare the solutions that different architectures arrive at. For instance, previous work on low-rank RNNs (using a different RNN architecture) revealed similar attractors (as in **Fig. 1**) in low-dimensional latent subspaces when solving memory tasks^11,33–35^. While earlier studies primarily used fixed-point analysis to compare learned computational strategies across trained networks^94^, comparing full LPU dynamics, not just equilibrium states represented by fixed-point attractors, may offer a more precise and general framework for studying the universality of neural computations across architectures and experimental conditions.

A further question is whether the rich geometry generated by nonlinear embeddings can itself serve as a computational resource. In our formulation, behaviorally relevant outputs are decoded from latent variables (**P3**), and linear decoders can but do not have to approximate nonlinear readouts of those variables with vanishing error in sufficiently large networks (**Theorem 3**). **Theorem 4** complements this result by showing that neural activity itself can also become an extrinsically high-dimensional nonlinear function of latent variables with ***r***(***κ***) ≈ ***φ***(***κ***). Thus, barring further implications for the allowed drift directions under **Theorem 6**, the nonlinear embedding could provide a rich feature expansion of the latent variables as a natural consequence of the LPU construction. An important future direction is to determine how this geometric richness supports decoding of more complex behavioral or cognitive variables beyond the categorical variables considered in **Fig. 3** (also see Refs. 38,46).

Another future direction involves extending our analysis to spiking recurrent neural networks (sRNNs). Prior work introduced a training paradigm in which latent factors, interpreted as time series data, serve as computationally central elements for training spiking networks^16^. However, it remains unclear whether a self-sufficient and universal latent dynamical system can emerge within such spiking architectures^95,96^. Nevertheless, incorporating LPUs into biologically realistic spiking networks could bridge the gap between theoretical models and neural circuit implementations, offering a promising avenue for further biological investigations.

Finally, LPUs, if implemented by biological networks, have direct implications for the development of brain-machine interfaces (BMIs)^24,97^. As discussed above, while biological systems require large numbers of neurons to sustain complex dynamics, manipulating the information stored in these systems could be achieved through relatively low-dimensional control signals. Specifically, it may be possible to interfere with low-dimensional LPUs by choosing the right neurons and perturbing them in a direction aligned with their encoding weights. Moreover, the linearity of encoding further suggests that local field potential recordings, arising from the linear summation of neural activity, may be as effective as single-unit recordings in decoding information from the brain^98^, especially if the encoding weights are spatially smooth. This insight implies that high-performance BMIs may require, not necessarily increased spatial resolution, but theoretical advances that enable the extraction of drift-robust encoding dimensions. Overall, LPU-based algorithms, designed to track the causally coding subspaces and compensate for occasional drifts, could potentially lead to significant improvements in BMI stability and performance.

## ACKNOWLEDGEMENTS

We thank Vasily Kruzhilin, Itamar Landau, Mert Yuksekgonul, Kanaka Rajan, Omri Barak, Sarah Kushner, Mathilde Papillon, Ali Cetin, Xavier Gonzalez, Adrian Valente, Chong Chen, and Alperen Cimen for helpful discussions. NM acknowledges funding from NSF grant 2313150, and the Noyce Foundation. MJS acknowledges funding from the Simons Collaboration on the Global Brain and the Vannevar Bush Faculty Fellowship Program of the U.S. Department of Defense. FD received funding from Stanford University’s Mind, Brain, Computation and Technology program. This research was supported in part by grant NSF PHY-2309135 and the Gordon and Betty Moore Foundation Grant No. 2919.02 to the Kavli Institute for Theoretical Physics (KITP). PY was additionally supported by the Shanghai Pilot Program for Basic Research - FuDan University 21TQ1400100 (22TQ019) and the Foundation of the Shanghai Municipal Education Commission No. 24RGZNA01.

---

**Figure S1.**
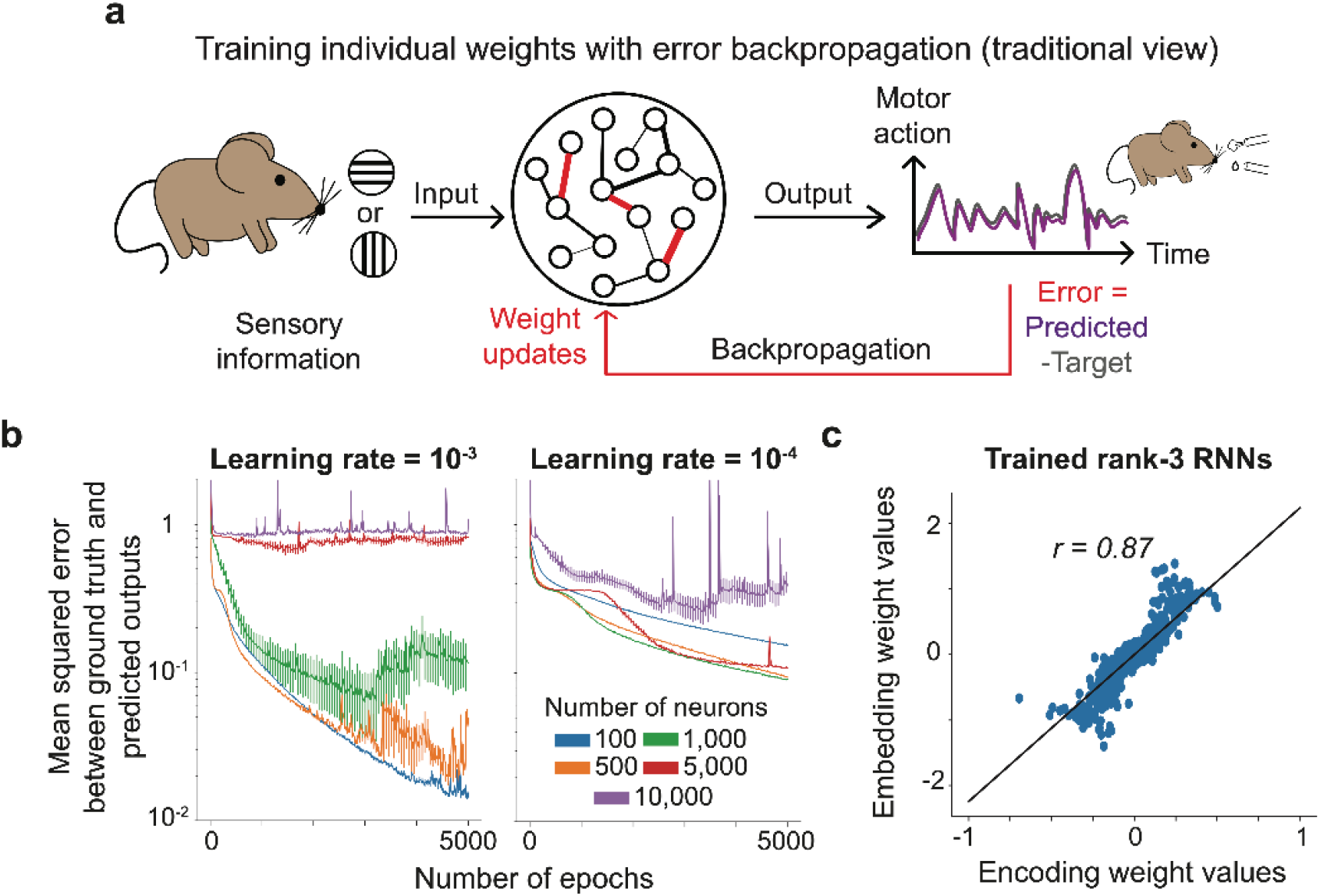
Motivating a training-free approach for generating large-scale RNNs capable of storing flip-flop states. **(a)** A schematic of credit assignment using backpropagation through time, illustrated for a perceptual decision-making task in which a fictional mouse learns to discriminate horizontal vs. vertical visual cues. The mouse brain is represented by a neural network whose synaptic connections are trained such that the mouse can perform a predefined action to signal its choice. The traditional view of training neural networks focuses on minimizing a loss function by propagating error signals (here represented by the difference between predicted and target motor actions) back through time and updating the weights accordingly. In this view, neural activities are relevant only insofar as they contribute to matching predictions to targets. Training large networks using this paradigm requires extensive computer memory to store and compute weight updates, among other demands. **(b)** RNNs with varying numbers of neurons were trained using backpropagation through time to perform 3-bit flip-flop tasks. The plots show the loss function as a function of training epochs for two learning rates (left vs. right), with distinct colors denoting networks of different sizes. For a given learning rate, training larger RNNs led to instabilities, whereas decreasing the learning rate increased the time to convergence. Coupled with the increased computational and memory demands, this makes the traditional training approach infeasible for networks substantially larger than those shown here. Trained RNNs were full-rank and no other constraints were enforced, to rule out trivial explanations of the observed learning deficiencies. Solid lines: means. Error bars: s.e.m. over 20 networks. **(c)** To assess the synaptic structure of emergent LPUs in RNNs trained to store flip-flop states, we analyzed 50 networks with 100 neurons each. RNNs were trained to perform 3-bit flip-flop tasks using the two-step paradigm described in the caption of **Fig. 1b**. Following rank distillation, each network was rank-3. The plot shows the aggregated encoding and embedding weights across all networks and neurons, with each dot representing the encoding-embedding weight pair of a single neuron belonging to a single flip-flop channel. The two quantities were significantly correlated (*r*=0.87, *p*<0.001). We used this observation to generate RNNs capable of solving 1-bit flip-flop tasks. Specifically, we sampled encoding and embedding weights for each neuron from a two-dimensional zero-mean Gaussian distribution with a prescribed positive correlation between the two weight types (see **Fig. 2** and **Supplementary Note 2**).

**Figure S2.**
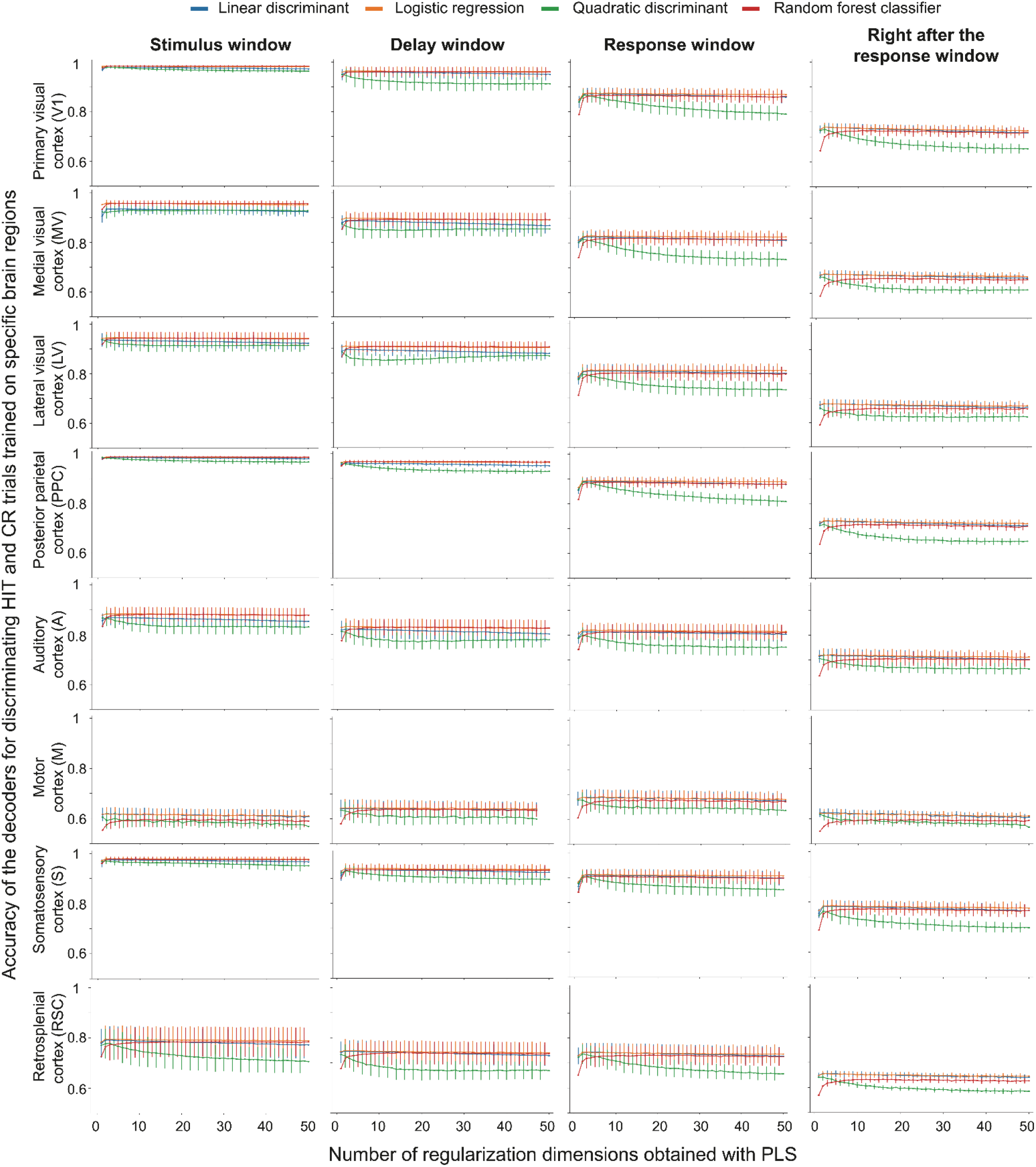
Decoding of trial identity using neural activities from eight neocortical regions. Accuracy results for decoding HIT versus CR trials using various classifiers trained on neural data from different cortical regions, including the primary visual cortex (V1), medial visual cortex (MV), lateral visual cortex (LV), posterior parietal cortex (PPC), auditory cortex (A), motor cortex (M), somatosensory cortex (S), and retrosplenial cortex (RSC). Each classifier – linear discriminant, logistic regression, quadratic discriminant, and random forest – was trained with a varying number of regularization dimensions obtained using partial least squares (PLS). Results are shown for multiple time windows: stimulus, delay, response, and immediately after the response. Each plot indicates mean accuracy across 100 training and testing splits, with s.e.m. shown across the six mice.

**Figure S3.**
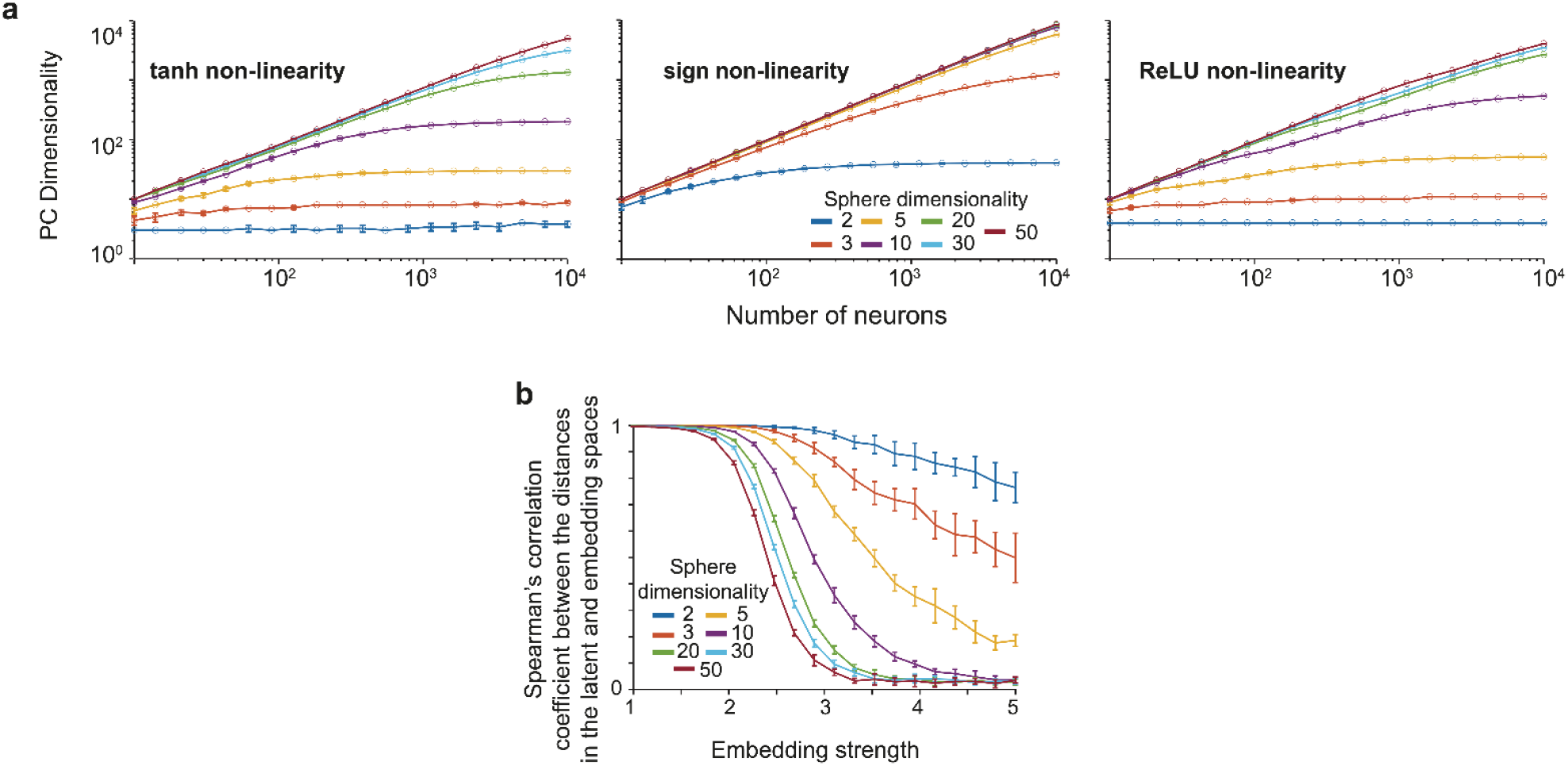
Linear dimensionality of neural manifolds in static embedding of unit spheres. **(a)** We performed the analysis in **Fig. 4b** for three distinct nonlinearities: tanh, sign, and ReLU. **(b)** The plot shows the Spearman’s correlation between the distances in the latent and embedding spaces when a sine nonlinearity is used as in **Fig. 4b**. Increasing the embedding strength beyond a certain threshold resulted in embedding manifolds that failed to preserve the distances between points in the latent space.

**Figure S4.**
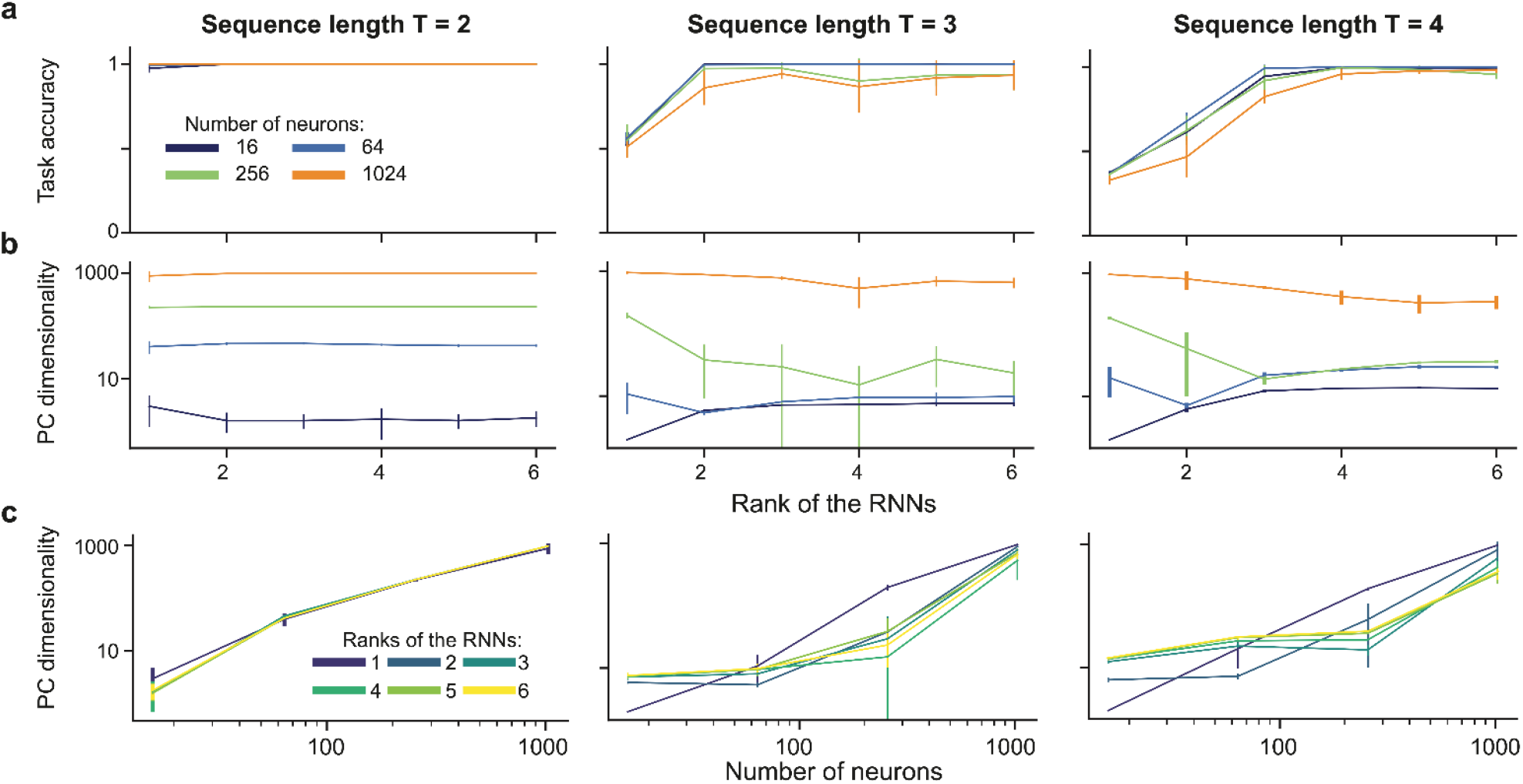
Linear dimensionality of neural activities in RNNs trained to perform sequence sorting. We varied the sequence length for RNNs performing a sequence sorting task and measured the resulting linear dimensionality. Points and bars represent the mean and standard deviation across 10 experimental repetitions with different random seeds. **(a)** Task accuracy plateaued once the RNN rank matched the task dimensionality, *T* − 1, where *T* is the sequence length of the sorting task. This plateau occurred independently of neuron count. **(b)** The extrinsic dimensionality of the neural manifold remained constant across increasing RNN ranks, unaffected by the task dimensionality *T* − 1. **(c)** The extrinsic dimensionality of the neural manifold increased linearly with the number of neurons, independent of RNN rank and task dimensionality.

**Figure S5.**
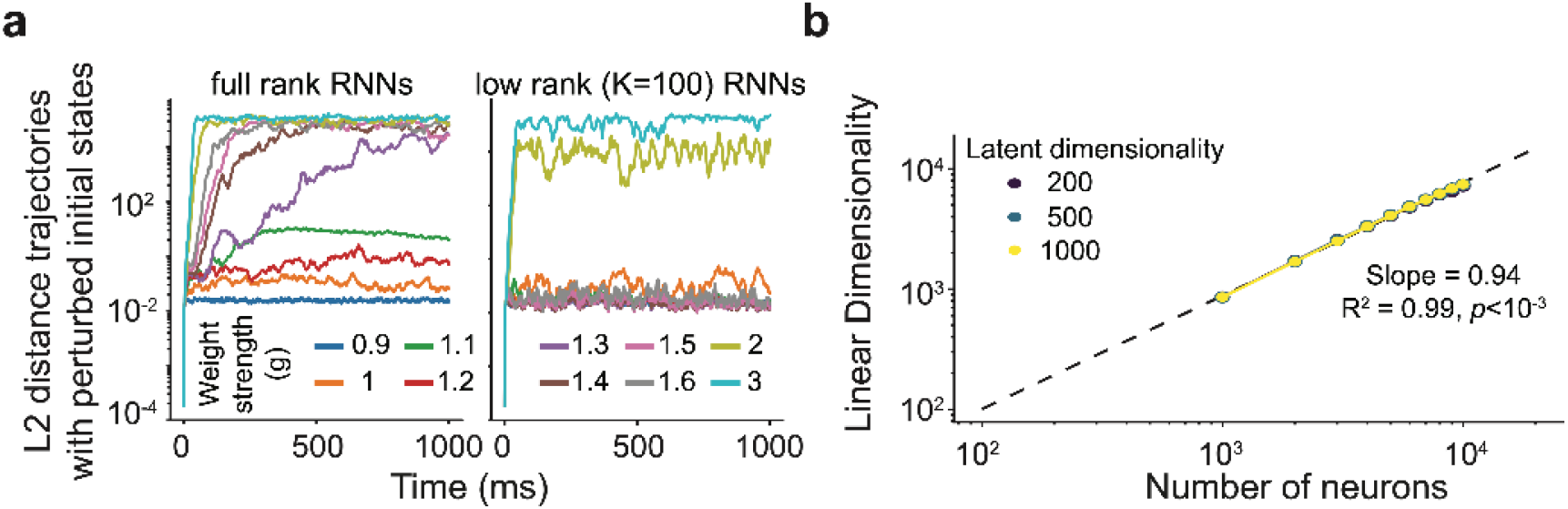
Linear dimensionality of neural activities in randomly connected low-rank RNNs. **(a)** When connections between neurons are randomly drawn from a Gaussian distribution with zero mean, the networks can sustain spontaneous activity and exhibit chaotic behavior. We designed a similar random sampling process for the weight matrices of low-rank RNNs. Even when only *K* = 100 out of 1, 000 ranks of the weight matrix were occupied, the RNNs were able to sustain spontaneous, diverging neural trajectories. Each line represents a single network. **(b)** Low-rank RNNs (*g* = 15) show a nearly linear increase in their linear dimensionality with added neurons, displaying little sign of saturation. Line: mean. Error bars: s.d. across three runs.

**Figure S6:**
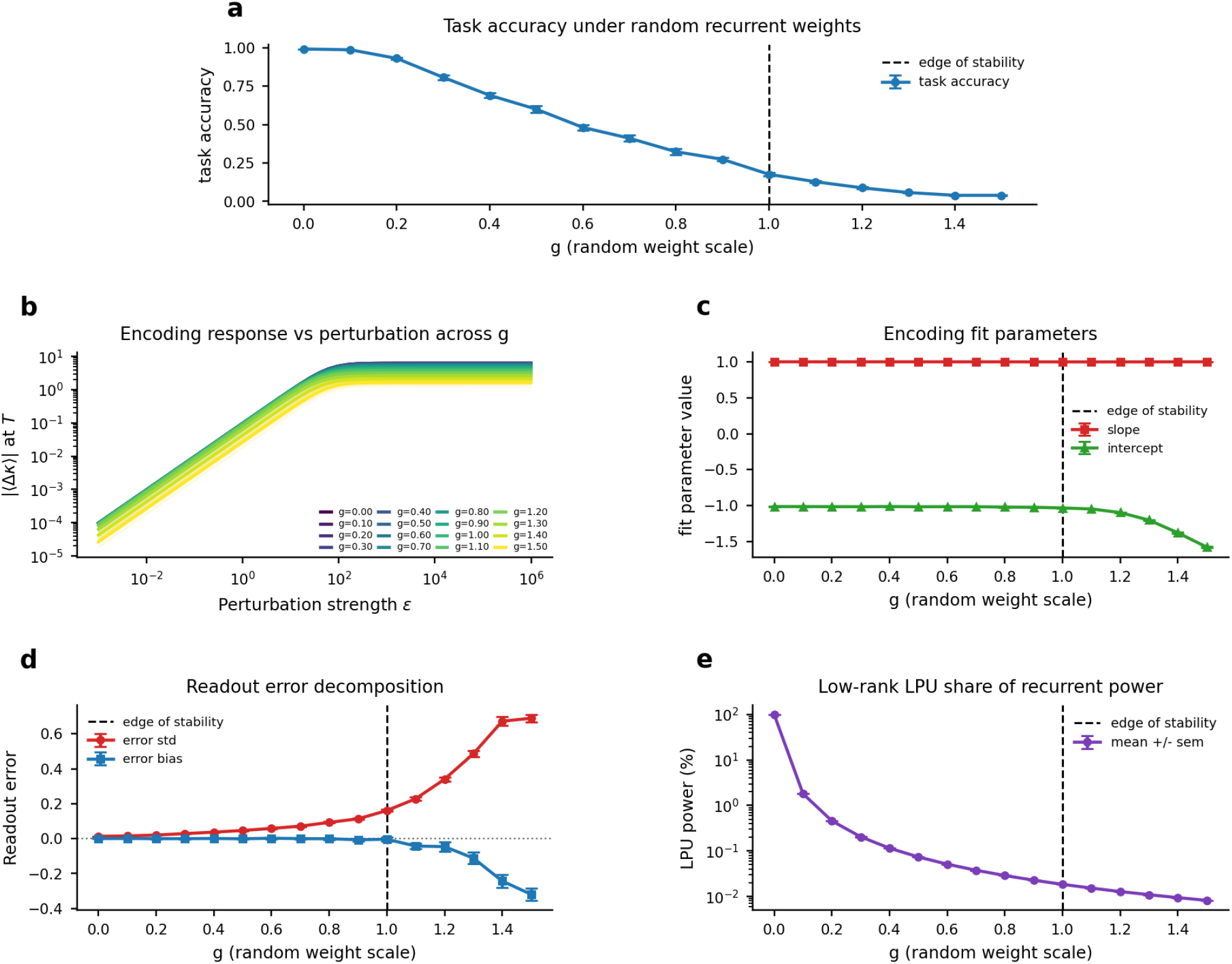
Re-entrant encoding weights enable continuous control of latent variable values even when the connectome consists primarily of random connections. To illustrate that one can continuously write into latent variables through the casual coding directions, we revisited the delayed addition task from **Figure 5d**. This task, when solved by a rank-one RNN, generates approximate line attractors in the one-dimensional LPU as illustrated in **Figure 5e**. The main figure has shown the results when the connectome was also one-dimensional with contributions coming only from LPU-subserving synaptic weights. Here, we augment the connectome via ***W*** = ***W***_*rand*_ + ***mn***^*T*^, where ***W***_*rand*_ is a random component whose elements are sampled from a zero-mean Gaussian distribution with variance *g*^2^/*N* and ***m, n*** are embedding and encoding weights, respectively. For each instantiation of the network, we used the learned ***m, n*** values while sampling the ***W***_*rand*_ component randomly. Similar to **Figure 5d-f**, we perturbed the augmented network through a normalized direction parallel to the encoding weights ***n*** (with the strength ε) at the half-point into the trial, *i.e*., *t* = *t*_1/2_. We then computed the deviation of the latent variable at the end of the trial *t* = 125 ms. All results are shown across 51 well-trained RNNs. **(a)** We first quantified the task accuracies (left y-axis) against the strength of the added random connections (x-axis). The task accuracy has steadily declined with increasing *g*, reaching near-zero values slightly beyond the edge of stability at *g* = 1 (dashed vertical black line). **(b, c)** We next asked whether continuous control of the latent variable remains possible as random connectivity is added, despite the accompanying decrease in task accuracy. We therefore repeated the control experiments from **Figure 5f** using the encoding weights for control. **(b)** We plotted the perturbation-response curves as in **Figure 5f** across varying values of *g*. Across a broad range of *g*, the perturbation response (|Δκ|) scaled approximately linearly with perturbation strength over several orders of magnitude before saturating for large perturbations. The curves remained nearly overlapping up to the edge of stability, indicating that random connectivity had little effect on the efficacy of continuous control within this regime. **(c)** To quantify the observation in panel **b**, we fitted a log-log curve to the perturbations via *log*_10_ |Δκ(*t* = 125 *ms*)| ~ *m log*_10_ ε + *b*, where a unity slope (*m* = 1) would signal a proportional control (|Δκ| ∝ ε) and the intercept (specifically, 10^*b*^) determines the gain. The slope remained close to unity throughout, whereas the gain began to decrease only slightly beyond the edge of stability. Thus, continuous control through the encoding direction remains effective up to, and slightly beyond, the edge of stability, despite the progressive decline in task accuracy. **(d)** We next asked why task accuracy decreases even while continuous control remains effective. One possibility is that random recurrent connectivity introduces variability into the latent dynamics, causing errors to accumulate over time without producing a systematic displacement of the computation. To test this possibility, we decomposed the readout error across networks into its bias and standard-deviation components and plotted both as functions of *g*. The standard deviation increased progressively with *g*, closely paralleling the gradual decline in task accuracy observed in panel **a**. In contrast, substantial systematic bias emerged only after the edge of stability was crossed. These results suggest that the initial loss of task accuracy primarily reflects increased variability in the latent dynamics, rather than a loss of the network’s capacity for continuous control. Beyond the edge of stability, the emergence of systematic bias indicates a qualitative destabilization of the computation, also consistent with panels **b-c. (e)** Finally, we quantified the percentage contribution of the low-rank LPU component to the total recurrent connectivity via the formula 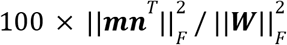 and plotted against increasing *g*. Already at *g* = 0. 1, which had only a minimal effect on task performance in **panel a**, the LPU component accounted for only a few percent of the total recurrent weight magnitude. At the edge of stability, *g* = 1, its contribution had decreased to 0.0184%.

**Table S1.** An overview of existing research and relevance to this work.

| Description | Latent variables | Identifiability and behavioral readouts | Causal predictions and neural activity manifolds | Tuning curves and representational drift | Biological relevance |
| --- | --- | --- | --- | --- | --- |
| Latent processing units | LPUs provide a causal model of latent dynamics. **Theorem 1** guarantees universal approximation capabilities, whereas **Theorem 2** introduces practical limits, *i.e.*, large-scale populations are needed for computations with extended timescales. | **Theorem 3** guarantees the optimality of linear readouts, and that they do not require any form of latent variable identification, thereby not suffering from identifiability issues. Latent variables themselves are linearly identifiable. | **Theorem 4** explains how low-dimensional LPUs can lead to high-dimensional neural activities with non-trivial manifold geometries. Deviations to these coding manifolds decay exponentially fast (**Theorem 5**). | LPUs can model non-trivial tuning curves due to the nonlinearity of the embedding map, which enables numerical simulations of representational drift in RNNs. **Theorem 6** guarantees that not only behavioral readouts, but also the latent dynamics could remain robust to the drift. | LPUs are designed with biologically relevant motivations, and can explain several experimentally observed phenomena. |
| Low-rank RNNs<sup>11,33–35</sup> and latent circuits<sup>36</sup> | The first line of work to establish the connection between low-rank connectivity matrices and low-dimensional coding subspaces. In a particular RNN architecture, these works also introduce a causal model of latent dynamics. An equivalent of **Theorem 1** exists for these models. | Linear readouts can circumvent identifiability issues, though no known results on their efficiency. Our **Theorem 3** could potentially extend to this architecture, though with some restrictions. Latent variables are linearly identifiable. | The coding subspaces lie on affine subspaces, *i.e.*, flat manifolds. Similar to our **Theorem 5**, deviations to these subspaces decay exponentially. | Tuning curves are defined as linear maps from latent variables. No known result on representational drift, and neurons have only linear tuning properties. | Analytical constraints lead to several unrealistic restrictions: coding dimensions are affine subspaces, tuning curves are linear, implications for representational drift currently unknown. Assumptions contradict known results. |
| Recurrently switching linear dynamical systems<sup>20,30–32,37,44</sup> | First line of work to prove the existence of line attractors in the brain with intervention experiments. rSLDS constitutes a causal model of latent dynamics. No known result on universal approximation, though in the limit of infinitely many internal states, it may be possible. | Linear readouts can circumvent identifiability issues, though no known results on their efficiency. Latent variables are linearly identifiable. | The coding subspaces lie on affine subspaces, *i.e.*, flat manifolds. | Linear tuning relationships, with no known discussion of representational drift. It is unclear whether our theorems cover these architectures. | rSLDS framework often considers the learned networks as effective models of brain activities, rather than making claims on the biological processes. Similar to above, the linear map from latent variables significantly constrain the manifold geometry and the tuning curves, though see Refs<sup>29,100</sup>. |
| Deep learning based models<sup>14,28,29</sup> | Most models do not constrain latent dynamics, and some enforce simple dynamics such as linear dynamical systems. For the latter, it is not clear if the models have universal approximation properties. | In some cases, linear identifiability of latent variables can be achieved. Since encoding in these models is often nonlinear, identifiability or efficiency of linear readouts we have proven in our work do not naturally extend to these models. | NA | NA | NA |

**Table S2.**
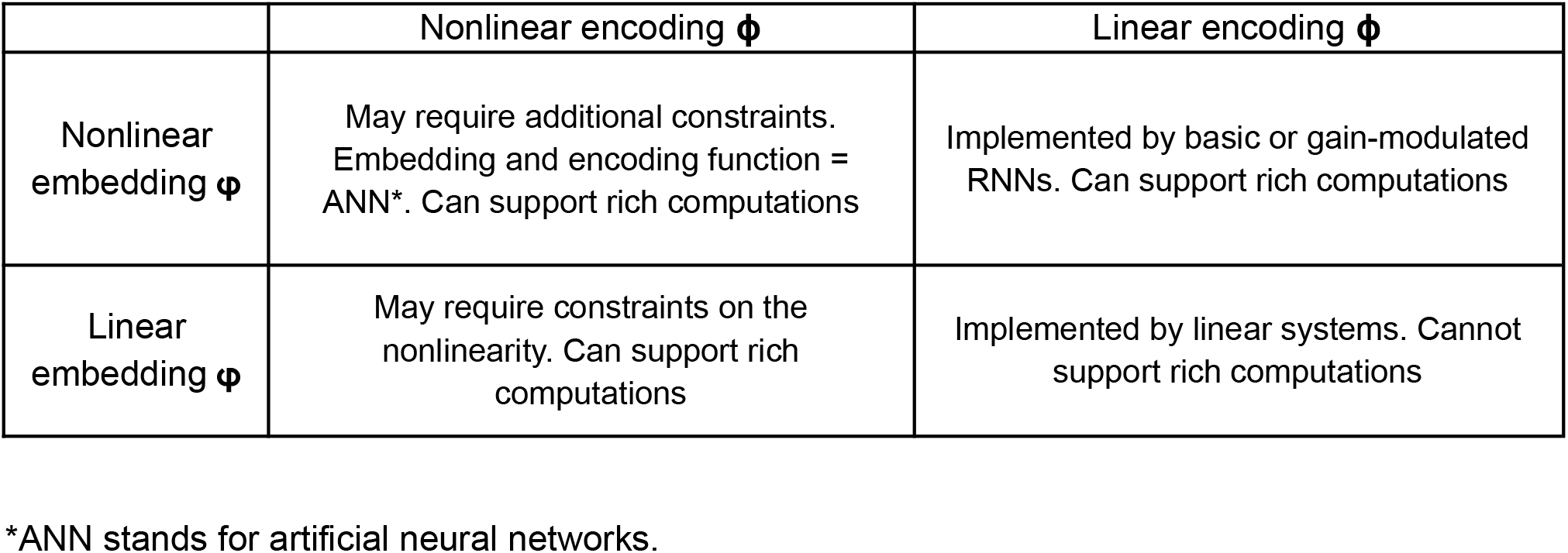
A summary of encoding and embedding types for dynamical systems.

## METHODS

### Mathematical notation

Throughout the methods section, we use boldface characters for vectors (lowercase) and matrices (capitals) and non-boldface characters for scalars. For instance, ***r***(*t*) ∈ ℝ^*N*^ refers to the vector containing the neural activities of *N* neurons at time *t*, whereas *r*_*i*_ (*t*) is the activity of the *i*th neuron. Similarly, ***κ***(*t*) ∈ ℝ^*K*^ refers to the activities of *K* latent variables at time *t*. The notation (*m*^(*p*)^, *n*^(*p*)^) refers to the random variables defining the embedding and encoding weights for the *p*th latent variable κ_*p*_ (we implicitly assume *p* = 1, …, *K* unless explicitly stated otherwise), whereas (***m***^(*p*)^, ***n***^(*p*)^) refers to the embedding/encoding vectors actualized in an RNN across *N* neurons. For *K* = 1, we simply refer to these variables as (*m, n*) and (***m, n***), respectively. We use the matrices, ***M*** ∈ ℝ^*N*×*K*^ and ***N*** ∈ ℝ^*K* ×*N*^, to jointly refer to all the embedding and encoding weights, respectively. For continuous-time representation of variables (*e.g*. consider a variable *a* that depends on time), we use the parenthesis notation *a*(*t*) with respect to the continuous time variable *t*, whereas for discrete time representation of variables, we use the bracket notation *a*[*s*] with respect to the discrete time variable *s*.

### A dynamical system framework for modeling neural computation

What distinguishes recurrent neural networks (RNNs) from their feedforward counterparts is the ability to model temporally varying computation. Studying temporal dynamics requires understanding dynamical systems, which is our starting point as we present the mathematics behind our latent processing units (LPUs). This exposition yields an *interpretable* and *rigorous* definition of LPUs, and generalizes existing work on a restricted class of low-rank RNNs^11,33–35^ and recurrently switching linear dynamical systems (rSLDS)^20,30–32,37,44^ to a broad class of artificial and biological networks. We prove that with some basic mathematical assumptions, this class is capable of universal computation and provide methodological details for the simulation experiments we performed with RNNs trained on flip-flop tasks.

#### Formalization of latent computations via an encoding-embedding framework

Latent computation framework builds on a significant assumption, *i.e*., neural computation results from the time evolution of a high-dimensional dynamical system, which follows a generic equation:

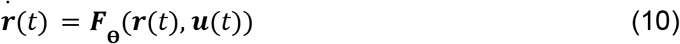

where ***r***(*t*) ∈ ℝ^*N*^ are the state variables of the dynamical system; 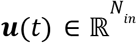 are inputs provided by the user to control the system; **θ** (of varying dimensions based on architecture) are the parameters of the system and ***F***: 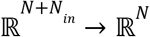 is the “flow map” that defines the time dynamics. Here, the state variables, ***r***(*t*), correspond to the activity of *N* neurons in a neural network. The parameters of the dynamical system, **θ**, would for instance include the combinations of synaptic connections between these neurons and the weights of the external inputs. Throughout this work, we use ***W***^*rec*^ ∈ ℝ^*N*×*N*^ to refer to the former, and 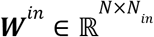 to refer to the latter. Lastly, in this biological setting, ***F***(·) defines the time dynamics of neural activities in the biological network.

In many RNNs, individual units, *e.g*., neurons, have predefined input-output relationships for transforming electrical currents into membrane voltages and subsequently to action potentials^101^. Even though these individual units are constrained, their populations can provide complex network outputs and model dynamics of experimentally observed variables^42^. In this work, to formalize the ability of a broad class of RNNs’ to model low-dimensional dynamical systems, we start by proposing the concepts of encoding and embedding in dynamical systems:

##### Definition 1

*(Encoding and embedding in dynamical systems). Let* ***r***(*t*) ∈ ℝ^*N*^ *be the state variables of a high-dimensional dynamical system with time dynamics as in Eq. (10). Let* ***κ***(*t*) ∈ ℝ^*K*^ *be state variables that constitute another, lower-dimensional, dynamical system with time dynamics* 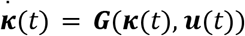 *with* ***G***: 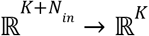 *its flow map, and K* ≪ *N* + *N*_*in*_. *Then, the dynamical system with variables* ***κ***(*t*) *is said to be an encoding of the dynamical system with variables* ***r***(*t*) *if:*

1. *There exist a differentiable encoding map* ***ϕ***: ℝ^*N*^ → ℝ^*K*^ *and an embedding map* ***φ***: 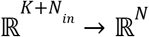 *that jointly satisfy the relationship:*

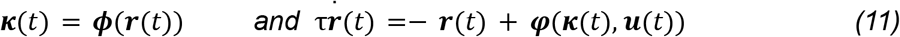
2. *The encoding and embedding maps satisfy the following property such that* ***κ***(*t*) *constitutes a dynamical system:*

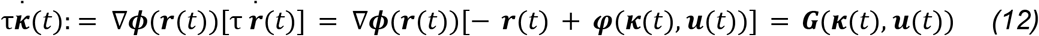 *where* τ > 0 *is the neuronal time constant*, [∇ϕ]_*ij*_ = *∂*ϕ_*i*_ /*∂r*_*j*_ *is the Jacobian of* ***ϕ***(***r***(*t*)), *and* ***G***(***κ***(*t*), ***u***(*t*)) *does not explicitly depend on* ***r***(*t*).

This definition states that for a set of variables, ***κ***(*t*), to be considered as embedded in a high-dimensional dynamical system, ***r***(*t*), their time evolutions should both be self-consistent (with the exception of external input variables, which are assumed to be predefined) and the former dynamical system should be recoverable from the latter. Although the encoding is a map, ***ϕ***, between state variables, we emphasize that the embedding map, ***φ***, only constrains the time derivative 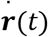. This allows a redundant representation, *i.e*., more than one ***r*** values can correspond to the same ***κ***. In contrast, as is done in standard literature, a static embedding function of the form ***r***(*t*): = ***φ***(***κ***(*t*), ***u***(*t*)) would uniquely define an ***r*** corresponding to a particular ***κ*** and be in that sense more restrictive. Hence, unless explicitly stated, we assume ***φ***(·) is a dynamical embedding.

#### Defining latent processing units

There are several trivial pairs of dynamical systems that can satisfy these encoding-embedding relationships. For example, consider ***κ***(*t*) = ***ϕ***(***r***(*t*)) = ***c*** with some vector ***c*** ∈ ℝ^*K*^. Technically, this would constitute a dynamical system for ***κ*** since 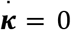 does not depend on ***r***(*t*). Such a system becomes a leaky integrator for the input, 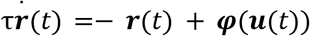 However, this trivial latent dynamical system cannot solve diverse computational tasks. Therefore, our definition of an LPU, *i.e*., the putative substrate of neural computation, cannot solely depend on the ability of the latent dynamical system to satisfy the encoding and embedding relationship. Instead, we define the LPUs as follows:

##### Definition 2

*(Latent processing units). We define low-dimensional dynamical systems, spanned by* ***κ***(*t*) ∈ ℝ^*K*^, *as LPUs if they satisfy two conditions:*

1. ***Accessible embedding of self-sufficient latent dynamics:*** *The low-dimensional dynamical system variables*, ***κ***(*t*), *are embedded in high-dimensional neural activities*, ***r***(*t*) ∈ ℝ^*N*^, *following* ***Definition 1***. *With this construction*, ***κ***(*t*) *are both accessible from* ***r***(*t*) *via* ***κ***(*t*) = ***Nr***(*t*) *and constitute a latent dynamical system since* 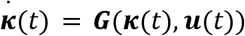 *does not explicitly depend on* ***r***(*t*).
2. ***Universal approximation property:*** *The network has trainable parameters that endow the low-dimensional latent dynamical system with the ability to become a universal approximator in the limit N* → ∞.

These conditions are (non-exhaustively) covered by the four principles we build our framework on:

- **P1** states the encoding map is linear,
- **P2** states the embedding map is nonlinear,
- **P3** states that computations carried in a network can be described by a low-dimensional latent dynamical system connected to neural dynamics via **Definition 1** plus **P1** and **P2**,
- **P4** states that the synaptic parameters describing the connections between neurons constitute the trainable LPU parameters and endow the network with universal capabilities.

Though **Definition 2** would in principle allow nonlinear encoding maps (see **Supplementary Note 1** and **Table S2** for detailed discussions of resulting constraints), linear encoding has several desirable properties as noted in the main text. For instance, as long as the encoding map is linear, the latent dynamical system is guaranteed to be self-sufficient, *i.e*., satisfy the encoding-embedding relationship in **Definition 1** without any additional constraints. Future work could consider extensions of these four principles, *e.g*., by allowing nonlinearity in the encoding map.

#### Universal approximation in a broad class of biologically plausible neural networks

The first three principles described above can be satisfied by a broad class of abstract networks and do not necessarily ground the framework in biology. To render these networks biologically plausible, *i.e*., so that they can be implemented by using well-known biological processes, we turn to the final principle. For ease of discussion, we will omit the inputs and biases when discussing networks with multiple synapses, though these can be reinstated trivially.

As described in Eqs. (3-4), we first start by defining the currents induced at presynaptic terminals of individual neurons:

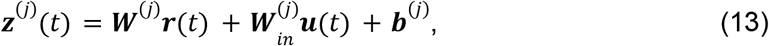

where ***z***^(*j*)^(*t*) is the net induced current at the *j*th synaptic compartment, ***W***^(*j*)^ is the weight matrix connecting presynaptic neurons to this compartment, and vice versa for input weights 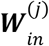 and biases ***b***^(*j*)^. Formally, we assume *j* = 1, …, *L* synaptic compartments, which combine these currents into a final element-wise drive that updates the firing rates following:

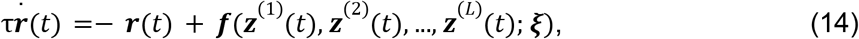

where ***ξ*** denotes any additional parameters not directly affecting presynaptic inputs ***z***^(*j*)^ (*t*). As discussed in Eq. (4), one can define the set of latent variables by performing a low-rank decomposition on the collection of weight matrices, *i.e*., ***W***^(*j*)^ = ***M***^(*j*)^ ***N***^(*j*)^. In this decomposition, there are two possibilities: i) each ***W***^(*j*)^ with rank *K* can support a distinct set of *K*_*j*_ latent variables, defining in total 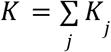 latent variables, or ii) some ***W***^(*j*)^ can share the same encoding weights, hence defining the same latent variables through distinct current contributions and thereby embedding them through distinct nonlinearities. In this work, the discussion below remains virtually the same regardless of the choice. Thus, to remain as general as possible, we will assume case i). Then, the full set of latent variables is given by the group ***κ*** = (***κ***^(1)^, ***κ***^(2)^, …) describing a 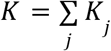 dimensional dynamical system and ***N*** denotes the concatenated collection of all ***N***^(*j*)^ for *j* = 1, …, *L*.

We now formalize one set of plausible and relatively mild assumptions that allow networks described by Eq. (14) to support LPUs:

##### Theorem 1

**Proof**: The latent dynamics follow 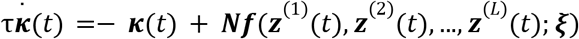 by construction. Let *f*_*_(·) denote an element-wise nonlinearity satisfying the stated conditions and occurring with probability *p*_*_ > 0. As *N* → ∞, the number of neurons with nonlinearity *f*_*_(·) diverges almost surely. By restricting each such neuron to a presynaptic input along which *f* (·) is nonpolynomial (call it 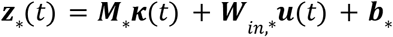, set all other presynaptic inputs to zero and fix ***ξ*** to some value that does not destroy the contribution of ***z***_*_ (*t*)), these neurons form a conventional single-hidden-layer network with a nonpolynomial activation function*f*_*_. The universal approximation theorem for nonpolynomial activations then implies that their weighted outputs can uniformly approximate each component of any continuous vector field ***G***(***κ***(*t*), ***u***(*t*)) on a compact domain to arbitrary accuracy^43^. The remaining neurons and unused synaptic inputs may be assigned zero weights. Therefore, with probability one as *N* → ∞, the induced latent dynamics can approximate ***G*** with arbitrarily small uniform error as long as *K* ≪ *N* remains fixed.

**Theorem 1** is one way through which the biologically plausible networks described by Eq. (3) can achieve universality, and therefore support LPUs. There are several equivalent ways these properties can likely be formalized. The key point is that the nonlinearity is needed to achieve universal approximation and therefore has been stated as a main principle (**P2**). Without this nonlinearity, the latent variables simply follow linear dynamical systems (**Table S2**).

#### Two canonical architectures with desirable properties in the general class of networks

We discuss two key architectures that are part of the class described by Eq. (3) and satisfy the requirements of **Theorem 1**. The first architecture (called basic RNNs) uses one of the simplest embedding functions, *i.e*., a linear function followed by nonpolynomial nonlinearity:

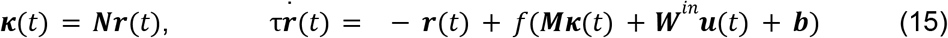

where ***M, N*, W**^*in*^, and ***b*** are learnable parameters as before, τ > 0 is the neuronal time constant, and *f* is an element-wise nonlinearity (taken as *f*(·) = tanh(·)). Defining ***W*** = ***MN*** as the weight matrix reveals the RNN and the latent dynamical system equations:

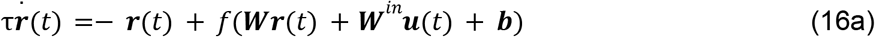

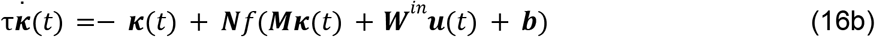

Importantly, Eq. (16b) describes a feedforward neural network with a single hidden layer and therefore is a universal approximator. However, the embedding map described via the equation ***φ***(***κ***(*t*), ***u***(*t*)) = *f*(***Mκ***(*t*) + ***W***^*in*^ ***u***(*t*) + ***b***) is constrained by the choice of the nonlinearity, and therefore does not universally describe the neural tuning properties.

The second architecture (called gain-modulated RNNs) follows the dynamics (cf. Eq. (6)):

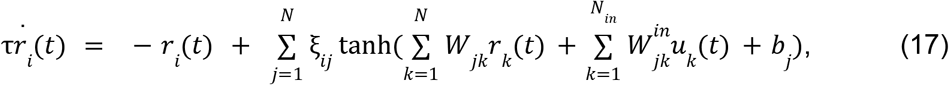

where we can define once again ***W*** = ***MN*** and arrive at the latent dynamics as follows:

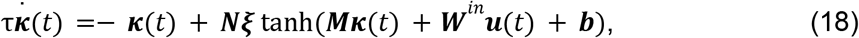

which is equal to Eq. (16b) when ξ_*ij*_ = δ_*ij*_. This particular architecture is more powerful than its basic counterpart, because the embedding map is now also universal:

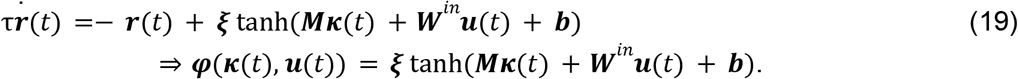

Specifically, ***φ***(***κ***(*t*), ***u***(*t*)) is now represented with a feedforward neural network with a single hidden layer. Note that, as is written in Eqs. (6) and (17), gain-modulated RNNs mix postsynaptic currents between neurons, which is not consistent with Eq. (3) that assumes ***f*** is a collection of element-wise nonlinearities that act locally on presynaptic currents. To resolve this, consider a slightly more complex architecture:

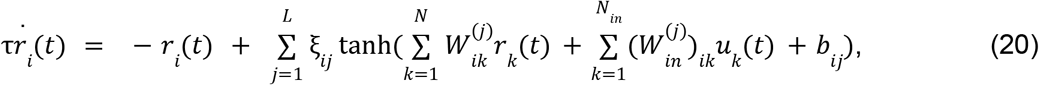

where we identify the variable 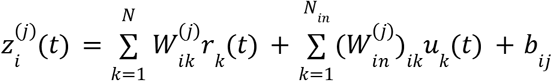 as the net presynaptic current into the *j*th synaptic compartment of the *i*th neuron. Then, the summation 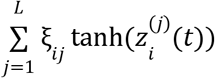 constitutes a local operation at the neuron *i*, which now has a local nonlinearity 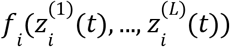. Here, the key is to realize that gain-modulated RNNs constitute a special version of this architecture. There are *N* = *L* compartments and the synaptic parameters are shared across all neurons such that 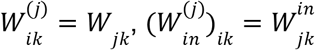, and *b*_*ij*_ = *b*_*ij*_, which turns Eq. (20) into Eq. (17).

We refer to the general formulation in Eq. (20) as a dendritic RNN, which is a step closer to biological realism than gain-modulated RNNs as one may wish to pick *L* ≪ *N* and not have the synaptic parameters be shared across neurons. In this work, we used gain-modulated RNNs for simplicity of exposition and the historical relevance of deep RNNs in machine learning literature.

#### RNNs performing K-bit flip-flop tasks

In K-bit flip-flop tasks (**Fig. 1c**), basic RNNs receive occasional pulses from K independent input channels and must produce corresponding outputs. Each input-output channel operates independently. In any given time bin, each input channel has a 0.05 probability of emitting a pulse with a value of either +1 or −1; otherwise, the input is 0. The network’s task is to remember the sign of the most recent pulse for each channel and output this sign accordingly. With K channels, there are 2^*K*^ possible output states. Each trial starts with a random initialization of the neural activities following a normal distribution with mean 0 and s.d. 1, and continues for 100 discrete time points as input pulses are sampled randomly and independently for each of the K channels. To train the basic RNNs on these tasks, we discretized the time evolution equations as:

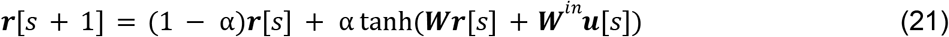

where α = 0. 5 is the discretization ratio (α = Δ*t*/τ for a discretization time Δ*t* = 5ms and τ = 10 ms) and we assumed no biases for simplicity. The RNNs were capable of solving the flip-flop tasks by forming bistable dynamical systems per latent dimension (**Fig. 1**), *i.e*., using odd-valued flow maps with zero biases.

To train low-rank RNNs performing the K-bit flip-flop tasks, we utilized the two-step training paradigm from prior work^33^. First, we trained full-rank RNNs to perform the K-bit flip-flop tasks, for which we re-used the publicly available code from Ref. 102. We employed Adam optimization with a learning rate of 1/*N*, and trained for 5000 epochs with a batch size of 50 trials. We used the mean squared error of the output as the loss function. We enforced *W*_*ii*_ = 0 for the training of full-rank RNNs by projecting the gradients at each epoch. We also used a learning rate scheduler, which reduced the learning rate if it plateaued for 50 epochs. All weight matrices were initialized using Xavier initialization.

Second, we constrained low-rank (K-rank) RNNs to reproduce the neural activities of these full-rank RNNs. To train the low-rank RNNs on the activities of the full-rank RNNs, we used a Pytorch model with a low-rank reconstruction for ***W*** = ***MN***, and used the logistic loss^102^. We used Adam optimization with a learning rate of 10^−3^ and weight decay of 10^−7^, and trained for 20000 epochs with a batch of 5000 trials. The resulting networks had weight matrices which were by design rank *K*, the same as the number of inputs.

To break the symmetry introduced by the linear identifiability, we aligned the latent variables such that each rank corresponded to a particular output, *i.e*., κ_*i*_ (*t*) ~ *o*_*i*_ (*t*). To achieve this, once RNNs were fully trained, we first computed the singular value decomposition of the weight matrices and initialized ***M*** with the left eigenvectors and ***N*** with the singular values times the right eigenvectors. Next, we sampled neural trajectories for 4000 data points as RNNs were performing the K-bit flip-flop task and performed a regression between the network outputs and the linearly encoded latent variables. Using the learned regression matrix ***S***, we transformed ***N*** → ***SN*** and ***M*** → ***MS***^−1^ such that the newly defined latent variables were aligned with the outputs, *i.e*., ***SNκ*** ~ ***o***(*t*). The resulting encoding and embedding weights, which satisfy ***MS***^−1^***SN*** = ***MN*** = ***W***, were used in Eq. (16) to draw and analyze the LPUs.

To compute the tuning curves, ***r***(***κ***), with respect to ***κ*** in these RNNs, we considered the steady-state response (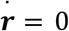, **Supplementary Note 1**) after fixing ***κ***(*t*): = ***κ*** indefinitely, leading to:

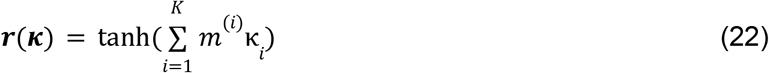

As long as the behavioral timescales that ***κ***(*t*) operates at are significantly larger than τ (*e.g*., fixed-point attractors lead to fixed ***κ*** values in the K-bit flip-flop task), Eq. (22) defines the tuning curves for the neurons.

We used basic RNNs trained on flip-flop tasks in **Figs. 1, 5, 6**, and **S1**. For illustrations in **Fig. 1c-e**, we trained a representative rank-2 RNN solving the 2-bit flip-flop task using the two-step training paradigm described above. The RNN had 100 neurons, a timescale τ = 10*ms*, and was discretized with Δ*t* = 5*ms*. The neural activities in **Fig. 1c** are collected for 1,000 time points, whereas for **Fig. 1d-e**, they were collected for 10,000 time points. For **Fig. 5**, we used the same RNN used in **Fig. 1c-e**. For **Fig. 6c**, each neuron was assigned to one of the 27 coding groups. We defined coding for a particular flip-flop bit by thresholding the corresponding embedding weights, ***m***^(*i*)^, by ± 0. 5. For instance, if 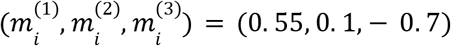, this neuron was assigned to the group (1, 0, −1). For **Fig. S1b**, we trained full-rank RNNs using the first step of the training paradigm discussed above, without the learning rate schedulers.

#### Gain-modulated RNNs performing 2-bit flip-flop tasks

For **Figure 1f-h**, we trained the gain-modulated RNNs to perform 2-bit flip-flop tasks. The networks contained *N* = 100 neurons and equal number of hidden units, with *K* = 2 latent variables defined as ***κ***[*s*] = ***Nr***[*s*]. Their discretized dynamics were:

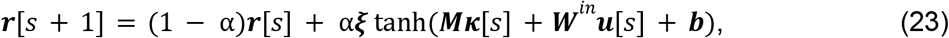

where α = 1 and the latent dynamics followed ***κ***[*s* + 1] = ***Nξ*** tanh(***Mκ***[*s*] + ***W***^*in*^ ***u***[*s*] + ***b***). We trained the networks by minimizing the mean squared error loss between the latent outputs and the target outputs using Adam optimization with a learning rate of 10^−3^. We clipped the gradient at norm 1 and trained for 500 epochs. Training batches contained 100 trials of 300 time points, with each input channel emitting a pulse with probability 0.1. We enforced a minimum interval of four time points between successive pulses. Neural activities were initialized at zero.

### Network timescales with latent variables

Here, we first state the formal assumptions underlying **Theorem 2** and prove it. Then, we discuss the empirical details of the bistable dynamical system analysis in **Figure 2** and **Supplementary Note 2**. We conclude this section by establishing this example’s connection to **Theorem 2**.

#### Computations in extended timescales

As an important consequence of **Theorem 1**, latent computation does not need to unfold at the timescales set by the neuronal time constants. However, for a network with a finite number of neurons, errors occur in implementing a target computation at a target timescale. Here, we state the formal assumptions leading to **Theorem 2** that quantifies these errors.

The zeroth assumption is not so much a formal assumption as a choice of basis and relies on the linear identifiability of latent variables. (This notion simply refers to the scale-equivariant definition of latent variables cf. Eq. (4) that keeps the physical network operation invariant. As an example, one can keep the synaptic connection ***W*** constant and achieve the exact same network dynamics by re-defining ***M*** → *a****M*** and ***N*** → *a*^−1^ ***N*** for any scalar *a* > 0. See the details in the next section.) We simply assume that the latent basis is chosen such that ***κ*** remains (locally) *O*(1), *i.e*., *N* and β independent, within the latent subspace that supports the computation. This assumption is in line with the notion that we expect the nonlinear embedding ***φ*** in Eq. (2) to be an intrinsic neuronal property and how different neuronal outputs are combined as in Eq. (5) should in principle give rise to the network timescales. Relatedly, this choice of basis is also implicit in Eq. (8), where we use ***h***(***κ***(*t*)) to describe a computation in unitless time and temporal information is described by the effective timescale τ_*eff*_. The bistable dynamical system in **Figure 2** presents one plausible example where the construction in Eq. (8) is possible within a compact subspace (see below).

The first assumption is a self-averaging assumption for the universal approximator. Biologically, this assumption corresponds to the notion that synapses between neurons scale *W*_*ij*_ ~ *O*(*N*^−1^), which has been the primary assumption of several prior works on low-rank RNNs^34,35,96^. To formalize this assumption, for convenience of notation, we first define the *p*th latent variable as 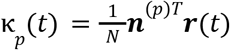, where 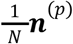 is the encoding weight that corresponds to a row vector of the encoding matrix ***N*** in Eq. (5). Then, the universal approximator in Eq. (7) becomes:

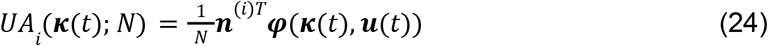

The assumption is as follows: each entry of ***n***^(*p*)^ for *p* = 1, …, *K* and ***φ***(***κ***(*t*), ***u***(*t*)) for the compact subspace (***κ, u***) that supports the latent computation is *O*(1), *i.e*., *N*-independent. Formally, we say that each entry (a neuron constitutes one entry) is sampled iid from a probability distribution governing the joint random variables (*n*^(1)^, *n*^(2)^, …, *n*^(*K*)^, φ(***κ***(*t*), ***u***(*t*))), and this probability distribution has bounded covariance terms. With these assumptions, as *N* → ∞, the law of large numbers ensures that ***UA***(***κ***; *N*) converges to its mean ***UA***(***κ***) with vanishing variance:

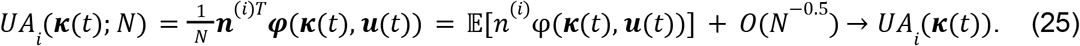

Eq. (25) is what is precisely meant when we assume a self-averaging latent dynamical system.

The second assumption is about spontaneous changes in synaptic parameters, *e.g*., failures of synapses. In the language of the previous assumption statement, this assumption is that the random variable (*n*^(1)^, *n*^(2)^, …, *n*^(*K*)^, φ(***κ***(*t*), ***u***(*t*))) has inherent variability, *i.e*., has finite covariance terms that are independent of β. One biologically interesting possibility is that a neuron may fail to fire spontaneously, giving φ = 0 instead of its expected *O*(1) value. Another one is that a synapse may fail and give 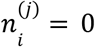, or a value that is off by its expectation by an *O*(1) value. Combined with the first assumption, write can formally write down both assumptions jointly as 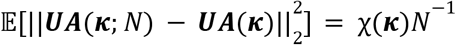 for the compact subspace (***κ, u***) that supports the latent computation, where 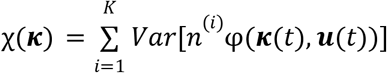 does not depend on β to the leading order. In short, the first assumption ensures that ***UA***(***κ***; *N*) converges in probability to its mean ***UA***(***κ***) as *N* → ∞, whereas the second assumption formalizes the inherent nature of the variability in synaptic parameters and therefore the deviations remain β*-*independent in the large but finite *N* and β.

We now state **Theorem 2** below for completeness and prove it:

##### Theorem 2

*(Computation over extended timescales). Let an LPU implement the function in Eq. (8) for a finite N using an approximator* ***UA***(***κ***; *N*) *that is self-averaging. Further, let the synaptic parameters have inherent variability that is independent of* β. *Then, the estimator* ***h***(***κ***; *N*) = β(***UA***(***κ***; *N*) − ***κ***) + ***κ*** *has an error scaling as* 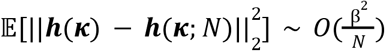, *where the expectation is taken over the synaptic parameter distribution for a given* ***κ***.

**Proof**: The proof follows directly from the two assumptions. First, we note the identity

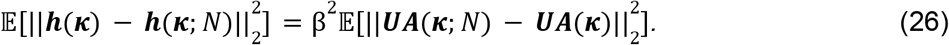

By self-averaging (assumption 1), each component 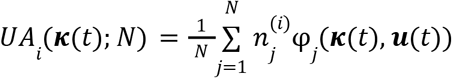 is a sample mean of iid per-neuron contributions and unbiased: E[*UA*_*i*_ (***κ***(*t*); *N*)] = *UA*_*i*_ (***κ***(*t*)). The variance is then given by:

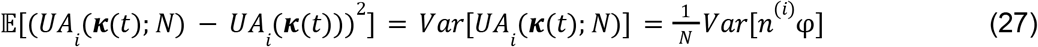

Summing over *K* components, 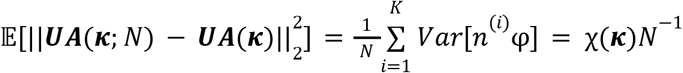, leads to a value that does not depend on β to the leading order with χ(***κ***) ~ *O*(1) (assumption 2). Substituting this value into Eq. (26), we arrive at the error 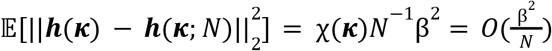.

#### Designing latent processing units with bistable dynamical systems

We designed the one-dimensional LPUs in **Fig. 2** by sampling the encoding and embedding weights for each neuron, (*m*_*i*_, *n*_*i*_), from a zero-mean two-dimensional Gaussian distribution with a positive correlation. Using a mean-field approach (**Supplementary Note 2**), in the limit *N* → ∞, the LPUs follow the equation:

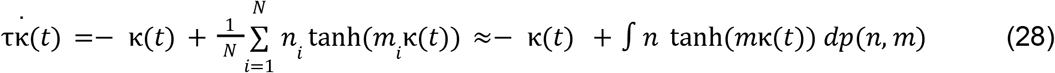

Where *p*(*n, m*) is the probability density function for the pair of random variables (*m, n*) and the right-hand side corresponds to the expectation over these variables. As shown in **Supplementary Note 2**, the target effective timescale is given by β = |*Cov*[*n, m*] ™ 1^−1^.

For **Figure 2**, we sampled the embedding and encoding weights from a zero-mean Gaussian distribution. We picked the parameters describing the (*m, n*) random variable pair to match the target timescales. Specifically, for **Figure 2b, d**, we used the standard deviations σ_*m*_ = 0. 1, σ_*n*_ = 14. 428, and the correlation ρ = 0. 7. Taking τ = 1*ms*, these values give β = |0. 1 × 14. 428 × 0. 7 − 1|^−1^ ≈ 100 and τ_*eff*_ ≈ 100*ms*. For **Figure 2c**, we used σ_*m*_ = 1. 38, σ_*n*_ = 2. 07, and ρ = 0. 7, giving β = |1. 38 × 2. 07 × 0. 7 − 1|^−1^ ≈ 1 and τ_*eff*_ ≈ 1*ms*. For **Figure 2e,f**, we design the networks by first defining the target τ_*eff*_ and consequently β value. Then, we set ρ = 0. 9, which jointly with the target constrains the product σ_*m*_ σ_*n*_. Since only the product matters for this analysis, we pick them equal such that σ_*n*_ = σ_*m*_ and sample the networks.

#### The connection between the bistable dynamical system and Theorem 2

To establish the equivalence of this example to the assumptions of **Theorem 2**, consider the linearization around the origin. Linearizing Eq. (28) around the origin leads to:

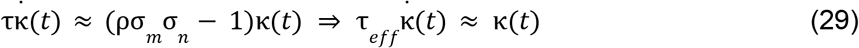

Comparing this equation to Eq. (8) leads to *h*(κ(*t*)) = 2κ(*t*), which does not depend on β. Moreover, cf. our discussion about the scale-equivariance of κ(*t*), the dynamics in Eq. (29) around the origin (the compact subspace of interest) is scale-invariant. Hence, around the origin, this network indeed performs a computation over an extended timescale β = τ_*eff*_ /τ. Now, since (*m, n*) are sampled from a joint Gaussian distribution, the self-averaging follows directly from Eq. (28). The second assumption can be tested by computing the following quantity around the origin (and assuming *Cov*[*n, m*] > 1 due to bistability):

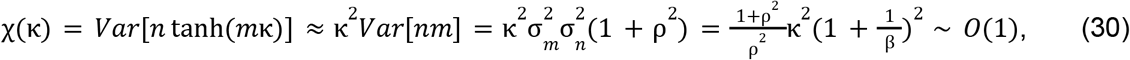

which means the synaptic variability has a leading β-independent term for β ≫ 1. Hence, the bistable dynamical system we consider in **Fig. 2** is an example of **Theorem 2**, which gives:

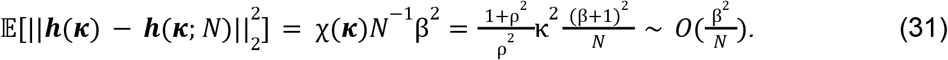

### Decoding universal outputs from the latent processing units

While LPUs are capable of universally approximating latent dynamical systems, the results of these dynamical computations should also be accessible to downstream units, *e.g*., motor neurons. Below, we lay out a theoretical foundation for studying this problem and prove that linearly encoded LPUs allow differentiable functions of latent variables to be accessible via linear readouts from neural activities. First, with the condition of increased dimensionality, we prove that LPUs allow linear readout of any smooth function of their state variables. Next, we illustrate that these readouts can be performed directly on neural activities, mitigating the commonly observed identifiability issues of latent variables^14^. Then, we combine both results into a universal decoding theorem (**Theorem 3**), which suggests that LPUs can subserve complex network output. Finally, we conclude with the details of the empirical analyses performed in **Figs. 3** and **S2**.

#### Sufficiency of linear readouts from extended latent processing units

Consider a K-dimensional LPU with *B* outputs defined as differentiable functions of its latent variables. Below, we formalize the possibility to design an extended LPU with *K* + *B* dimensions that can provide the same *B* outputs with its latent variables:

##### Lemma 1

*(Extended LPUs). Let* ***r***(*t*) ∈ ℝ^*N*^ *be activities in a network that can support LPUs. Let* 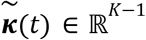 *constitute an LPU with the time evolution* 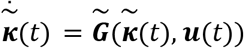. *Let the output, o*(*t*) ∈ ℝ, *of this system be defined as* 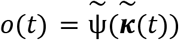 *for some, potentially nonlinear, continuously differentiable function* 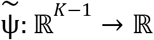. *Then, it is possible to design an extended LPU with dynamics* 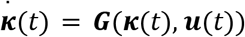 *such that* 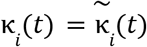 *for i* = 1, …, *K* − 1 *and* κ_*K*_ (*t*) = *o*(*t*) + *c for some constant c* ∈ ℝ.

**Proof:** Let us consider the time derivative of the output in the original LPU:

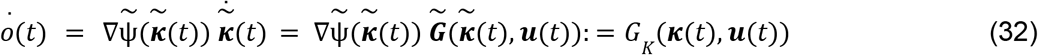

which is a function of ***κ***(*t*) and ***u***(*t*) once we define 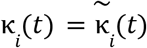 for *i* = 1, …, *K* − 1. This is a self-sustained dynamical system equation for the *K*th component of the new LPU. Similarly, we can also define 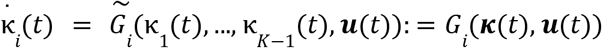 for *i* = 1, …, *K* − 1 making all *K* components of the new LPU evolve in a self-contained manner, *i.e*., 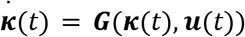. Since κ_*K*_ (*t*) evolves with the same dynamics as *o*(*t*), they can differ only by a constant. This concludes the proof.

One can repetitively apply the process described in **Lemma 1** to accommodate *B* outputs, though at the expense of increased latent dimensionality. However, this lemma has two caveats:

1. **Lemma 1** does not guarantee that the neural tuning properties with respect to the original latent variables will remain unchanged after the re-definition. Similarly, the weight matrix that supports the computation before and after the extension does not necessarily remain invariant. Instead, **Lemma 1** suggests that the same group of neurons can be repurposed, perhaps with a different set of parameters, to support the extended LPU as long as *N* ≪ *K*.
2. As a second caveat, extending the latent system introduces one additional initial condition κ_*K*_ (0) = *o*(0) that needs to be specified alongside 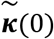 so that the constant in **Lemma 1** becomes *c* = 0. In the most general case, there are *N* initial values, ***r***(0), that can be chosen to satisfy the *K* initial conditions for ***κ***(0). But, a more biologically realistic treatment should likely focus on ***r***(0) values that are randomly sampled. For instance, if neural activities ***r***(0) are randomly initialized from an independent zero-mean distribution, then for each latent variable, we have the relationship: 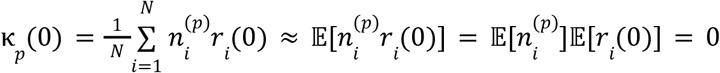. In other words, all latent variables, including κ_*K*_ (0), would be initialized approximately at the origin. In our work, we focus on dynamical processes and the behavior of randomly initialized networks processing temporal information. We believe a detailed treatment of initial condition design, such as how context or network state might cue stable initial conditions, constitutes viable research directions. Finally, **Lemma 1** has a corollary that is relevant for biological computation with LPUs:

##### Corollary 1

*(Linear decoding). An output defined as part of the LPU, i.e*. 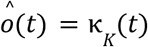, *can be directly and linearly decoded from neural activities*, ***r***(*t*).

**Proof:** Since κ_*K*_ (*t*) is linearly encoded, one can define the output as 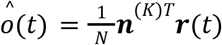. Here, 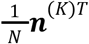 is defined as the *K*th row vector of the encoding matrix ***N***.

The extension of dimensionality in **Lemma 1** may seem like a restriction. However, since we assume *K* ≪ *N*, increases in latent dimensionality at the scale *B* ~ *O*(*K*) do not change the scaling relationships, and may even be desirable for robust operation, *e.g*., to allow feedback from the output to affect the latent variable operation. It is worth highlighting that **Corollary 1** does not guarantee that an output from the LPU will be linearly accessible; only that they could be. A larger (yet equivalent) dynamical system can be designed that contains the same latent variables as a subsystem, but also allows linear readout. By using slightly more resources, biological and/or artificial networks can, but not necessarily have to, allow linear readouts from their neural activities, and consequently from the latent variables of the LPUs.

#### Linear readouts from LPUs without explicit identification

**Corollary 1** suggests that the output, if linearly encoded into a latent variable, can be accessed with a linear readout from neural activities. However, latent variables have what is called a *GL*_*K*_ (ℝ) symmetry. Specifically, if ***κ***(*t*) constitute the latent variables of an LPU, so would 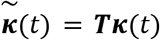, where ***T*** ∈ ℝ^*K*×*K*^ is an invertible matrix. This property is known as linear identifiability. For our purposes, this property means that assigning the behavioral output to a particular variable would break the generality of our mathematical treatment, which should respect this symmetry.

To prevent losing generality, instead of enforcing κ_*K*_ (*t*): = *o*(*t*), we define the output as a linear readout from LPUs as 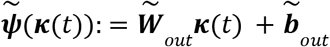 for some parameters 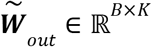 and 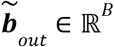. This definition respects the *GL*_*K*_(ℝ) symmetry, since the transformation on the latent variables would still lead to another linear readout, *i.e*., a re-definition of 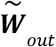. But, which particular collection of ***κ***(*t*) would be most relevant to explicitly keep track of? The following lemma suggests that 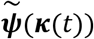 can be computed directly from ***r***(*t*), *i.e*., the behavioral readout can be achieved without having to identify any particular instantiation of the latent variables, ***κ***(*t*).

***Lemma 2*** *(Equivalence of decoding from neural and latent activities). Let* ***r***(*t*) ∈ ℝ^*N*^ *represent neural activities and* ***κ***(*t*) ∈ ℝ^*K*^ *be the corresponding latent variables obtained via a linear encoding. Let* ***κ***(*t*) = ***Nr***(*t*) *for* ***N*** ∈ ℝ^*K*×*N*^. *Let* ***P***_***N***_ *be the projection operator onto the row space of* ***N***, *and* 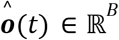 *denote the behavioral readouts from the network with N* ≫ *K* ≥ *B. We define a linear decoder on neural activities as* ***ψ***(***r***(*t*)) = ***W***_*out*_ ***r***(*t*) + ***b***_*out*_, *where* ***W***_*out*_ ∈ ℝ^*B*×*N*^ *and* ***b***_*out*_ ∈ ℝ^*B*^ *are trainable parameters. Then, the following statements are true:*

1. *For any linear latent decoder* 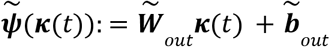 *with* 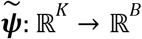, *there is at least one equivalent linear decoder* ***ψ***(***r***(*t*)) *such that* 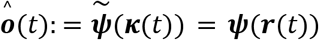.
2. *Conversely, for a linear decoder* ***ψ***(***r***(*t*)) *satisfying the relationship* ***W***_*out*_ (1 − ***P***_***N***_) = 0, *which is both sufficient and necessary, there is at least one equivalent linear latent decoder such that* 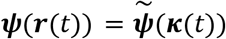.

**Proof:** By simply replacing ***κ***(*t*) = ***Nr***(*t*), we prove the first statement of the theorem following the equality 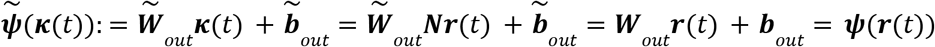, where the parameters of the latter decoder relate to the former as 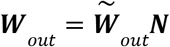 and 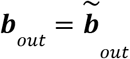. To prove the converse direction, we start with:

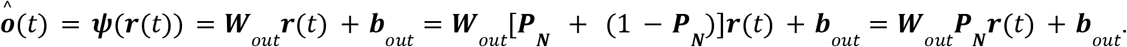

Here, we have used the assumption that ***W***_*out*_ (1 − ***P***_***N***_) = 0. Next, we can explicitly write the projection matrix as ***P***_***N***_ = ***N***^*T*^ (***NN***^*T*^)^−1^ ***N*** such that the behavioral readout becomes:

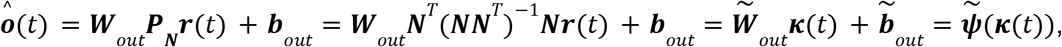

where we identify 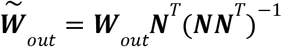 and 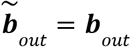. Thus, ***W***_*out*_ (1 − ***P***_***N***_) = 0 is a sufficient condition to define an appropriate 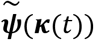.

Next, we prove the necessity of ***W***_*out*_ (1 − ***P***_***N***_) = 0 using contradiction. Suppose that this condition does not hold such that the linear readout can be written as 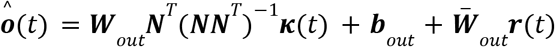, where we define 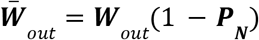. The presumption that 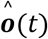 is a function of ***κ***(*t*) means that for two distinct ***r***_±_ (*t*) corresponding to the same ***κ***(*t*) value, the output 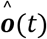 should be the same. Now, consider a general ***r***(*t*) for which (1 − ***P***_***N***_)***r***(*t*) ≠ 0. Define ***r***_±_ (*t*) = ***P***_***N***_ ***r***(*t*) ± (1 − ***P***_***N***_)***r***(*t*). Both map to the same ***κ***(*t*) since ***κ***(*t*) = ***Nr***_±_ (*t*) = ***NP***_***N***_ ***r***(*t*). However, they produce two distinct outputs following 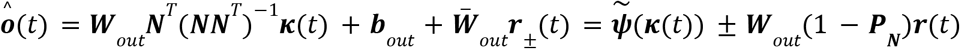. Here, the second term can be made non-zero by choosing ***r***(*t*) such that ***W***_*out*_ (1 − ***P***_***N***_)***r***(*t*) ≠ 0. This can be done because ***W***_*out*_ (1 − ***P***_***N***_) ≠ 0 by construction. Hence, ***o***(*t*) achieves two distinct values for the same ***κ***(*t*), which is a contradiction to its function nature. This proves the second statement, concluding the proof.

One might wonder whether the linear bottleneck condition, *i.e*., ***W***_*out*_ (1 − ***P***_***N***_) = 0, is too restricting. Similar to **Lemma 1**, this readout too can be incorporated into the LPU dynamics. Specifically, if ***W***_*out*_ (1 − ***P***_***N***_) ≠ 0 and for simplicity assume *B* = 1, we can define 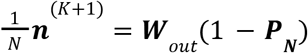 and subsequently 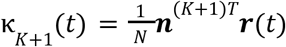. Then, by inspection, 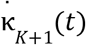 depends only on (κ_1_, …, κ_*K*_), as 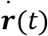 was defined following a (nonlinear) embedding equation of the K-dimensional LPU. Thus, similar to our calculations in **Lemma 1**, it is possible to extend the LPU.

#### Linear invariance of the causally coding subspace

Returning to the identifiability concerns, if ***P***_***N***_ were to transform with different bases of latent variables, enforcing ***W***_*out*_ (1 − ***P***_***N***_) = 0 might require biological neural networks to explicitly encode and utilize a preferred basis of latent variables. Fortunately, as we show below, this is not the case, *i.e*., ***P***_***N***_ is invariant under *GL*_*K*_(ℝ) transformations. Defining the *K*-dimensional subspace defined by the projection operator ***P***_***N***_ as the causally coding subspace, we have the following lemma:

##### Lemma 3

*(Invariance of the causally coding subspace). Let* ***r***(*t*) ∈ ℝ^*N*^ *represent neural activities encoding* ***κ***(*t*) = ***Nr***(*t*) *for* ***N*** ∈ ℝ^*K*×*N*^. *Let* ***P***_***N***_ *be the projection operator onto the row space of* ***N***. *Define an invertible matrix from GL*_*K*_ (ℝ) *group*, ***S***, *that defines a new set of latent variables following* 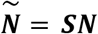. *Then, the causally coding subspace is invariant under the transformations defined by the GL*_*K*_ (ℝ) *symmetry of the latent variables, i.e*.,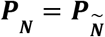.

**Proof:** The proof follows from the explicit calculation of 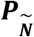 and using the property that ***S*** is an invertible matrix: 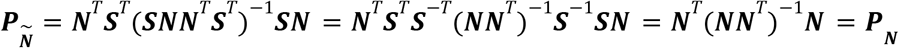. Here, we used the fact that both matrices ***NN***^*T*^ and ***S*** are invertible. The latter is true by definition of *GL*_*K*_ (ℝ), whereas the former is true as long as ***N*** is rank K (as assumed throughout this work).

Here, it is worth noting that we defined the latent variables following 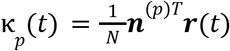, where 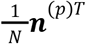 is the *p*th column vector of ***N***. Combining this with **Lemma 3** suggests that the symmetry transformation (***S***) changes the coordinate system that defines the variable basis ***κ***(*t*), but keeps the causally coding subspace invariant. Hence, ***W***_*out*_ (1 − ***P***_***N***_) = 0 condition respects the *GL*_*K*_ (ℝ) symmetry, *i.e*., does not constrain the network to choose a particular biological basis for ***κ***(*t*). If ***W***_*out*_ (1 − ***P***_***N***_) = 0 is true for one basis of latent variables, it is true for all bases.

#### Optimality of linear LPU readouts

Now, we return to the question of linear identifiability of ***κ***(*t*), specifically to the question: Should a network keep explicit track of a particular coordinate system for ***κ***(*t*), or is it sufficient to keep track of the causally coding subspace itself? Fortunately, combining the results of **Lemmas 1, 2**, and **3** enables to prove a joint theorem on the identifiability of LPUs and the universality of their outputs:

##### Theorem 3

**Proof: Theorem 1** ensures that, in the limit *N* → ∞, the LPU can approximate the latent dynamical system of interest with vanishing approximation errors as long as its dimensionality is fixed with respect to *N*. Applying **Lemma 1** repetitively (*B* many times), one can design an extended LPU in which a particular set of *B* outputs from the target dynamical system can be readout linearly from the remaining *B* latent dimensions. Hence, it is possible to design a *K* + *B*-dimensional LPU that models the *K*-dimensional target dynamical system and provides *B* linear outputs from the latent variables. Finally, **Lemmas 2** and **3** jointly ensure that any linear readout from the LPU can be achieved directly from the neural activities, which does not require explicit identification of a basis for the latent variables.

The true power of **Theorem 3** lies in its agnosticism to individual neural processes. As long as the overall nonlinearities are expressive enough, *e.g*., as in the case for basic RNNs above, (extended) LPUs can store desired outputs of the computation in a linearly accessible manner. **Lemmas 2** and **3** jointly suggest that as long as the behavioral readouts are achieved with linear decoding from LPUs, there is no reason for the downstream units (*e.g*., a motor neuron) to explicitly keep track of latent variables in a particular coordinate system.

#### Connections to existing universal approximation theorems

It is worth noting that the universal approximation property we discuss in this work follows directly from the universal approximation theorem for feedforward neural networks^43^. This theorem later inspired Schaefer and Zimmermann’s proof^42^, which demonstrated that discrete recurrent neural networks (RNNs) could universally approximate open dynamical systems. Their proof utilized two distinct neural populations, one implementing the latent dynamics and another handling the readout function. Subsequently, Ref. 35 advanced the field by extending these concepts to another RNN architecture, in which the parameters are sampled using mixtures of Gaussians.

Jointly, our **Theorems 1, 2**, and **3** advance this framework in three significant ways. First, **Theorem 1** establishes the universal approximation properties for a broad class of biologically plausible neural networks, with a careful consideration of increased latent dimensionality and identifiability issues carried through in **Lemmas 1, 2**, and **3**. Second, **Theorem 2** explicitly quantifies the finite-size errors when computation needs to occur in extended network timescales. Finally, when discussing **Theorem 3**, by assuming that the (nonlinear) readout is continuously differentiable, an optimal linear readout can be designed using the same group of neurons as the ones supporting the latent dynamical system, extending the original proof in Ref. 42 that assumed a distinct group of neurons for the readout. One other key point is that none of our theorems are bound to a particular architecture, in line with the rich diversity in biological neural networks.

#### Analysis of the neural recordings from the neocortex

In this work, we re-used our cortex-wide neural imaging datasets from Ref. 8. Thus, the procedures for obtaining the Ca^2+^ traces, including the subsections below from mouse preparation to cell extractions, are the same. For completeness, we summarize the mouse preparation below.

All animal procedures were approved by the Stanford University Administrative Panel on Laboratory Animal Care. For the imaging studies of layer 2/3 neocortical pyramidal neurons, 4 male and 2 female triple transgenic GCaMP6f-tTA-dCre mice were used. Surgeries were performed on mice aged 10–16 weeks under isoflurane anesthesia in a stereotaxic frame. To minimize inflammation and pain, preoperative carprofen (5 mg/kg) was administered subcutaneously and continued for 3 days post-surgery. We created a cranial window (5-mm diameter) over the right cortical area V1 and the surrounding cortical tissue. We covered the exposed cortical surface with a glass coverslip (#1 thickness, 64-0700, CS-5R, Warner Instruments) attached to a steel annulus (1 mm thick, 5 mm *out*er diameter, 4.5 mm inner diameter, 50415K22, McMaster) and secured it with ultraviolet-light curable cyanoacrylate glue (Loctite 4305). For head-fixation during imaging, we cemented a metal head plate to the skull using dental acrylic. In vivo imaging sessions commenced at least 7 days post-surgery.

We recorded Ca^2+^ neural activity videos (20 fps; 2048 × 2048 pixels) using a fluorescence microscope with 40-160 μW/mm–2 illumination. Custom Matlab software controlled visual stimulus presentation, operated the behavioral apparatus via a NI-USB 6008 card, and initiated video capture on the microscope. Post-acquisition, we downsampled the videos to 1024 × 1024 pixels and 10 fps. Lateral brain movements were corrected using Turboreg software for image alignment. Using gaussian spatial high-pass filtering (σ = 80µ*m*), we removed scattered fluorescence and background activity. We then calculated the relative fluorescence changes by computing Δ*F*(*t*)/*F*_0_, where *F*_0_ is the mean activity of each pixel over the entire session and Δ*F*(*t*) is the mean subtracted activity of each pixel at time *t*. Maximum projection images of each session’s Δ*F*(*t*)/*F*_0_ movies were analyzed to quantify lateral spatial displacements. We used *imregtform* in Matlab to determine optimal transformations between the first session’s maximum projection image and subsequent sessions, aligning all videos accordingly. Aligned Δ*F*(*t*)/*F* _0_ videos were concatenated for individual cell extraction and Ca^2+^ activity trace analysis.

We applied principal and independent component analyses (PCA/ICA^103^) to extract individual neuron activity from concatenated Δ*F*(*t*)/*F*_0_ movies. Each mouse’s preprocessed Ca^2+^ video, approximately 1 TB in size, was divided into 16 tiles, each covering ab*out* 1 mm × 1 mm. We ran PCA/ICA in parallel on 16 computing nodes (1 tile per node) and identified neuron Ca^2+^ activity traces and spatial filters. We then isolated cell somas by thresholding each spatial filter at 4 standard deviations of noise fluctuations and replacing filter weights below this threshold with zeros. These spatial filters were reapplied to the Δ*F*(*t*)/*F*_0_ movie to obtain Ca^2+^ activity traces.

For 3 of the 16 image tiles per mouse, individual neurons were manually identified based on their morphology and Ca^2+^ transient waveforms. For the remaining 13 tiles, we trained three binary classifiers (Support Vector Machine, Linear Generalized Model, and Neural Network) using manually identified cells as training data. To train the classifiers we used a set of 12 predefined features to characterize a neuron’s morphology (eccentricity; diameter; area; orientation; perimeter; and solidity) and Ca^2+^ activity waveform (mean peak amplitude of Ca^2+^ transients; signal-to-noise ratio between Ca^2+^ transients and baseline fluctuations; number of Ca^2+^ transients peaks above 3 s.d. of the baseline; number of Ca^2+^ transients peaks above 1 s.d. of the baseline; the difference of the mean decay and mean rise times of the Ca^2+^ transients, normalized by the sum of these two values; and the FWHM of the average Ca^2+^ transient). Classifiers were then used to identify cells in the 13 remaining tiles through a majority vote approach. Manual inspection verified that all cells identified met the criteria for neuron classification.

As described in the main text, we used data from six mice performing a visual discrimination task. In this task (**Fig. 3a**), the animal was shown horizontal or vertical moving gratings as stimuli for 2s. After a short delay of 0.5s, the mice reported the type of the trial by either licking a sp*out* or abstaining (Go-NoGo). For this work, when training our shared decoders, we concatenated the correctly performed trials across multiple imaging days for each animal. Since Ca^2+^ traces collected from the one-photon mesoscope across days can have arbitrary units, we normalized the Ca^2+^ activity traces to have unit power on each day. In total, we designed a massive dataset containing six mice across a total of 30 imaging days (5, 5, 5, 5, 3, 7) and 7778 correctly performed trials (1211, 998, 895, 1812, 1041, 1821). The dataset contained a total of 21570 layer 2/3 pyramidal neurons (5292, 2761, 2236, 4193, 3334, 3754), collected from eight neocortical regions: 5249 neurons from primary visual cortex (V1; 970, 431, 803, 1085, 850, 1110), 1130 neurons from lateral visual cortex (LV; 144, 166, 14, 223, 120, 463), 1797 neurons from medial visual cortex (MV; 409, 123, 250, 335, 445, 235), 3240 neurons from posterior parietal cortex (PPC; 677, 427, 370, 573, 642, 551), 961 neurons from the auditory cortex (A; 106, 347, 0, 157, 18, 333), 7161 neurons from the somatosensory cortex (S; 2297, 1185, 223, 1717, 736, 1003), 324 neurons from the motor cortex (M; 272, 51, 0, 0, 1, 0), and 1708 neurons from the retrosplenial cortex (RSC; 417, 31, 576, 103, 522, 59). For the experiments comparing linear decoders with nonlinear decoders, we enforced that a brain region should at least have 50 neurons.

For the decoding analysis in **Figs. 3c** and **S2**, we used the partial-least squares regression, linear/quadratic discriminant analysis functions, random forest classifiers, and the (cross-validated) logistic regression implemented in Python’s *scikit-learn* package^104^. For the decoding analysis, we used four distinct windows: stimulus window included the mean of the neural activities inside [0.5,2]s time bins; the delay window [2,2.5]s, the response window [2.5,5.5]s; and the trial end window [5.5,6.5]s. To regularize these methods, we first trained PLS (with respect to trial identity) regularized with *K* dimensions on the data from the first set. Then, using the second set, we trained the linear or nonlinear decoders to compute the optimal decision direction in this dimensionally reduced space. Then, we computed the decoder accuracies on the third set by using the dimensional reduction and decoder weights acquired from the first two sets.

### Latent dynamics on the neural activity manifolds

To study the geometrical properties of the LPUs embedded in high-dimensional neural activities, we start by making a connection between neural manifolds commonly studied in systems neuroscience and the dynamical latent variable models we have been studying (**Theorem 4**). In the same theorem, we claim that *K*-dimensional LPUs can lead to an extensive scaling in the linear dimensionality of the neural activities. To prove this statement, we first show that nonlinear embedding functions can lead to curved neural manifolds and complete the proof with a positive example (**Lemma 4**). Finally, we conclude with the details of our simulation experiments in **Figs. 4, S3, S4**, and **S5**.

#### Emergence of neural manifolds

One consequence of using a dynamical embedding map is that the same latent state, ***κ***(*t*), can result from two distinct neural activity patterns, ***r***_±_(*t*), as we have discussed above. Yet, the separation between neuronal (*O*(*ms*)) and behavioral (*O*(*s*)) timescales^63^ enables defining an approximately static embedding relationship between ***κ***(*t*) and ***r***(*t*). The following theorem shows that this property naturally defines what neuroscientists commonly refer as “neural manifolds”:

##### Theorem 4

*(Neural manifolds). Let* ***r***(*t*) ∈ ℝ^*N*^ *represent neural activities of an LPU, which evolve via* 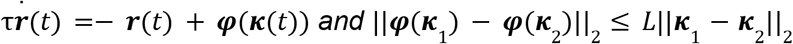 *for some L* > 0. *If the latent variables evolve slowly relative to neural dynamics*, β = τ_*eff*_ /τ ≫ 1, *neural lie near the low-dimensional structure activities* ***r***(***κ***) = ***φ***(***κ***) ***+*** *O*(β^−1^), *whose extrinsic dimensionality can attain the upper bound N*.

**Proof**: We start by solving the equation 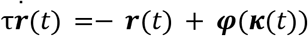 by using an integration factor:

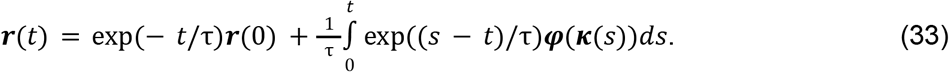

The trick is to write ***φ***(***κ***(*t*)) with the same kernel as

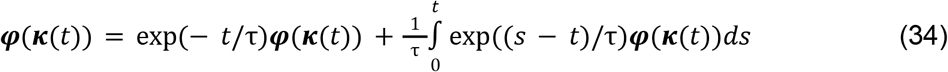

using the identity 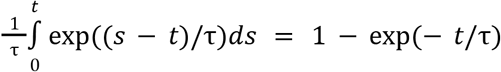. Then, we combine both equations:

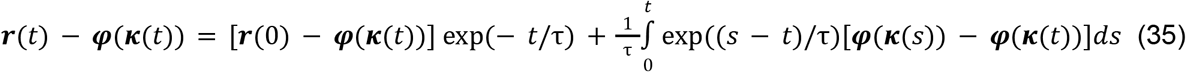

The first term decays to zero at times τ_*eff*_ ≫ τ. To make progress with the second term, we need to bound the difference ||***κ***(*s*) − ***κ***(*t*)||_2_. Fortunately, this bound comes from Eq. (8) and the fact that ***h***(***κ***(*t*)) is continuous and β-independent on a compact subspace of ***κ***, *i.e*., it is also bounded by a constant. Therefore, we can define *M:* = sup || − ***κ + h***(***κ***)||_2_ < ∞ on the same compact subspace and arrive at the intermediary result:

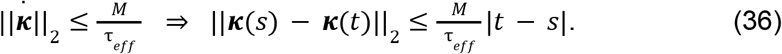

Combined with the Lipschitz inequality in the theorem statement, this bounds the second term in Eq. (35) as:

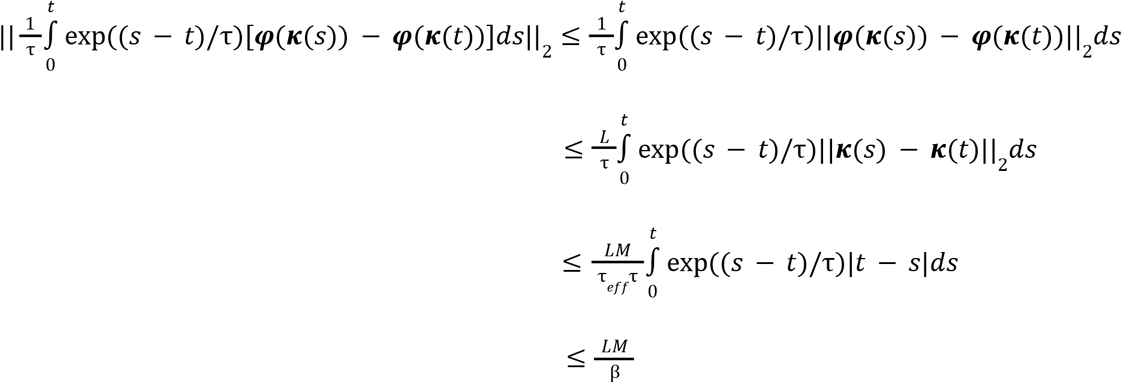

Bringing everything together, we arrive at the final inequality:

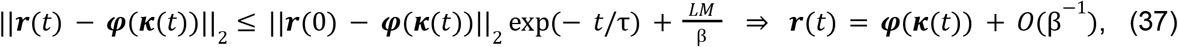

where we have noted that the first term, ~ exp(− β), can be omitted compared to *O*(β^−1^) on the timescale of latent dynamics, *t* ~ τ_*eff*_, concluding the first statement of the proof. To prove the second statement, *i.e*., “*whose extrinsic dimensionality can attain the upper bound N”*, we first introduce some terminology and prove it with a positive example (**Lemma 4**) below.

For now, we briefly note that in the strictest sense, the structure ***r***(***κ***) = ***φ***(***κ***) defined by Theorem 4 is not necessarily a mathematical manifold. Some additional conditions would be needed on both ***κ*** and ***φ***. For instance, if ***κ*** ranges over a compact ball around the origin and ***φ*** is a smooth injective immersion (full-rank Jacobian), the image ***φ***(***κ***) is an embedded submanifold of ℝ^*N*^; weaker conditions also suffice ^105^.

#### Geometrical properties of neural activity manifolds

To prove the statement ab*out* extrinsic dimensionality in **Theorem 4**, we first need precise definitions of the geometric properties of the neural manifolds. We start by defining extrinsic dimensionality, which through*out* this work we use as a synonym for linear dimensionality:

##### Definition 3

*(Extrinsic dimensionality). The extrinsic (or linear) dimensionality of a set of discretized neural activities (r*_i_ [*s*] fo*r s* = 1, …, *T and i* = 1, …, N) *is defined as the smallest dimension d* su*ch that there exists a d*-di*mensional affine subspace containing all the points*.

Extrinsic dimensionality differs from ambient dimensionality, the latter is always *N* for a network with *N* neurons. The former quantifies how many ambient dimensions are actually used by the neural activity traces. With the assumption of sufficient samples, *i.e*., *T* ≫ *N*, the maximum attainable extrinsic dimensionality of a neural manifold is *N*. Both definitions stand in direct contrast to latent (or intrinsic) dimensionality, which we take to be the dimensionality of the latent dynamical system, *i.e*., *K*. We briefly note that more precision would be needed in defining latent versus intrinsic dimensionality if, *e.g*., latent space were not fully utilized. For our purposes, we do not consider these finer distinctions, though note that any such degeneracy only sharpens the separation between extrinsic vs. intrinsic dimensionality.

Since noise and transient responses can artificially inflate the linear dimensionality, in this section, we restrict our analysis to latent variables that are noiseless, slowly varying for timescales much larger than τ, and have negligibly fast transient responses such that the embedding relationship becomes effectively static as shown in **Theorem 4**. Incidentally, as a consequence of these assumptions, the results in this section apply to the traditional framework of deep learning models. With these assumptions, we next define flat and curved manifolds:

##### Definition 4

*(Flat and curved manifolds). If the linear dimensionality exceeds the latent dimensionality, the neural activities are said to lie on a curved manifold. Otherwise, the manifold is flat*.

Here, it is worth noting that we are not interested in the curvature of structures forming in latent or neural activity subspaces as a result of task demands, rather the curvature that stems from the embedding of the former in the latter. While more quantitative definitions are possible (see, *e.g*., Ref. 106), our focus in this work is on a binary distinction between the two classes. The binary nature of the neural manifold curvatures (*i.e*., whether it is zero) plays a central role in our ability to estimate the dimensionality of neural activities. As we discuss in **Supplementary Note 1**, the traditional formulation of low-rank RNNs leads to flat manifolds^11,33–35^. In contrast, as illustrated in **Fig. 4**, basic RNNs have curved neural activity manifolds.

Now, inspired by previous work on counting the piecewise-linear regions of deep neural networks^107,108^, we show that even 1D LPUs can have a maximal extrinsic dimensionality, finishing the proof of **Theorem 4**:

##### Lemma 4

*(Maximal extrinsic dimensionality). Let* ***r***(*t*) ∈ ℝ ^*N*^ *be the neural activities that embed a low-dimensional LPU* κ(t) ∈ ℝ *via a set of static embedding functions, f*_i_ (·), *applied element-wise such that r*_*i*_ (κ) = *f*_*i*_ (*m*_*i*_ κ ***+*** *b*_*i*_) *and m*_*i*_, *b*_*i*_ ∈ ℝ *for i* = 1, …, N. *For a set of functions satisfying the condition f*_*i*_(*x*) = 0 *for x* ≤ 0 *and f*_*i*_(*x*) > 0 *for x* > 0, *it is possible to design a network that attains an extrinsic dimensionality of N*.

**Proof:** We prove this proposition by constructing a positive example. Consider the embedding relationship *r*_*i*_ (κ) = *f*_*i*_ (m_*i*_ κ ***+*** *b*_*i*_) and set all embedding weights to *m*_*i*_ = 1. Then, by defining an element-wise bias 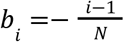 for *i* = 1, …, N, we obtain 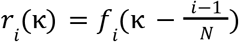. Now, consider the collection of neural activities (***r***^(*j*)^) that result from the embedding 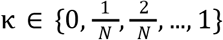 for *j* = 1, …, N ***+*** 1. Then, the affine subspace defined by the points (***r***^(*j*)^) is *N* dimensional. Specifically, for *j* = 1, …, N ***+*** 1 and any given *i* = 1, …, N, the following is true:

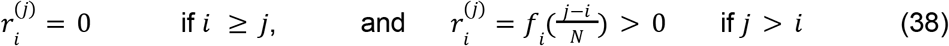

Then, we first note that ***r***^(1)^ = 0, *i.e*., this collection of neural activities contains the origin. Then, we can define the set of vectors spanning this subspace via the matrix 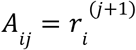 for *i, j* = 1, …, N. Notably, this matrix is upper-triangular with strictly positive diagonal terms, hence full-rank. Therefore, the vectors (***r***^(*j*)^ − ***r***^(1)^) for *j* = 2, …, N ***+*** 1 are linearly independent, *i.e*., span an *N* dimensional subspace.

For simplicity and illustration purposes, **Lemma 4** discusses the case with *K* = 1. Yet, the proof trivially generalizes to higher dimensional LPUs, *e.g*., by separately considering sets of data points sampled alongside a coordinate κ_d_ such that κ_i_ = 0 for *i* ≠ *d*. Finally, **Lemma 4** proves the final statement of **Theorem 4**, finishing the proof.

#### Evaluations of the neural manifold dimensionality

In our simulation experiments, we considered full-rank and low-rank RNNs and defined the linear dimensionality as the number of principal components (PCs) required to explain 99% of the variance in the neural activities.

For the experiments involving static embedding maps (**Figs. 4b-c** and **S3)**, we sampled uniform samples from the K-dimensional unit sphere by first sampling a vector ***κ*** ∈ ℝ^*K*^, in which each element was sampled from standard normal distribution. Then, we normalized the sampled vectors by their norm, leading to samples on the K-dimensional unit sphere. Since a given vector had independent entries, it had rotational symmetry around the origin, leading to uniformly sampled data points. We used several nonlinearities (tanh, sign, ReLu, and sin) and varying levels of embedding strengths to embed the K-dimensional unit spheres into the *N* dimensional neural activity spaces. Specifically, for **Figs. 4b** and **S3a**, we sampled each entry of the embedding weights from a standard normal distribution Ɲ(0, 1) and had 20000 samples per draw. For **Fig. 4c**, we instead sampled from Ɲ(0, 2. 56) and collected 3000 samples per draw. Specifically, to test whether each principal component has information ab*out* the latent variables, we generated two independent sets of 3,000 points on the unit 10-sphere as train and test sets, embedded each through the random sinusoidal nonlinearity, and mean-centered the embedded activities using the training-set mean. We performed PCA on the training activities and a linear regression from the projection onto the *k*th principal component to the latent variables and quantified the prediction accuracy (*R*^2^) on the held-*out* test set.

For the experiments involving the sequence sorting task (**Figs. 4d-e** and **S4**), we trained a set of low-rank RNNs on sequence lengths from 2 to 4. The number of neurons per network ranged from 16, 64, 256 or 1024, whereas the ranks were between 1 and 6. We trained using the Adam optimizer with weight decay set to 10 ^−4^ and a learning rate of 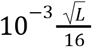, where *L* = 8192 is the batch size. We trained for 10,000 steps with early stopping when the running loss ceased to improve. We had random data draws for each batch and repeated the whole experiment 10 times with different random seeds, leading to 10 distinct networks per condition. We chose Δ*t* = τ and added a random noise sampled from a normal distribution with zero mean and 0.1 s.d. at each time step. Neural activities were initially zero.

To obtain chaotic full-rank RNNs, we randomly initialized RNNs with *N* recurrent units and no external inputs. The entries of the recurrent weight matrix were sampled from a Gaussian distribution 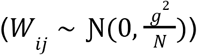, after which the diagonal elements were set to zero 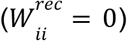. The dynamics of these full-rank RNNs exhibit a transition to chaos at a critical value of *g* = *g*_*crit*_(N). In the mean-field limit^90^, we have *g*_*crit*_ (*N* → ∞) = 1. To obtain the low-rank counterpart, we initialized a full-rank RNN and later applied a rank-reduction operation based on the singular value decomposition and reconstructed a scaled and low-rank version following ***W***_*lr*_ *= std*(***W***_*K*_*)/std*(***W***_*K*_) × ***W***_*K*_. *H*ere, ***W***_*K*_ *is* a reconstruction with the top *K* singular components. The dynamics of these low-rank RNNs exhibited divergent trajectories at a critical value *g* = *g*_*crit*_ *(N*, K) (**Fig. S5**), similar to their full-rank counterpart. For *g* < *g* _*crit*_ *(N*, K), the dynamics decayed to zero, while for *g* > *g*_*crit*_ *(N*, K) there was self-sustained activity in the network, marked by the rapid separation of trajectories with nearby initial conditions (perturbation magnitude 10^−5^).

### Neural coding with and with*out* causal influence on the behavior

As we discussed in **Theorem 1**, when encoding is linear, embedding has to be nonlinear to enable universal approximation of latent dynamical systems. This section discusses the consequences of this nonlinear embedding on the coding properties of neurons. We first prove that any noise added to the neural manifold in the direction that is orthogonal to the causally coding subspace decays exponentially (**Lemma 5**). Second, we show that neural tuning properties can be correlated with latent variables even along directions that have no causal influence on latent computations (**Lemma 6**). These two results combine to give rise to **Theorem 5** in the main text. Finally, we provide the methodological details of the experiments performed in **Figs. 5** and **S6**.

#### Neural activities on and around the neural manifolds

For our calculations in this section, it is instructive to divide the neural activities into two orthogonal components:

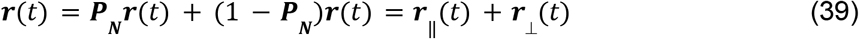

where we recall that ***P***_***N***_ is the projection matrix to the causally coding subspace and is rank *K* by assumption. Here, ***r***_‖_ (t) and ***r***_⊥_(t) are parallel and orthogonal neural activity components to the causally coding subspace, respectively. We first note the following:

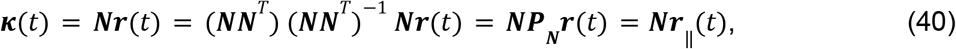

which means that latent variables are defined solely by the parallel activity components. Then, with*out* loss of generality, we can rewrite the arguments of the embedding map in Eq. (2) as:

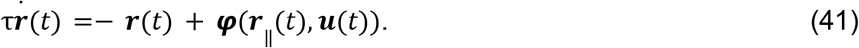

Using this relationship, we now prove the following lemma:

##### Lemma 5

*(Decay of residual activity differences). Let* ***P***_***N***_ be *the projection operator to the\ causally coding subspace, and let* ***r***_1_ (*t*_0_), ***r***_2_(*t*_0_) *be two distinct neural activities encoding the same latent state so that* ***N***r_1/2_ (*t*_0_) = ***κ***(*t*_0_). *If both evolve forward from t*_0_ *under the neural dynamics in Eq. (2), then the latent variables coincide for all t* ≥ t_0_, ***κ***_1_ (*t*) = ***κ***_2_ (*t*), *while the residual difference* Δ***r***(*t*) = ***r***_2_ (*t*) – r_1_(*t*) *decays exponentially at rate* τ^−1^

**Proof:** We start by defining ***r***_‖_(t_0_) = ***P***_***N***_r_1/2_(*t*_0_), which is the same for both initial conditions by Eq. (40). Then, following Eq. (41), we can write the time evolution of ***r***_‖_(t) as:

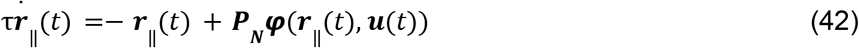

which is a self-contained equation that does not depend on ***r*** _⊥_ (t). Moreover, ***κ***(*t*) = ***N***r_‖_(*t*) is the same for both initial conditions. Consequently, both ***r***_‖_ (t) and ***κ***(*t*) start with the same initial conditions and follow the same equations. Hence, ***κ***_1_ (*t*) = ***κ***_2_(*t*) for *t* ≥ t_0_. In other words, since Eq. (42) does not depend on ***r***_⊥_ (t), the residual difference has no effect on latent dynamics.

Now, to prove the second statement, we first argue that ***P***_***N***_ ***r***_1_ (*t*) = ***P***_***N***_ ***r***_2_ (*t*). This follows directly from Eq. (42) and the equivalence ***r***_‖_ (t_0_) = ***P***_***N***_ ***r***_1/2_ (*t*_0_) we established. Then, we rewrite Δ***r***(*t*) = (***r***_2_ (*t*) – r_1_ (*t*)) = ***P***_***N***_ (***r***_2_ (*t*) – r_1_ (*t*)) ***+*** (1 − ***P***_***N***_)(***r***_2_ (*t*) – r_1_ (*t*)) = (1 − ***P***_***N***_)(***r***_2_ (*t*) − ***r*** _1_(*t*)) as the difference between the two orthogonal components. Noting that ***r***_‖_ (t), hence ***φ***(r _‖_ (t), ***u***(*t*) components, for both initial conditions evolve identically, we can compute the equation governing the difference as 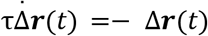. This decays exponentially with a rate τ^−1^, concluding the proof.

#### Neural coding without causal influence on latent computations

**Lemma 5** suggests an interesting observation. While neural activity manifolds can be curved due to nonlinear embedding, deviations off the manifold, lying within the (*N* − *K*)-dimensional subspace defined by the projection operator 1 − ***P***_***N***_, decay exponentially back to the neural activity manifold. A similar observation has been made for the existing low-rank RNN architectures, whose neural activities are constrained to K-dimensional flat manifolds^33^. Unlike the prior architectures, the non-zero extrinsic curvature endows the general class of RNNs defined by Eq.with the possibility to also support redundant information in their tuning properties:

##### Lemma 6

*(Redundancy in neural tuning properties). Let* ***r***(*t*) ∈ ℝ ^*N*^ *be neural activities encoding* ***κ***(*t*) ∈ ℝ^*K*^ *with* ***N*** ∈ ℝ^*K*×*N*^. *Let* ***κ***(*t*) *be slowly varying such that the embedding map becomes approximately static* ***r***(***κ***) ≈ ***φ***(***κ***) *with respect to the latent activities following the setup of* ***Theorem 4***. *Then, neural activities projected to the complement of the causally coding subspace, despite not contributing to the computation, can be tuned to the latent variables*.

**Proof:** The proof follows from defining the tuning curves as ***r*** ≈ ***φ***(***κ***) following **Theorem 4**. Then, for a generic nonlinear embedding function, we have (1 − ***P***_***N***_)***φ***(***κ***) ≠ 0, *i.e*., ***r***_⊥_ is in general a non-constant function of ***φ***(***κ***). As per **Definition 4**, these deviations (when they exist) increase the linear dimensionality beyond *K* and are associated with a non-zero manifold curvature. Finally, since **Lemma 5** ensures that ***κ***(*t*) does not depend on ***r***_⊥_(t), this representation is redundant as far as LPU operation is concerned.

**Lemma 6** suggests that even though the neural activity components orthogonal to the causally coding subspace do not affect the LPU operation, they can contain redundant information on the latent variables due to nonlinear embedding maps. On the other hand, the embedding has to be nonlinear to achieve universal approximation in the case of linear encoding (**Table S2**), thus this redundancy becomes practically unavoidable (See **Fig. 5**). **Theorem 5** combines **Lemmas 5** and **6** into a unified statement:

##### Theorem 5

**Proof:** As shown in **Lemma 6**, neural activities in directions perpendicular to the causally coding subspace can be tuned to latent variables. In the main text, we refer to such directions as acausal coding directions. These directions are tuned to the latents, so perturbing them might appear to affect the computation. However, **Lemma 5** shows that latent variables and their dynamics depend only on the components of neural activities residing on the causally coding subspace. Finally, the same lemma shows that deviations in the complementary subspace decay exponentially. Bringing these facts together concludes the proof.

Essentially, **Theorem 5** formalizes the notion that, for the general class of biologically plausible networks formalized in Eq. (2), neurons contribute to the latent code through their encoding weights, ***N***, whereas their tuning properties are determined jointly by the embedding weights, ***M***, and the element-wise nonlinearities. Latter, which is essential for universal computation under linear encoding maps, inevitably gives rise to redundancies in neural tuning properties. Therefore, designing perturbation experiments (using the embedding properties ***M***; as illustrated in **Fig. 5b-c**) based on neural tuning properties would be a suboptimal experimental strategy, as these properties may not reliably indicate the neuron’s true computational role in the LPU, for which the properties of the encoding, ***N***, need to be known.

#### Simulation experiments on effective and ineffective network perturbations

For the perturbation experiments in **Figure 5b-c**, we used the example RNN with 100 neurons from **Fig. 1c-e**, which was trained to perform a 2-bit flip-flop task. We generated 100 new neurons by sampling embedding weights from a discrete probability distribution defined on (−1, 0, 1), with probabilities (0. 1, 0. 8, 0. 1) such that roughly 20 *out* of the new 100 neurons would be tuning for the flip-flop states. The encoding weights for these neurons were zero. We refer to these as acausally coding neurons, whereas the original 100 neurons are the causally coding ones. Then, we generated the weight matrix subserving the LPU by first defining 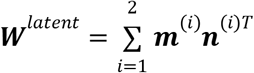. The total weight matrix was the summation of this structured component plus a randomly sampled component (***W***^*random*^) from a Gaussian distribution with zero mean and s.d. of 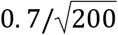, *i.e*., ***W***^*rec*^ = **W**^*latent*^ + ***W***^*random*^. The random connections allowed the newly generated neurons to influence the first group, thus preventing a trivial interpretation of the perturbation results due to lack of connectivity between causally and acausally coding neurons. We used the latent encoding weights to define the read*out* from the coding neurons. To obtain a read*out* from the acausally coding neurons, we trained a linear regression on a sample of 1,000 data points.

For the intervention experiments, we constructed two input vectors that let us perturb the causally and acausally coding neurons separately. In each vector, every entry was set to ± 1 according to the sign of that neuron’s embedding weight onto the first flip-flop (the first latent dimension). The first vector was non-zero only on the causally coding neurons and zero on the acausally coding ones; the second was non-zero only on the acausally coding neurons and zero on the causally coding ones. We appended these two vectors as additional input-weight columns to the RNN, giving independent interventional access to each group while keeping the perturbation aligned with the first flip-flop’s (causally or acausally) coding direction.

For the first experiment in **Fig. 5c**, we initialized the new network at the state (1,-1), and then applied a single pulse (pulse value = −1) at [500,505]ms interval to the causally coding neurons. For the second experiment in **Fig. 5c**, we initialized the new network at the state (1,−1), and then applied multiple pulses (pulse value = −1) at [500,600]ms interval to the acausally coding neurons. Both pulses were designed to switch the state to (−1,−1), though only the first intervention was successful.

### Continuous control of latent variables

This section describes the simulations we performed in **Figs. 5d-f** and **S6**. In these experiments, the encoding weights enable steering a continuous latent state with a linear gain. This linear control capability persists even when the connectome is dominated by random connectivity. We first describe the task and the trained networks, then the identification of the latent variable, the perturbation protocol, and finally the methodological details of the analyses performed.

To study control of a continuous variable, we trained rank-one basic RNNs (Eq. (15)) on a single-channel delayed addition task. In each trial, a scalar input channel was mostly zero through*out* except at two random time points *t*_1_, t_2_ with *t*_1_ < t_2_, during which two random amplitudes *a*_1_ and *a*_2_ were drawn independently from a uniform distribution on [0, 1]. The two pulses were placed in the first and second halves of the trial, respectively. Specifically, *t*_1_ was sampled uniformly from the first half and *t*_2_ from the second half, with a margin of three time steps from the trial edges and from the midpoint. The network was trained to *out*put the sum of these values, *a*_1_ + *a*_2_, at the end of the trial through a separately trained read*out*. Each network had N = 10,000 neurons, rank one, a hyperbolic-tangent nonlinearity, and followed the discretized dynamics

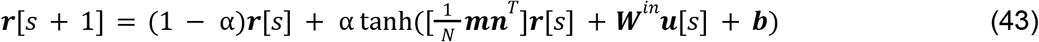

with discretization ratio α = 0. 25, Δ*t* = 2. 5ms, and τ = 10ms. Each trial consisted of *T* = 50 time steps, *i.e*., lasted 125-ms trial. The behavioral *out*put was read *out* linearly from the neural activities following 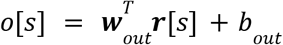, where ***w***_*out*_ ∈ ℝ ^*N*^ is the learned read*out* vector and *b*_*out*_ ∈ ℝ is the *out*put bias. The single causal coding direction of each network was stored in its encoding vector, ***W*** = ***mn***^T^ /N *i.e*., the non-trivial right eigenvector of the recurrent connectivity, which is proportional to ***n*** (cf. Eq. (4)). In the end, we successfully trained 51 rank-one RNNs whose held-*out* task accuracy reached at least 0.95.

For a rank-one network, the latent variable is linearly encoded as 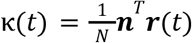. Because the latent basis is identifiable only up to a scalar (rescaling ***m*** → ***am*** and ***n*** → ***a***^−1^ ***n*** leaves ***W*** = ***mn***^*T*^ /*N*, and hence the neural dynamics, unchanged), we calibrated κ into the units of the target *out*put before comparing interventions. Concretely, on an unperturbed calibration batch (2,000 trials, discarding the initial transient by using time points *s* > 10), we performed a linear regression of the network read*out* onto the encoded latent variable. We then rescaled the weights as ***m*** → *a****m*** and ***n*** → *a*^−1^ ***n*** using the slope of the regression used this to define the latent variables. When we visualize the latent variable in **Fig. 5e**, we also use the offset for direct comparison with the *out*put read*out*. All latent deviations reported are in these calibrated units.

For our intervention experiments (**Fig. 5d-f**), we delivered a single additive pulse at the midpoint of the trial as an external input through (unit norm) input weights, which are sampled to act as perturbation vectors. The perturbation strength therefore quantifies the norm of the perturbation across all neurons. In our experiments, we performed perturbations through the following directions (all unit-norm unless noted): (i) the *encoding* direction, the unit vector along ***n***, which re-enters the LPU and therefore constitutes a causal intervention; (ii) the *readout* direction, the unit vector along ***w***_*out*_, which correlates with the latent variable by construction but does not necessarily re-enter the latent dynamics (often an acausal direction); (iii) the *embedding* direction, the unit vector along *m*; (iv) the *encoding-orthogonal* direction, a random unit vector with components parallel to ***n*** deleted and renormalized; (v) a *random* unit direction; and (vi) the trial-averaged locally optimal direction. The locally optimal direction for a given trial is the fixed-norm perturbation that maximizes the squared instantaneous change in the latent variable. This can be analytically computed from Eq. (43) with a constrained optimization setup:

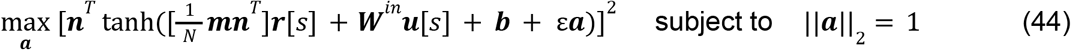

In the limit ε → 0, the solution is 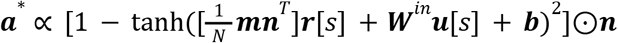, where ⊙ refers to element-wise multiplication. We computed this direction at the time of the intervention, *s* = *T*/2, for each trial. To obtain one direction that can be used for a generic application, we averaged these vectors across all trials.

For each network and each perturbation direction, we swept the perturbation strength over 48 values logarithmically spaced from 10^−3^ to 10^6^. For each ε value, we generated a batch of 500 delayed-addition trials, run the network dynamics forward once with*out* any perturbation to obtain the baseline latent variable κ_0_ at the final time step, and ran the same batch again with the perturbation injected at *s* = *T*/2 to obtain κ_*int*_. We defined the end-of-trial latent deviation per trial as Δκ = κ_int_ – κ_0_. We used both Δκ and |Δκ| and averaged them across trials to obtain a network-level summary statistics for a given experiment. We aggregated these across 51 well-trained networks. **Fig. 5f** plots the mean |Δκ| at the end of the trial against ε values on log-log axes, with shaded regions denoting ±1 s.d. across networks. To quantify the proportional control achieved through the encoding direction, we fitted a line to log_10_ (Δκ) versus log_10_ (ε) over the ε ∈ [10^−3^, 10^2^] moderate range (restricted to points with positive signed mean) and reported the slope and *R*^2^.

For the illustrative single trial in **Fig. 5d**, we applied the protocol above with ε = 1 to an example trial of a trained network. The perturbations were registered through the encoding and read*out* directions. For the phase portrait in **Fig. 5e**, we sampled latent variables and the *out*put variables from a grid, *i.e*., κ ∈ [0, 2] and *y* ∈ [0, 2]. The deviations in both variables were computed using Eq. (43) projected onto 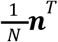 and 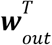, respectively. Specifically, for a given pair (κ, *y*), we computed the changes (Δκ, Δ*y*) predicted by a single step evolution of the network dynamics and plotted the phase portrait. To compute the log-speed values, we first computed the norm of the vector (Δκ, Δ*y*) and then took the natural logarithm of the resulting value. The trials from **Fig. 5d** were plotted on top of this phase portrait.

We next tested whether the capability to continuously control the latent computations survives the addition of dense random connectivity. Specifically, we augmented the full connectome as 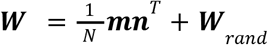 where the entries of ***W***_ran*d*_ were drawn i.i.d. from a zero-mean Gaussian with variance *g*^2^/N. For each network we kept the learned ***m*** and ***n*** and drew ***W***_*rand*_ afresh for four samples per network and a given *g* value. For each subsequent test of continuous control, we swept over *g* ∈ [0, 1. 5]. In a randomly drawn full-rank network, *g* = 1 denotes the edge of stability (see **Fig. S5**).

For the task-accuracy curve (**Fig. S6a**), we ran the dynamics of the augmented networks in new trials and measured the final read*out*s. Accuracy was assessed based on the original training criteria of the networks; namely, a trial was deemed accurate when (prediction - target) < 0.04. For each of the four samples of given network and *g* value, batches of 500 trials were assessed. For the perturbation-response curves (**Fig. S6b** and **S6c**), we repeated the sweep procedure of **Fig. 5f** in the augmented networks, but this time using only the encoding weights for control and using batches of 64 trials. Furthermore, we defined the read*out* error as “prediction − target values at the end of unperturbed trials” and reported the mean (the systematic bias) and standard deviation (the spread) as functions of *g* (mean ± s.e.m. across networks, **Fig. S6d**). Finally, to quantify how much of the recurrent connectivity remained attributable to the low-rank (LPU) component (**Fig. S6e**), we computed the percentage power 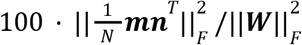.

### Robustness of LPU dynamics to representational drift

In this section, we detail our studies regarding the robustness of LPU dynamics to representational drift. First, starting from a formal definition of tuning curves in our framework, we develop a theoretical model of representational drift in RNNs. Then, we discuss some statistical assumptions on the coding properties of individual neurons and three types of qualitatively different drifts. With these assumptions, we then state and prove our main theoretical result showing LPU dynamics can remain robust, but not invariant to the representational drift (**Theorem 6**). Finally, we conclude with the experimental details of our RNN simulations in **Figure 6**.

#### A theoretical model of representational drift in RNNs

For any artificial and biological neural network that satisfies **Definition 1, Theorem 4** readily allows defining tuning curves through the relationship ***r***(***κ***) ≈ ***φ***(***κ***). Yet, since representational drift refers to the changes of neural tuning properties in biological neural networks, we now return to the particular class of biologically plausible networks described by Eqs. (13-14) and **Theorem 1** and define the tuning curves of individual neurons.

Recall that this class of networks makes use of *L* synaptic compartments, each of which contributes a set of synaptic parameters ***W***^(*j*)^ to implement latent variables (cf. discussion right after Eq. (14)). Here, we first reorganize the inputs of the nonlinearity in Eq. (14) to explicitly emphasize these resulting latent variables, omitting the explicit synaptic compartment they originate from. Specifically, we define the *p*th latent variable as 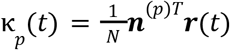, where 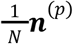 the encoding weight that corresponds to one of the row vectors in the collection (***N***^(1)^, …, ***N***^(*L*)^) In total, there are *K* many such variables described by the collection (***n***^(1)^, …, ***n***^(K)^). The latent variables embed the neural dynamics via the vectors (m^(1)^, …, m^(K)^) with corresponding to one of the column vectors in the collection (***M***^(1)^, …, ***M***^(L)^). Then, Eq. (14) can be rewritten as:

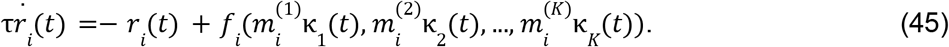

Representational drift refers to constant changes in the tuning curves of neural responses to specific stimuli or behaviorally relevant variables^7^. In our framework, these experimental and behavioral constructs are presumably described by latent variables that evolve slower compared to the neural time constants (**Theorem 5**), which allows defining the tuning curves (**Theorem 4**):

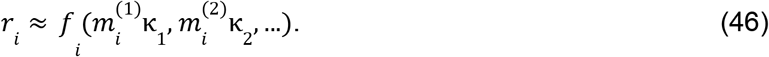

This definition suggests a straightforward method for studying these changes in RNNs. Once the latent variables are aligned to the behaviorally relevant quantities (*e.g*., each κ_i_(t) corresponds to *o*_*i*_ (*t*), the *i*th *out*put variable of the task), we can vary the tuning curves (simulating representational drift) by introducing changes to a particular embedding weight, ***m***^(*d*)^:

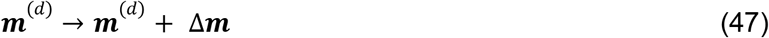

where the superscript *d* refers to the specific embedding weight receiving the update. Notably, a generic drift of the form in Eq. (47) will induce a change in the tuning curves in Eq. (46). For a network that is robust to the drift, the following is expected: If the latent variables, ***κ***(*t*), follow a dynamical system that is invariant to these updates, Δ***m***, then even though the coding properties of the individual neurons undergo changes, the LPUs and the behavioral read*out*s from these LPUs remain robust to these changes.

#### Robustness of LPUs to the representational drift

Now, we study the inherent robustness of LPUs to the representational drift under certain theoretical conditions. To be able to state these conditions, we start by defining the Jacobian of the function in Eq. (46) as:

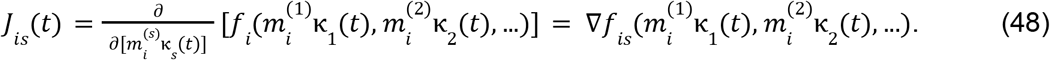

This Jacobian term can intuitively be thought as quantifying the saturation level of individual neurons, *i.e*., quantifying the changes in neural activities when κ_s_ (*t*) are briefly perturbed. As an intuitive illustration, consider a neuron with the tuning relationship *r*(κ) = taNh(κ). Here, the neuron can be considered inactive when κ = 0 and active when |κ| is large. Then, for an inactive neuron, we have *J*(κ) = 1 – ta*N*h+^2^ (κ) ≈ 1. In words, this means that small changes in κ linearly effect neural activities. On the other hand, active neurons are almost insensitive to changes in the latent variable since *J*(κ) ≈ 0 for |κ| → ∞. Hence, *J*(κ) intuitively provides a quantification for the saturation level of the neuron’s responses, *i.e*., whether further increases in the latent variable values would lead to pronounced changes in the neural activities. With this definition, we now make two key assumptions:

1. We assume that the drift in the embedding vector ***m***^(*d*)^ is either purely random, or for non-random drift, we require a conditional orthogonality between drift and encoding dimensions, *i.e*., *E*[*n*^(p)^ Δ*m*|*J*] = 0 for *p* = 1, …, K. Here, *J*_*d*_ refers to the random variable defined over neurons describing the Jacobian in Eq. (48).
2. Each entry of the encoding and embedding weights are sampled following some unknown probability distribution and have self-averaging expectations *E*[*n*^(p)^ *m*^(*p*’)^] for any (*p, p*’) combination such that the law of large numbers ensures convergence to mean. Similarly, combinations with *J*_*d*_ are also self-averaging.

The second assumption is common in the field as discussed earlier. It suggests that the distributional characteristics of synaptic connections, rather than the precise values of individual synapses, are the primary factors driving computation. The first assumption is more restrictive, suggesting that either the system is subject to random drift, or, for systems with structured drift, changes in the synaptic weights are assumed to be orthogonal to the causally coding subspace and by factors independent of which neurons are active (and thus saturated), *i.e*., the conditional orthogonality with respect to *J*_*id*_. With these assumptions, we are now ready to state the following theorem:

##### Theorem 6

*(Computational robustness to representational drift). Let* ***r***(*t*) *denote the neural activity encoding an LPU with latent variables* ***κ***(*t*). *Let the embedding weights change via* ***m***^(*d*)^ → ***m***^(*d*)^ + Δ***m***. U*nder the statistical assumptions defined above, the latent dynamics remain preserved up to second-order corrections* 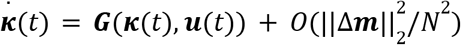 *long as the drift is purely random or* Δ***m*** is *orthogonal to the causal coding dimensions*.

**Proof:** Consider the embedding relationship 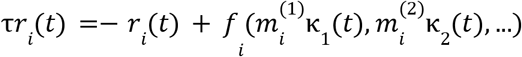. For a small change in a particular embedding weight, ***m***^(*d*)^ → ***m***^(*d*)^ + Δ***m***, the neural activities change following: 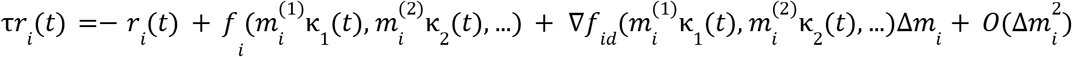

As long as 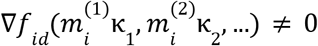, a generic update changes the time dynamics of the neuron *i* proportional to *O*(Δ*m*_i_) for each time point. Hence, as a first step, this shows that neural activities will generically change as a result of the drift.

Next, we consider the latent dynamics that follow the equation:

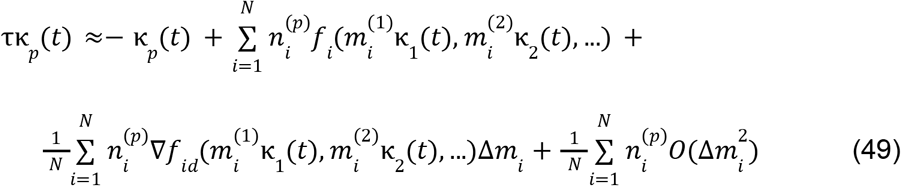

where we performed the Taylor expansion of the nonlinearity up to a second order 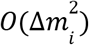. The final term is 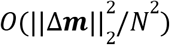, whereas in the limit *N* → ∞, we can rewrite the first-order correction as:

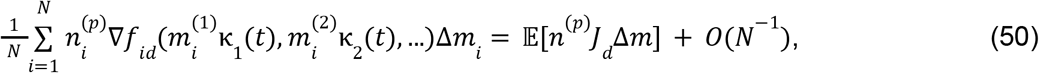

where the sample mean (over the neurons) is replaced by the expectation value over the probability distributions governing these quantities. This is possible by the second regularity assumption. Then, for a purely random and uncorrelated Δ*m*, Eq. (50) vanishes. Similarly, the second assumption covers the second case via the tower property of the expectations *E*[[*n*^(N)^ *J*_d_ Δ*m*]] = *E*[*J*_*d*_ *E*[*n*^(p)^ Δ*m*|*J*_*d*_]] = 0. This concludes the proof.

**Theorem 6** suggests that as long as the drift is in an orthogonal direction to the encoding weights, conditioned on the saturation levels of the neurons (*J*_*d*_) which is assumed to be sampled from an unknown distribution randomly for each neuron, the latent dynamics will remain invariant to the first-order changes in *O*(Δ***m***). Yet, how do we ensure this conditional orthogonality? For this, we can consider two distinct cases:

1. For a fully random drift, the condition is trivially satisfied if the mean of the drift is zero.
2. For a structured drift scenario, however, this implies that changes in the synaptic weights are driven by factors independent of the specific saturation properties of individual neurons.

To understand the second condition more intuitively, consider the Jacobian *J*_*id*_’s relation to the operation regime of the neuron dynamics (*e.g*., saturation or linear for taNh(·) nonlinearity in our example above). Then, assuming that the LPU dynamics remains invariant, *J*_*id*_ depends primarily on the embedding, not encoding, weights, *i.e*., the tuning (not coding) properties of the individual neurons. Hence, it is not implausible, though open to experimental confirmation, that neuronal tuning properties that are known to be redundant can be chosen as such to approximately satisfy constraints similar to one presented here. Finally, it is worth noting that **Theorem 6** states robustness (up to first-order changes), not invariance. In the event of substantial changes in Δ***m***, the element-wise nonlinearities mix and match the linear subspaces (mathematically arising as increasing numbers of higher order terms in the Taylor approximation above), preventing the existence of a perfectly drift-invariant LPU in general, as demonstrated in **Fig. 6**. Having stated **Theorem 6**, we can combine it with **Lemma 2** to define a class of linear decoders that can remain robust to representational drift:

##### Corollary 2

*(Class of linear decoders robust to representational drift). Let* ***r***(*t*) ∈ ℝ ^*N*^ *represent the neural activities and* ***κ***(*t*)∈ ℝ^*K*^ *be the corresponding latent variables obtained via a linear encoding. Let* ***P***_***N***_ *be the projection operator onto the causally coding subspace and ô* ∈ ℝ^*B*^ de*note the behavior readouts with N* ≫ *K* ≥ *B. Define a linear decoder* ***ψ***(r(*t*)) = ***W***_*out*_ ***r***(*t*) ***+ b***_*out*_, *where* ***W***_*out*_ ∈ ℝ^B×N^ *and* ***b***_*out*_ ∈ ℝ^B^. *Assume that the network undergoes representational drift with the conditions specified in* ***Theorem 6***. *As long as* ***W***_*out*_ (1 − ***P***) = 0, ***ψ***(r(*t*)) *remains robust to the first-order changes in* Δ***m***.

P**roof: Lemma 2** allows defining an equivalent latent decoder from the latent variables and **Theorem 6** ensures that these latent variables follow an invariant set of dynamical systems. Thus, the *out*put of the linear read*out* is also robust to the representational drift, *i.e*., remains invariant to the first-order changes in Δ***m***, concluding the proof.

It is worth recalling that the dimensional bottleneck (*i.e*., ***W***_*out*_ (1 – ***P***_*N*_) = 0) enforced on this class of linear decoders may prevent them from using redundant information in the full neural activity, leading to suboptimal decoding performances for a given time point. Similar trade-offs between robustness and absolute optimality are common and well known in the statistics literature^109^.

#### Evaluations of representational drift in task-trained RNNs

For the drift experiments in the main text, we used several networks trained to perform 3-bit flip-flop tasks. Specifically, we used the 50 networks with 100 neurons using the two-step training paradigm described above. Since training RNNs with many neurons is computationally expensive, we used an alternative resampling approach to simulate RNNs with more than 100 neurons. Specifically, each neuron in the RNN is unique defined by 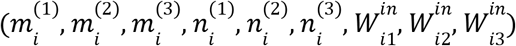. We artificially created new neurons by resampling from this set and later normalizing the encoding weight values 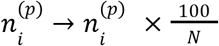. Since adding new neurons and then normalizing their contribution to the latent variables lead to the same dynamical system equations in expectation, this approach for increasing the neuron numbers exactly maintains the original LPUs.

To simulate the drift in RNNs, we evolved each network for 2000 data points, after which we performed an instantaneous update in embedding weights, ***m***^(*p*)^: = ***m***^(*p*)^ ***+*** Δ***m***^(*p*)^ for *p* = 1, 2, 3, and allowed the network to run for another 3000 time points after the drift. To test the robustness of the networks to different types of representational drift, we picked the changes, Δ***m***^(*p*)^, in three distinct manners. First, we sampled Δ***m***^(*p*)^ from a random Gaussian with zero mean and *g*_*drift*_ sd. This constituted the case of the fully random drift. For the targeted drifts, we projected the random weights to the directions parallel or orthogonal to the encoding weight space, spanned by ***P***_N_. To account for changes in power due to projecting, we re-scaled Δ***m***^(*p*)^ to have the same standard deviation as the one before the projection.

To decode the internal states of the networks, we first estimated the latent variables by either using the full set of neurons, or by using only the first 100 neurons. Then, we computed the internal states of the RNN by thresholding the estimated latent variables wrt zero (*e.g*., κ_i_ = 0. 1 corresponds to a state of 1, whereas κ_i_ =− 0. 8 to a state of −1). These states were compared with the *out*puts using a 1-0 loss function. For each network, we performed three distinct runs using randomly chosen initial conditions and calculated mean accuracies across the three runs. When calculating these mean accuracies, we used only the last 2000 time points, allowing the transient responses right after the drifts to settle.

For the illustrative tuning curve example in **Fig. 6b**, we reused the example RNN from **Fig. 1c-e**. The applied drift had a strength of *g*_dri*ft*_ = 0. 2 and was orthogonal to the causally coding subspace. We used the same network for the illustrations in **Fig. 6e-h**, where we evolved the RNN with an input noise drawn from a Gaussian distribution of mean zero and s.d. 10^−2^. The noise was added independently at each time point before the evaluation of the nonlinearity. For the RNNs solving the 3-bit flip-flop task, which we reported in **Fig. 6d, i-j**, we set the standard deviation of the noise to 10^−1^. To obtain the fitted curves in these panels, we fit a sigmoid function between 1 and 1/8 (chance level) for the accuracy against the logarithm of the drift strength values. *g*_*half*_ values correspond to the transition points of the fitted sigmoid curves, in which the sigmoid terms halve.

## Supplementary Note 1: Additional theoretical considerations on the latent processing units

### 1. Overview

In this supplementary note, we first discuss the properties of three distinct encoding and embedding maps, excluding the linear encoding and nonlinear embedding combination that has been the focus of the main text. Then, we demonstrate three distinct theoretical advantages of linear encoding and nonlinear embedding maps: guaranteed existence of LPUs (both maps), complex tuning curves (nonlinear embedding), and linear identifiability of latent dynamics (linear encoding). Finally, we conclude our theoretical discussion in this note with how our encoding-embedding framework can describe the existing dynamical system models regularly studied in the existing literature.

### 2. Types of encoding and embedding maps

Here, we show that the dynamical nature of the embedding function leads seamlessly to dynamical system definitions for both ***κ***(*t*) and ***r***(*t*). This represents a significant departure from the static reconstruction of observed neural activities, *e.g*., performed by traditional encoder-decoder models^1–3^. The type of encoding and embedding maps, linear or nonlinear, divides the networks into four groups with distinct geometrical properties. The geometry then significantly impacts the statistical properties and complexity of the neural computation that can be performed by the dynamical systems (discussed below and summarized in **Table S2**). For simplicity and since we are interested in computations that can be intrinsically achieved by the dynamical systems, we assume that there are no inputs (and biases) unless otherwise specified. We now study some (non-exhaustive) properties of dynamical systems belonging to each of the four groups mentioned above.

#### 2.1. Linear encoding and linear embedding

Assuming that both the encoding and the embedding are linear leads to the following relationship:

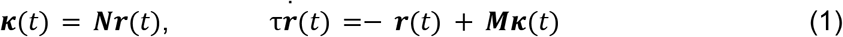

where ***N*** ∈ ℝ ^*K*×*N*^, and ***M*** ∈ ℝ ^*N*×*K*^ are encoding and embedding matrices, respectively. Unless otherwise stated, we always assume that these matrices are maximally ranked, *i.e*., both have rank *K*, and ***r***(*t*) ∈ ℝ^*N*^, ***κ***(*t*) ∈ ℝ^*K*^, and τ > 0 are defined as in the main text. Combining both equations, we obtain the following relationship for the neural dynamics:

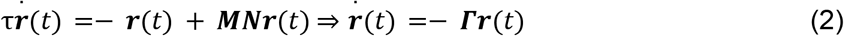

where we define an effective decay matrix as ***Γ*** = τ^−1^ (*I* − ***MN***). In the case, where both encoding and embedding maps are linear, neural dynamics are driven by a linear dynamical system. Then, the neural dynamics can be written analytically as 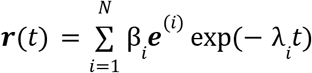 for some β_*i*_ ∈ ℂ, and the eigenvalues (λ_*i*_ ∈ ℂ) and eigenvectors (***e***^(*i*)^ ∈ ℂ^*N*^) of ***Γ***. Consequently, the latent dynamics also follow a linear equation, 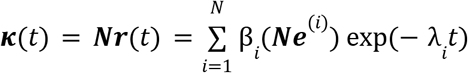. In turn, this implies that ***κ***(*t*) can only decay, oscillate, or blow up. Hence, for ***κ***(*t*) to support a diverse set of computations, both encoding and embedding cannot be linear.

#### 2.2. Nonlinear encoding and linear embedding

Next, we consider the case where encoding is nonlinear and embedding is linear:

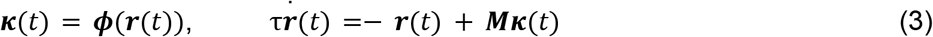

for some nonlinear encoding function, ***ϕ***(·): ℝ^*N*^ → ℝ^*K*^, and rest are defined as before. This type of encoding requires additional conditions on the nonlinearity following:

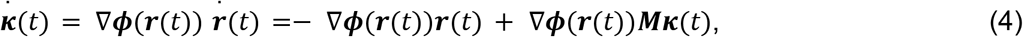

in which the RHS should not explicitly depend on ***r***(*t*) for ***κ***(*t*) to constitute a closed dynamical system. Currently, the breadth of nonlinear encoding functions that can produce self-sustained dynamics for ***κ***(*t*) are unknown, though there should likely be some constraints on ∇***ϕ***(***r***(*t*)).

For the sake of studying the properties of such a system, we next consider the embedding equation closely. Specifically, consider the formal solution to the embedding equation, 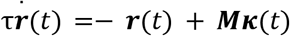, which takes the following format:

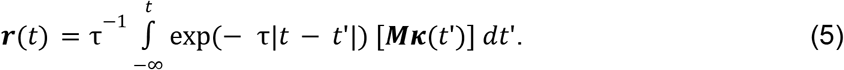

Here, ***r***(*t*) becomes a weighted sum of latent variables, with some memory kernel τ^−1^ exp(− τ(*t* − *t*′)). When |*t* − *t*′| ≫ τ^−1^, the exponential term is nearly zero, and thus the non-negligible contribution to the integral comes from a region around |*t* − *t*′| ~ *O*(τ)^−1^.

An important insight, one we have formalized in the main text for **Theorem 5**, is that there is often several orders of magnitude separation between the synaptic and behavioral timescales. Specifically, in biological neural networks, the synaptic timescales are τ ~ *O*(*ms*), whereas the behavioral timescales, which presumably correspond to the timescales with significant variations in ***κ***(*t*′), are *O*(*s*). In mathematical terms, this separation of scales can be stated by assuming that ***κ***(*t*′) would be approximately constant for |*t* − *t*′| ~ *O*(τ^−1^) such that we can approximately fix ***κ***(*t*′) ≈ ***κ***. Then, the embedding state variables, ***r***(*t*), becomes a linear transformation of the latent variables following 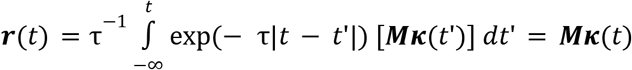. Thus, the dimension of the linear subspace spanned by the state variables ***r***(*t*) is at most equal to the latent dimensionality, *i.e*., dimension of the linear subspace spanned by the latent variables ***κ***(*t*). Yet, since nonlinearity is provided by the (nonlinear) encoding by definition, this class of networks could support complex computation other than linear dynamics. How ***κ***(*t*) can both be a nonlinear function of ***r***(*t*) but also have the same linear dimensionality as ***r***(*t*) remains to be explored, though authors are currently unaware of a construction in the literature fitting this combination of encoding-embedding maps.

#### 2.3. Nonlinear encoding and nonlinear embedding

A typical neural network model involves reconstructing neural activities using low-dimensional bottlenecks. Defining the output of the decoder to be the time derivative of neural activities would be consistent with the definition of a dynamical embedding. Then, the resulting model would have nonlinear encoding and nonlinear embedding maps. Since this class of relationships subsumes the linear encoding and nonlinear embedding case, which we show can lead to universal approximators in the main text (**Theorem 1**), the resulting RNN architectures can allow complex computations. However, it is not exactly clear under which conditions the latent variables follow closed-form dynamical system equations, *i.e*., constitute LPUs. Moreover, while previous work has proven that latent variables can be identified up to linear transformations in this scenario^4^, it is not a priori clear whether these guarantees generalize to latent dynamics governing these variables. We leave it to future work to study these models, though note that several biologically desirable properties we proved in this work require assumption of linear encoding (see, *e.g*., **§3**).

### 3. Theoretical benefits of linear encoding and nonlinear embedding

The case of linear encoding and nonlinear embedding follows the set of equations:

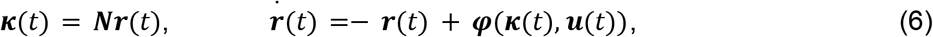

where ***N*** refers to the encoding matrix and ***φ***(·) nonlinear embedding function. This linear encoding and nonlinear embedding relationship leads to well-known recurrent neural network architectures, and reproduces two regularly used architectures, which we will discuss in the next section. Here, we discuss mathematical details behind the advantages of assuming linear encoding and nonlinear embedding maps.

#### 3.1. Self-sustained latent dynamical systems

The linear encoding map in Eq. (6) provides the latent dynamics with the property of self-sufficiency, *i.e*., the first condition for a latent system to be considered as LPU, following:

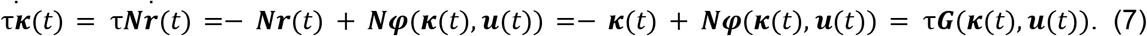

On the other hand, the nonlinearity of the embedding can endow the network with universal approximation property. Specifically, consider the general class of neural networks we considered in the main text, which led to the following latent dynamics:

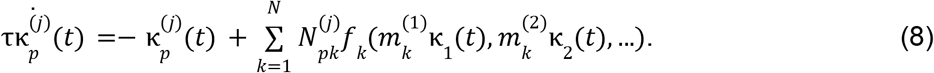

This equation has the form of a multilayer neural network with input ***κ*** and output τ***G***(***κ***(*t*)) + ***κ***(*t*). Assuming a nonpolynomial nonlinearity, depending on the form of the network (for example, when the inputs of the nonlinearity are additive, *i.e*., *f*_*i*_ (*x*_1_, *x*_2_, …) = tanh(*x*_1_ + *x*_2_ +…) holds), this multilayer neural network becomes a universal approximator for the latent flow maps (**Theorem 1**). The study of more general forms for *f*_*i*_(·), especially inspired by the biological models of dendritic computation, is an interesting direction left for future work.

#### 3.2. Nonlinear tuning curves

In the main text, we argued that the separation of timescales between latent variables and neural activities may allow the LPU dynamics to remain approximately invariant for short time intervals. In this section, we first formalize this statement such that *κ*(*t*′) ≈ *κ*(*t*) for *t*′ ∈ [*t, t* + *O*(τ)], but may vary for some desired latent timescales τ ≫ τ. Realistically, one can imagine τ ~ 1 − 10*ms* for the processes occurring within individual neurons and τ_*eff*_ ~ 100 − 1, 000*ms* being the timescale of the latent variables that putatively subserve the behavior. In this case, despite the dynamical nature of the embedding, it is possible to define approximate tuning curves as if the embedding function was static. To do so, we first define ϵ-stability:

##### Definition A1

(ϵ-stability). We refer to a set of variables ***κ***(*t*) as ϵ-stable for the time window [*t, t* + *T*] if for ϵ > 0, following holds: ∀*i* = 1, …, *K*, ∀*t*′, *t*′′ ∈ [*t, t* + *T*], |*κ*_*i*_ (*t*′) − *κ*_*i*_ (*t*′′)| ≤ ϵ.

Here, ϵ is a user-defined (small) parameter which can, for example, be set to the noise level. Then, if we assume that ***κ***(*t*) are ϵ-stable for the time interval *t* ∈ [*t*_1_, *t*_1_ + *T*] with *T* ≫ τ, in the absence of any input and setting 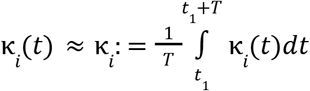 as constant within this regime, the embedding map becomes 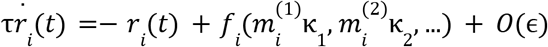. Ignoring the terms in *O*(ϵ), the above equation leads to the steady-state solution:

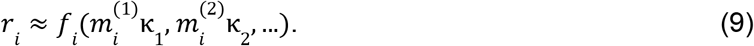

Notably, this relationship resembles what is traditionally referred to as a tuning curve. Unlike traditional research, however, latent variables here are defined with respect to the internal latent variables, not experimentally relevant stimuli or behavioral variables. Moreover, it should be noted that the noisy nature of neural activities, *e.g*., addition of random noise to the embedding, would mean that this relationship likely holds on average over, *e.g*., trials. This is in line with how tuning curves are often computed in practice^5^. However, since internal variables have an inherent *GL*_*K*_(ℝ) symmetry, *i.e*., linear transformations of ***κ***(*t*) are also valid as latent variables (see below), we refer to Eq. (9) as a tuning curve with the understanding that latent variables are aligned with the network outputs (see **Fig. 6**). Notably, Eq. (9) represents a nonlinear function that can potentially model complex tuning properties, *e.g*., by designing *f*_*i*_ as multilayer neural networks, as opposed to assuming a fixed form, *e.g*., *f*_*i*_ = tanh(·).

#### 3.3. Linear identifiability of latent dynamical systems

As the final note, we discuss the identifiability concerns of LPUs. Recall that for ***κ***(*t*) to be considered an LPU, its dynamics should not depend explicitly on ***r***(*t*). Although latent dynamical systems with linear encoding automatically satisfy this criterion (see above), there are infinitely many transformations that can lead to equivalent LPUs with different latent variables. Specifically, consider the class of transformations, 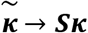, where ***S*** ∈ ℝ^*K*×*K*^ is an invertible matrix. After such transformations, 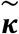 follows a dynamical system equation 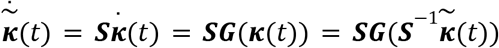. For any function of the latent variables, ***Ψ***(***κ***(*t*)), we can define a new (equivalent) decoder such that 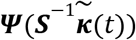 provides the same outputs. Finally, the embedding equation can be modified to satisfy 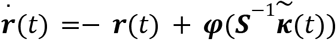. Combining all these facts together, after the transformation 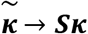, the resulting latent dynamical system leads to the same set of neural activities and can provide the same set of outputs. However, the new variables follow a different latent dynamical system, whose flow map is transformed following 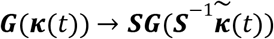.

This type of mathematical equivalence between latent dynamical systems, resulting from so called *GL*_*K*_ (ℝ) symmetry, is referred to as “linear identifiability” and is often the goal to achieve in the broader literature^4^. Broadly, the latent non-identifiability problem has been well known in independent component analysis (ICA), with established conditions for linear identifiability^6^. In the case of a nonlinear mixing function (called a decoder traditionally^7^, or a static embedding in our terminology), recent work has provided nonlinear identifiability conditions (See Ref. 6 for a review). Some nonlinear ICA methods have also been proposed for neural data, *e.g*., based on VAEs^8^ or contrastive learning^4^. Interestingly, unlike the prior work focusing on identifying latent variables without constraints on their dynamics^4^, enforcing linearity on the encoding map uniquely constrains the transformation that the flow map of the latent dynamical systems transforms in our framework.

Finally, it is worth noting that if a neuron is not tuned to any of the latent variables, then *GL*_*K*_(ℝ) symmetry cannot be used to turn the neuron into a tuned one. Specifically, if 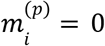 for all *p* = 1, …, *K*, such that the neuron *i* is not tuned to any of the latent variables, then the neuron 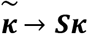 since 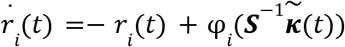 but φ (·) did remains untuned under the transformation not depend on ***κ*** to begin with.

### 4. Existing models in encoding-embedding framework

As an illustration of our framework, we first study the static nonlinear ICA models, which are defined with a direct map from latent variables to neural activities, in contrast with our dynamical embedding mapping latent variables to the time derivative of the neural activities. Then, we reproduce the prior results on low-rank RNNs^9–12^, and later a deterministic limit of recurrently switching linear dynamical systems^13–18^.

#### 4.1. Static nonlinear ICA models

A reasonable question one might ask is why define the embedding in a dynamical manner as we have done in **Definition 1** of the main text. To answer this question, we now consider a more traditional deep learning model, assuming no input for simplicity. In this model, the embedding map is static and therefore becomes a “decoder” of the neural activities (not of their time dynamics) from the latent variables:

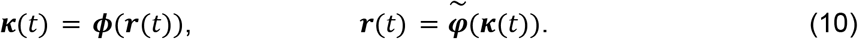

In this formulation, the time derivative of the latent variable depends on 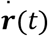 following 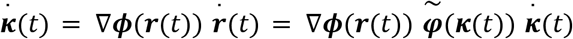. This is a tautology, following from the implicit assumption that the embedding reverts the encoding map: 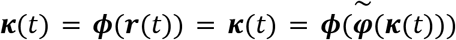. In other words, such a static model construction provides no information about latent or neural dynamics. Instead, as long as 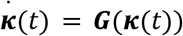 for some ***G***(·), it is possible to rearrange the terms to define a dynamical embedding:

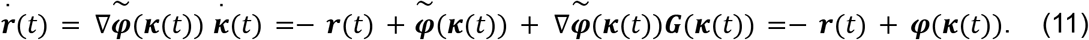

Thus, though it is possible to fit a post hoc model on a statically learned latent variable system such that 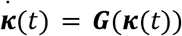, the resulting fit would be subsumed by the family of the dynamical embedding maps we introduced in **Definition 1** of the main text. Specifically, to define 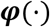, one needs to learn both 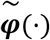 (from deep learning training) and ***G***(·) (from post-hoc fitting). In contrast, our formalism in **Definition 1** defines ***φ***(·) through the network architecture, *i.e*., without intermediate steps. An interesting recent direction enforces linear evolution in the latent variables and trains a neural network model for the encoding and (static) embedding maps^19^, which is likely limited in its modeling capabilities due to the linearity assumption but further research is needed to fully understand the capabilities of these models.

#### 4.2. A secondary form of basic RNNs

Next, we reproduce the previous work on low-rank RNNs within the encoding-embedding framework:

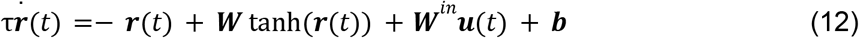

where the parameters are defined similar to before. Previous work subjected the network to a low-rank constraint such that ***W*** = ***MN***, where ***M*** and ***N*** are defined as encoding and embedding matrices. Then, this network can support a self-sustained latent dynamical system if neural activities are constrained in low-dimensional subspaces (reasons for which will be clear below)^10,12^. Here, for completeness, we briefly summarize the mathematical steps^10,12^. We start by defining the latent variables via the relationship:

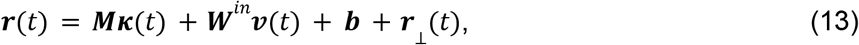

where 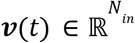 refers to the transformed inputs 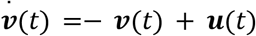, and ***r***_⊥_(*t*) is the neural activity orthogonal to the coding subspace spanned jointly by the column vectors of ***M*** and ***W***^*in*^. Inputting this into Eq. (12), Refs.^10,12^ arrive at the set of equations:

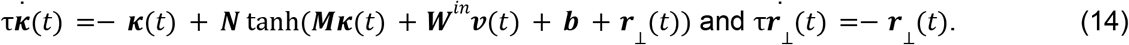

The second equation implies that any perturbation in the orthogonal direction would die out exponentially. Yet, any finite deviation, ***r***_⊥_ (*t*), leads to the first equation being not self-sufficient.

If we enforce the condition that ***r***_⊥_ (*t*) = 0 and assuming that ***M*** and ***W***^*in*^ are full-rank, Eq. (13) implies that the linear dimensionality of the neural activities is equal to the dimensionality of the latent dynamical system plus the number of inputs. Under these assumptions, the latent dynamical system equation can be rewritten to be self-contained as:

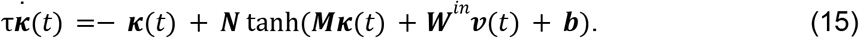

It should be noted that previous work has established this latent dynamical system, defined in Eq. (15), as universal approximators of *K*-dimensional dynamical systems with continuous flow maps^12^.

Notably, however, none of the equations above are in the form of the encoding-embedding relationship we defined in **Definition 1**. To cast them in this form, we first assume (for simplicity) that the column vectors of ***M*** and ***W***^*in*^ are all orthogonal to each other. Then, Eq. (13) can be inverted to define the encoding map as^10^:

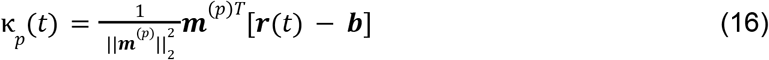

where ***m***^(*p*)^ is the *p*th column vector of ***M***. Notably, this is a linear function of ***r***(*t*). Similarly, using Eq. (12), we can define the embedding map as:

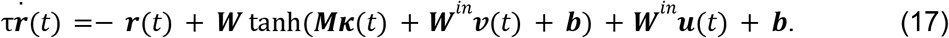

This is a nonlinear function of ***κ***(*t*), *i.e*., a nonlinear embedding. Thus, this latent dynamical system can be cast within our **Definition 1** and ***κ***(*t*) constitute an LPU as long as ***r***_⊥_(*t*) = 0. However, this also assumes that neural activities are restricted to lie in low-dimensional flat manifolds, in disagreement with the experimental work suggesting high linear dimensionality^20,21^.

#### 4.3. Recurrently switching linear dynamical systems

Now, we discuss a variant of the latent variable models known as recurrently switching linear dynamical systems or rSLDS^13–17^. We start by defining *L* (hyperparameter) latent states (*Z*_1_, *Z*_2_, …, *Z*_*L*_) such that for a given time, the network is in one of the given states *z*(*t*) = *Z*_*i*_ for some *i* = 1, …, *L*. Then, depending on the state, the latent variables follow a linear dynamical system:

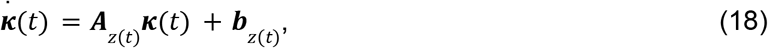

where ***A***_*z*(*t*)_ ∈ ℝ ^*K*×*K*^ and ***b***_*z*(*t*)_∈ ℝ are state dependent parameters, *i.e*., selected from a group of parameters, 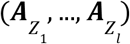 and 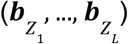, respectively. Similar to the RNNs described above, in this model, the neural activities are assumed to be defined by linear maps from the latent variables:

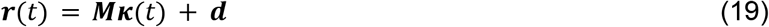

for some ***M*** ∈ ℝ^*N*×*K*^ and ***d*** ∈ ℝ^*N*^.

The traditional rSLDS model is not deterministic, rather considers a set of transition probabilities between latent states, which is often defined as a function of ***κ***(*t*) values. Therefore, in the most general case, it is not possible to assign individual states to particular ***κ***(*t*) values, since these internal states follow a stochastic process. Our encoding-embedding framework above, however, is designed for deterministic networks. Fortunately, in practice, it is often possible to approximately assign states to distinct latent variable combinations^14,16^. Inspired by this observation, we rewrite an equivalent form for Eq. (18) as:

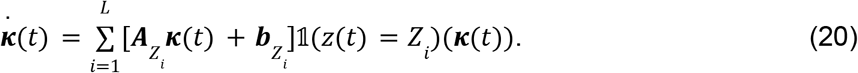

Here, 1(*z*(*t*) = *Z*_*i*_)(***κ***(*t*)), a function of ***κ***(*t*), is one for the state *Z*_*i*_ that corresponds to the current ***κ***(*t*) variables, zero otherwise. Notably, Eq. (20) constitutes a latent dynamical system, which is linearly encoded in neural activities (to see why, perform the inverse of Eq. (19) similar to before). Then, for the embedding relationship, we obtain:

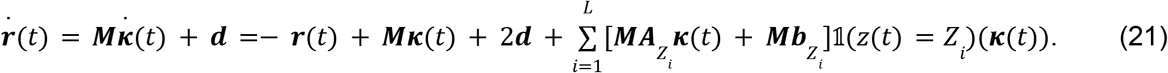

This embedding function is piecewise linear (hence nonlinear) in ***κ***(*t*), where the regions of linearity are marked by the internal states (*Z*_1_, …, *Z*_*L*_). Consequently, we can reformulate the deterministic limit of rSLDS as an instance of our encoding-embedding framework with linear encoding and nonlinear embedding maps.

## Supplementary Note 2: Constructing latent processing units in finite networks

### 1. Overview

In this supplementary note, we illustrate some common problems associated with network reconstruction, especially in the case of partial observation of the neural dynamics^1–3^. First, we discuss some regularity conditions on the distributions governing rank-one RNN parameters and introduce a sampling method to design one-dimensional latent processing units (LPUs) that have bistable dynamics. Then, we derive constraints on the synaptic connections between neurons such that latent dynamics can take place in desired timescales. Afterwards, inspired by previous work that aims to model functional connections between neurons using data-constrained RNN models^4,5^, we design a simple, yet insightful, procedure for reconstructing LPUs under partial observation of neurons. Our analyses reveal an analytical scaling relationship for the errors resulting from LPU (re-)construction, with two distinct empirically relevant predictions: i) a quantitative relationship for how many neurons are needed to accurately design the bistable LPU with desired timescales, and how well existing LPUs in networks with finite number of neurons can be reconstructed, with particular focus on the contrast in the sparse (*N*_*obs*_ ≪ *N*) and rich (*N*_*obs*_ ~ *N*) observation limits. Here, *N* refers to the number of recorded neurons, which can come close to *N* with the recent large-scale recording technologies^6^.

### 2. Theoretical analysis of timescales in a bistable latent dynamical system

Our goal in this section is to constrain the distribution of the embedding and encoding vectors, described by the random variable pair (*m, n*), such that the latent dynamical system approximates a bistable toy model in the mean-field limit, *i.e*., has a repelling fixed-point at the origin and two attractive fixed-points at *κ*^*^ =± 1. Then, we will study the quantitative behavior and stability of this system as a function of finite *N* neurons in the network.

#### 2.1. Regularity of the latent flow maps in the infinite networks

Assuming zero inputs and no biases, we note a generic flow-map for a one-dimensional LPU:

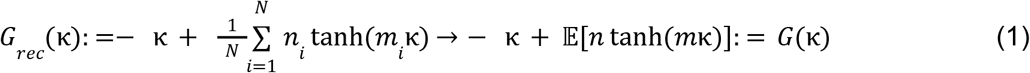

where → in Eq. (1) refers to the mean-field limit, *i.e*., *N* → ∞. Here, for a given *κ* ∈ ℝ, the central limit theorem (under the conditions that the expectation exists and the variance is finite, see **Proposition A1** below) ensures that the flow map, *G*_*rec*_ (*κ*), approximates a random variable with mean *G*(*κ*) and some vanishing variance as *N* → ∞.

##### Proposition A1 (Continuous differentiability of the flow map in 1D)

Let *κ* constitute a one-dimensional latent dynamical system governed by the flow map, *G*(*κ*), defined in Eq. (1) in the limit *N* → ∞. Let the pair (*m, n*) be sampled from a distribution with finite fourth moments, *i.e*., E[*m*^4^] < ∞ and E[*n*^4^] < ∞. Then, the following statements are true:

1. the flow map, *G*(*κ*), is continuously differentiable on ℝ, *i.e*., *G* ∈ *C*^1^ (ℝ),
2. its derivative is given by:

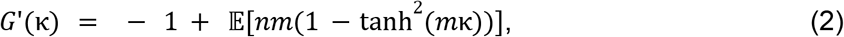

and empirical averages converge to the expectations in *G*(*κ*) and *G*′(*κ*) by the law of large numbers.

**Proof:** We will perform the proof in three steps. First, we show that *G*(*κ*) is a proper function that has finite values for any finite *κ* and the sample averages converge to a distribution centered on the population mean with a vanishing variance. Second, using dominated convergence, we show that it is differentiable, whereas its derivative, given by Eq. (2), is a proper function. We show that averages over neurons converge to *G*′(*κ*). Finally, using dominated convergence, we prove the continuity of *G*′(*κ*) and conclude the proof.

First, since tanh(·) is a bounded function that satisfies the relationship | tanh(*x*)| < 1 for *x* ∈ ℝ, we can bound the expression inside the expectation in Eq. (1) from above as: |*n* tanh(*mκ*)| ≤ |*n*|, *i.e*., the integrand in Eq. (1) for *G*(*κ*) is dominated by |*n*|. Since we assume that E[*n*^2^] E[*n*] < ∞, it follows from the Cauchy-Schwarz inequality that 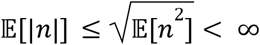. Therefore, the integrand is integrable, and *G*(*κ*) is a well-defined real-valued function for all *κ* ∈ ℝ. Moreover, since *Var*[*n* tanh(*mκ*)] ≤ E[*n*^2^tanh^2^(*mκ*)] ≤ E[*n*^2^], in the limit *N* → ∞, the empirical average over neurons converges to *G*(*κ*) by the law of large numbers.

Second, we can compute the derivative of *G*(*κ*) following 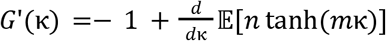. To justify switching the derivative and expectation, we first consider the function *h*(*κ*) = *nm*(1 – tanh^2^ (*mκ*)). Then, similar to the first step, we observe that |*h*(*κ*)| ≤ |*nm*|. Once again using the Cauchy-Schwarz inequality, we find that 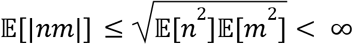 by definition. Hence, by the dominated convergence theorem, we can switch the expectation and the derivative, which leads to Eq. (2). Moverover, we have the set of inequalities following a similar arguments as before: 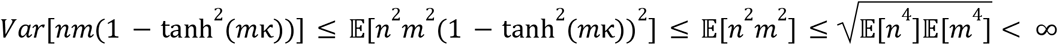. Hence, in the limit *N* → ∞, the empirical average over neurons converges to *G*′(*κ*) by the law of large numbers.

As the final step, since the integrand, *h*(*κ*), is continuous and dominated by an integrable function, *G*′(*κ*) is continuous.

**Proposition A1** establishes the regularity conditions, under which the latent dynamical system given in Eq. (1) converges to the mean-field limit as more samples are added and the flow map is well-defined as a function of *κ* under the expectation over (*m, n*).

#### 2.2. Bistability of the latent dynamical system

Having discussed the regularity of the resulting latent dynamical system in the mean-field limit, we now state a proposition describing the qualitative behavior of this latent dynamical system:

##### Proposition A2

**(Bistable latent dynamical system)**

Let *κ* be a latent dynamical system governed by the flow map, *G*(*κ*), defined in Eq. (1) in the limit *N* → ∞. Let the pair (*m, n*) be sampled from a two-dimensional Gaussian distribution that has zero mean and finite covariance. Let this distribution satisfy the condition E[*nm*] > 1. Then, *G*(*κ*) defines a bistable latent dynamical system with a repeller fixed-point at the origin (*κ* = 0), whereas its two attractive fixed-points reside at *κ* =± *c* for some *c* > 0.

**Proof:** This is a special case of the results shown in Section 4 of Ref. 7 for rank-one RNNs. Here, we provide a simple and intuitive proof for this special case. First, we prove that for E[*nm*] > 1, the origin becomes a repeller. Since we assumed no inputs and no biases, the origin of the latent dynamical system is always a fixed-point with *G*(*κ* = 0) = 0. In the absence of any excitation or inhibition between neurons, *i.e*., ***m*** = ***n*** = 0, the origin is an attractor since *G*′(*κ* = 0) =−1, where *G*′(*κ*) refers to the derivative with respect to the latent variable *κ*. Since our interest is to design a bistable LPU (*e.g*., as in those in **Fig. 2**), this requires the origin to turn into a repeller fixed-point, *i.e*., *G*′(*κ* = 0) > 0. To study this, we consider the behavior of the latent dynamical system around the origin for small *κ* such that:

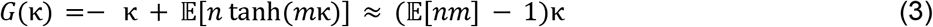

where we note E[*nm*] as the covariance of the random variable pair (*m, n*), as both have zero means by assumption. Inspecting the linearized dynamics in Eq. (3) reveals that, for E[*nm*] > 1, the origin becomes a repeller as *G*′(*κ*) = E[*nm*] − 1 > 0.

To prove that this system shows bistable dynamics, *i.e*., has only two more attractive fixed-points, we now consider the derivative of the flow map that follows *h*(*κ*) = *G*′(*κ*) = − 1 + E[*nm*(1 – tanh^2^ (*mκ*))]. The claim is that *h*(*κ*) is a decreasing function of *κ* > 0, where *h*(*κ* = 0) = E[*nm*] − 1 and *h*(*κ* → ∞) → − 1. To prove this, we take the derivative *h*′(*κ*) =− 2E[*nm*^2^ tanh^2^(*mκ*)(1 − tanh (*mk*))]. Using the tower property, we can compute the expectation:

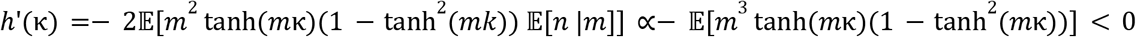

This follows from the fact that the expression inside the expectation is a positive function of any *m* ∈ ℝ and for *κ* > 0, thereby *h*(*κ*) is a decreasing function for *κ* > 0. This in turn means that, for E[*nm*] − 1 > 0, *G*′(*κ*) changes sign only once and therefore *G*(*κ*) has only one local maximum and no local minima for *κ* > 0. Then, there is only one *c* > 0 for which *G*(*c*) = 0 and *G*′(*c*) < 0, *i.e*., there is an attractive fixed-point at *κ* = *c*. Since *G*(*κ*) is odd-symmetric, a similar condition holds for *κ* =− *c*, making this system have bistable dynamics. On the other hand, when E[*nm*] < 1, then *G*′(*κ*) is always negative. In this case, no local maximum exists for *G*(*κ*), which decays monotonically for *κ* > 0 and only has an attractive fixed-point at the origin, concluding the proof.

In practice, using the *GL*_*K*_ (ℝ) symmetry discussed in the main text, it is possible to define *c* = 1 without loss of generality. Specifically, we can always scale ***n*** and ***m*** with appropriate factors as long as 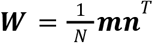 remains invariant. After setting this scale, there are two free parameters of the Gaussian distribution: i) the joint scale of the parameters σ = σ_*m*_ σ_*n*_, and ii) the correlation, ρ, between the variables. These variables jointly define E[*nm*] = σρ and will be tuned below to design bistable LPUs with predefined timescales.

#### 2.3. Designing the latent processing unit with target timescales

To design the LPU with a target timescale, we now return to the linearization around *κ* in Eq. (3), which defines a local decay/growth-rate following:

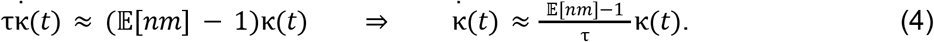

Here, we can define 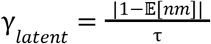 as the latent decay (when E[*nm*] < 1) or growth (when E[*nm*] > 1) rate. Noting that E[*nm*] = σρ (and enforcing σρ > 1 since we are interested in bistable dynamics), we can design a latent dynamical system with a target growth rate as:

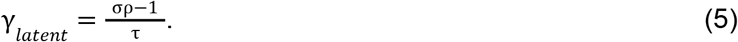

By choosing σρ → 1 + ϵ with ϵ ≪ 1, or σρ → ∞, it is theoretically possible to design a bistable latent dynamical system with a growth rate γ_*latent*_ ∈ (0, ∞). However, how robust are these timescales when networks have a finite number of neurons, *i.e*., outside of the mean-field limit?

To study this question, we return to the finite-neuron limit. Instead of the mean-field limit, E[*nm*], we focus on the empirical covariance *C*(***m, n***) = ***n***^*T*^ ***m***/*N* that sets the empirical timescale in a network with finite number of neurons following:

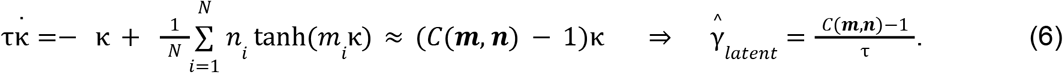

Notably, this value can become negative (hence no bistable dynamics) due to the fluctuations of *C*(***m, n***) around E[*mn*] even though the latter is positive. Next, we compute the mean and the variance for this distribution as: 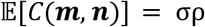 and 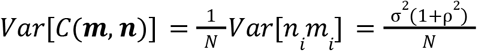. Then, the normalized error in the growth rates of the designed LPUs follow the relationship:

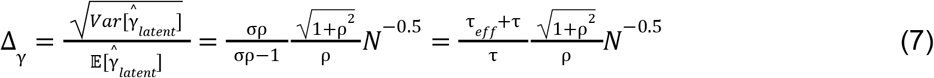

For notational convenience, we defined the latent timescale as 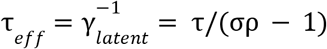. The scaling relationship already provides a clear prediction. The error in the target growth rate is minimized for ρ = 1, *i.e*., when the variables are perfectly correlated resulting in the symmetric Hopfield limit.

Given the presumed separation of neural and kinematic (*i.e*., latent) timescales, we are particularly interested in the limit β = τ_*eff*_ /τ ≫ 1, which leads to the following scaling relationship:

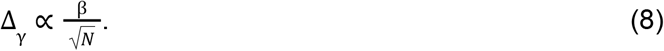

This relationship becomes particularly relevant for biological neural networks, in which biological neurons have *O*(*ms*) timescales, whereas behaviors often take place in seconds-long timescales. For example, decision-making processes often require extended periods of evidence accumulation before committing to one of two stable states (here, fixed-points located at *κ* =± 1). In such cases, the variable *κ* must maintain a transitory position near the center until sufficient evidence accumulates, rather than prematurely settling into either attractor state. In such a case, τ_*eff*_ ≫ τ is a necessity. For instance, to achieve τ_*eff*_ /τ ~ 10^2^ with Δ_γ_ ~ 10% errors, the network should have about one million neurons. Therefore, while **Theorem 1** guarantees LPUs to model any latent dynamical system in the limit *N* → ∞, the scaling relationship in Eq. (8) quantifies the need for large networks when designing LPUs with timescales that are much larger than the intrinsic timescales of the neurons (cf. **Theorem 2**).

#### 2.4. Empirical reconstruction of LPUs from partial observation

The theoretical analysis above can be extended to an empirically relevant consideration: LPU reconstruction under partial observations of neurons. Specifically, the scaling relationship in Eq. (7) applies to the scenario *N*_*obs*_ ≪ *N*_*obs*_ → ∞, where *N* refers to the number of observed neurons. In this limit, partial observations from the pairs of encoding and embedding weights, (*n*_*i*_, *m*_*i*_) for *i* = 1, …, *N*, would be equivalent to designing a network with a smaller number of neurons using the ground truth probability distribution. Below, we perform an extension of this analysis for a general *N*_*obs*_, *e.g*., incorporating the limit *N*_*obs*_ ≈ *N* for a finite *N*.

First, we introduce the reconstruction procedure for the LPUs under partial observation of neurons, but for an idealized limit that allows access to the (*n*_*i*_, *m*_*i*_) variables belonging to the observed neurons. This scenario is unrealistic, as partial observation often involves only the neural activities, *r*_*i*_ (*t*), which is not necessarily enough to pin down these parameters (for an example, see **Fig. 5**). Instead, for now, we unrealistically assume that the functional connections, *i.e*., the pairs (*n*_*i*_, *m*_*i*_), are perfectly estimated through some means. Notably, though often one focuses on reconstructing the dynamics, enforcing that the functional connections are properly reconstructed is one important aspect of how RNNs enable causal predictions when trained to reproduce brain recordings^5^. In the end, even under this unrealistically informative assumption, we will show that large-scale observations are necessary to faithfully reproduce neural dynamics.

For a group of observed neurons and a one-dimensional LPU in Eq. (1), we reconstruct the recurrent weights by simply using their ground truth encoding and embedding values and replace the division by *N* with a division by *N*_*obs*_ such that 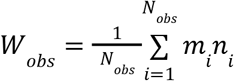. where neurons *i* = 1, …, *N*_*obs*_ are assumed to be observed. If *N*_*obs*_ = *N*, this estimation procedure reconstructs the ground truth weight matrix. Then, we can reconstruct the latent flow map as:

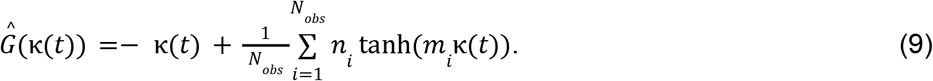

This, on average, reconstructs the time dynamics of the original LPU following:

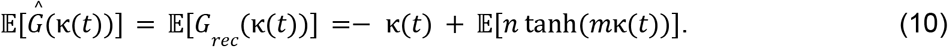

Hence, the estimator 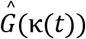 is an unbiased estimator of the full population dynamics in the mean-field limit. This idealized estimation procedure can illustrate a fundamental problem with reconstructing latent dynamics using subsampled neuronal populations even when the ground truth dynamical system equations are known: The reconstructed LPU at a particular reconstruction instance can have high variations from the ground truth if the neurons are extremely sparsely sampled. To demonstrate this, we derive the theoretical errors between the estimated timescales as 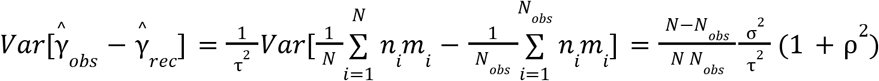, where 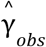 refers to the reconstructed latent growth rate under partial observation, whereas 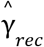 refers to the actual growth of the network with *N* neurons. This leads to the normalized estimation error:

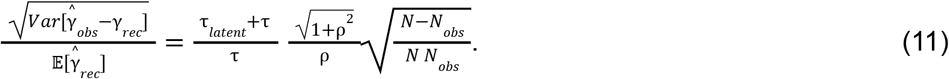

In the limit *N* ≫ *N*_*obs*_, this equation converges to Eq. (7), as expected. In words, the reconstruction of the latent growth rates scales as 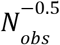 when only a sparse subset of neurons are observed. In the other limit, where Δ*N* = *N* − *N*_*obs*_ ≪ *N, i.e*., a large population of neurons are observed, we obtain a different scaling 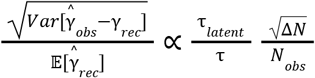. The added neurons now rapidly decrease the reconstruction errors following Δ*N* → 0 and 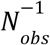 scaling.

### 3. Conclusion

Our findings in this supplementary note demonstrate that large-scale neural recordings are essential for accurately estimating bistable dynamics, which represent one of the most fundamental phenomena in dynamical systems theory^8^. It is nowadays common for dynamical models trained on recordings from a small subset of neurons to capture the observed neural dynamics. However, the example scenario we considered in this section reveals that limited/partial observation of neurons introduces substantial variance that may obscure the link between LPUs and the true functional connections (here, the (*m, n*) pairs). This insight has implications for a recent direction in system neuroscience, in which training RNNs on neural recordings could capture their functional connections. Then, these functional connections can be studied to uncover inter-area communication patterns^5^. Recent work has shown that such models could lead to spurious interactions inferred between brain regions^9^, whereas our analysis here indicates that training such dynamical models requires, first and foremost, comprehensive large-scale recordings. In our example scenario studied here, the latter was necessary in order to achieve reliable reconstruction of latent dynamics while enforcing true functional interactions, *i.e*., ground truth (*m, n*) pairs.

## REFERENCES

1. Duncker, L. & Sahani, M. Dynamics on the manifold: Identifying computational dynamical activity from neural population recordings. Curr Opin Neurobiol 70, 163–170 (2021).

2. Urai, A. E., Doiron, B., Leifer, A. M. & Churchland, A. K. Large-scale neural recordings call for new insights to link brain and behavior. Nat Neurosci 25, 11–19 (2022).

3. Langdon, C., Genkin, M. & Engel, T. A. A unifying perspective on neural manifolds and circuits for cognition. Nat Rev Neurosci 24, 363–377 (2023).

4. Deitch, D., Rubin, A. & Ziv, Y. Representational drift in the mouse visual cortex. Curr Biol 31, 4327–4339.e6 (2021).

5. Marks, T. D. & Goard, M. J. Stimulus-dependent representational drift in primary visual cortex. Nat Commun 12, 5169 (2021).

6. Rokni, U., Richardson, A. G., Bizzi, E. & Seung, H. S. Motor learning with unstable neural representations. Neuron 54, 653–666 (2007).

7. Driscoll, L. N., Duncker, L. & Harvey, C. D. Representational drift: Emerging theories for continual learning and experimental future directions. Curr Opin Neurobiol 76, 102609 (2022).

8. Ebrahimi, S. et al. Emergent reliability in sensory cortical coding and inter-area communication. Nature 605, 713–721 (2022).

9. Driscoll, L. N., Pettit, N. L., Minderer, M., Chettih, S. N. & Harvey, C. D. Dynamic Reorganization of Neuronal Activity Patterns in Parietal Cortex. Cell 170, 986–999.e16 (2017).

10. Qin, S. et al. Coordinated drift of receptive fields in Hebbian/anti-Hebbian network models during noisy representation learning. Nat Neurosci 26, 339–349 (2023).

11. Mastrogiuseppe, F. & Ostojic, S. Linking Connectivity, Dynamics, and Computations in Low-Rank Recurrent Neural Networks. Neuron 99, 609–623.e29 (2018).

12. Yuste, R. From the neuron doctrine to neural networks. Nat Rev Neurosci 16, 487–497 (2015).

13. Williams, A. H. et al. Unsupervised Discovery of Demixed, Low-Dimensional Neural Dynamics across Multiple Timescales through Tensor Component Analysis. Neuron 98, 1099–1115.e8 (2018).

14. Schneider, S., Lee, J. H. & Mathis, M. W. Learnable latent embeddings for joint behavioural and neural analysis. Nature 617, 360–368 (2023).

15. Pandarinath, C. et al. Inferring single-trial neural population dynamics using sequential auto-encoders. Nat Methods 15, 805–815 (2018).

16. DePasquale, B., Sussillo, D., Abbott, L. F. & Churchland, M. M. The centrality of population-level factors to network computation is demonstrated by a versatile approach for training spiking networks. Neuron 111, 631–649.e10 (2023).

17. Zimnik, A. J. et al. Identifying Interpretable Latent Factors with Sparse Component Analysis. bioRxiv (2024) doi:10.1101/2024.02.05.578988.

18. Versteeg, C., Sedler, A. R., McCart, J. D. & Pandarinath, C. Expressive dynamics models with nonlinear injective readouts enable reliable recovery of latent features from neural activity. ArXiv (2023).

19. Jazayeri, M. & Ostojic, S. Interpreting neural computations by examining intrinsic and embedding dimensionality of neural activity. Curr Opin Neurobiol 70, 113–120 (2021).

20. Glaser, J., Whiteway, M., Cunningham, J. P., Paninski, L. & Linderman, S. Recurrent Switching Dynamical Systems Models for Multiple Interacting Neural Populations. Advances in Neural Information Processing Systems 33, 14867–14878 (2020).

21. Cunningham, J. P. & Yu, B. M. Dimensionality reduction for large-scale neural recordings. Nature Neuroscience 17, 1500–1509 (2014).

22. Bjerke, M. et al. Understanding Neural Coding on Latent Manifolds by Sharing Features and Dividing Ensembles. in The Eleventh International Conference on Learning Representations (2022).

23. Kohli, D., Nieuwenhuis, J. S., Cloninger, A., Mishne, G. & Narain, D. RATS: Unsupervised manifold learning using low-distortion alignment of tangent spaces. bioRxiv (2024) doi:10.1101/2024.10.31.621292.

24. Degenhart, A. D. et al. Stabilization of a brain-computer interface via the alignment of low-dimensional spaces of neural activity. Nat Biomed Eng 4, 672–685 (2020).

25. Fortunato, C. et al. Nonlinear manifolds underlie neural population activity during behaviour. bioRxiv (2024) doi:10.1101/2023.07.18.549575.

26. Perich, M. G., Narain, D. & Gallego, J. A. A neural manifold view of the brain. Nat Neurosci 28, 1582–1597 (2025).

27. Gardner, R. J. et al. Toroidal topology of population activity in grid cells. Nature 602, 123–128 (2022).

28. O’Shea, D. J. et al. Direct neural perturbations reveal a dynamical mechanism for robust computation. bioRxiv 2022.12.16.520768 (2022) doi:10.1101/2022.12.16.520768.

29. Abbaspourazad, H., Erturk, E., Pesaran, B. & Shanechi, M. M. Dynamical flexible inference of nonlinear latent factors and structures in neural population activity. Nat Biomed Eng 8, 85–108 (2024).

30. Linderman, S. et al. Bayesian Learning and Inference in Recurrent Switching Linear Dynamical Systems. in Artificial Intelligence and Statistics 914–922 (PMLR, 2017).

31. Vinograd, A., Nair, A., Kim, J. H., Linderman, S. W. & Anderson, D. J. Causal evidence of a line attractor encoding an affective state. Nature 634, 910–918 (2024).

32. Hu, A. et al. Modeling Latent Neural Dynamics with Gaussian Process Switching Linear Dynamical Systems. ArXiv (2025).

33. Valente, A., Pillow, J. W. & Ostojic, S. Extracting computational mechanisms from neural data using low-rank RNNs. Advances in Neural Information Processing Systems 35, 24072–24086 (2022).

34. Dubreuil, A., Valente, A., Beiran, M., Mastrogiuseppe, F. & Ostojic, S. The role of population structure in computations through neural dynamics. Nat Neurosci 25, 783–794 (2022).

35. Beiran, M., Dubreuil, A., Valente, A., Mastrogiuseppe, F. & Ostojic, S. Shaping Dynamics With Multiple Populations in Low-Rank Recurrent Networks. Neural Comput 33, 1572–1615 (2021).

36. Langdon, C. & Engel, T. A. Latent circuit inference from heterogeneous neural responses during cognitive tasks. Nat Neurosci 28, 665–675 (2025).

37. Liu, M., Nair, A., Coria, N., Linderman, S. W. & Anderson, D. J. Encoding of female mating dynamics by a hypothalamic line attractor. Nature 634, 901–909 (2024).

38. Rigotti, M. et al. The importance of mixed selectivity in complex cognitive tasks. Nature 497, 585–590 (2013).

39. Stringer, C., Pachitariu, M., Steinmetz, N., Carandini, M. & Harris, K. D. High-dimensional geometry of population responses in visual cortex. Nature 571, 361–365 (2019).

40. Manley, J. et al. Simultaneous, cortex-wide dynamics of up to 1 million neurons reveal unbounded scaling of dimensionality with neuron number. Neuron 112, 1694–1709.e5 (2024).

41. Pashakhanloo, F. & Koulakov, A. Stochastic Gradient Descent-Induced Drift of Representation in a Two-Layer Neural Network. in International Conference on Machine Learning 27401–27419 (PMLR, 2023).

42. Schäfer, A. M. & Zimmermann, H.-G. Recurrent Neural Networks are universal approximators. Int J Neural Syst 17, 253–263 (2007).

43. Multilayer feedforward networks with a nonpolynomial activation function can approximate any function. Neural Networks 6, 861–867 (1993).

44. Nair, A. et al. An approximate line attractor in the hypothalamus encodes an aggressive state. Cell 186, 178–193.e15 (2023).

45. Semedo, J. D., Zandvakili, A., Machens, C. K., Yu, B. M. & Kohn, A. Cortical Areas Interact through a Communication Subspace. Neuron 102, 249–259.e4 (2019).

46. Posani, L., Wang, S., Muscinelli, S. P., Paninski, L. & Fusi, S. Rarely categorical, always high-dimensional: how the neural code changes along the cortical hierarchy. bioRxiv (2025) doi:10.1101/2024.11.15.623878.

47. Schmutz, V. et al. High-dimensional neuronal activity from low-dimensional latent dynamics: a solvable model. bioRxiv (2025) doi:10.1101/2025.06.03.657632.

48. Moser, M.-B., Rowland, D. C. & Moser, E. I. Place Cells, Grid Cells, and Memory. Cold Spring Harb Perspect Biol 7, a021808 (2015).

49. Priebe, N. J. Mechanisms of Orientation Selectivity in the Primary Visual Cortex. Annu Rev Vis Sci 2, 85–107 (2016).

50. Bounds, H. A. & Adesnik, H. Network influence determines the impact of cortical ensembles on stimulus detection. Neuron (2025) doi:10.1016/j.neuron.2025.04.023.

51. Zatka-Haas, P., Steinmetz, N. A., Carandini, M. & Harris, K. D. Sensory coding and the causal impact of mouse cortex in a visual decision. (2021) doi:10.7554/eLife.63163.

52. Steinmetz, N. A., Zatka-Haas, P., Carandini, M. & Harris, K. D. Distributed coding of choice, action and engagement across the mouse brain. Nature 576, 266–273 (2019).

53. Rule, M. E. et al. Stable task information from an unstable neural population. Elife 9, (2020).

54. Vyas, S., Golub, M. D., Sussillo, D. & Shenoy, K. V. Computation Through Neural Population Dynamics. Annu Rev Neurosci 43, 249–275 (2020).

55. Donoho, D. L. Compressed Sensing. (2004).

56. Ganguli, S. & Sompolinsky, H. Compressed sensing, sparsity, and dimensionality in neuronal information processing and data analysis. Annu Rev Neurosci 35, 485–508 (2012).

57. Ganguli, S. & Sompolinsky, H. Short-term memory in neuronal networks through dynamical compressed sensing. Advances in Neural Information Processing Systems 23, (2010).

58. Thibeault, V., Allard, A. & Desrosiers, P. The low-rank hypothesis of complex systems. Nature Physics 20, 294–302 (2024).

59. Barak, O. Recurrent neural networks as versatile tools of neuroscience research. Curr Opin Neurobiol 46, 1–6 (2017).

60. Sussillo, D. & Barak, O. Opening the black box: low-dimensional dynamics in high-dimensional recurrent neural networks. Neural Comput 25, 626–649 (2013).

61. Krause, R., Cook, M., Kollmorgen, S., Mante, V. & Indiveri, G. Operative dimensions in unconstrained connectivity of recurrent neural networks. Advances in Neural Information Processing Systems 35, 17073–17085 (2022).

62. Gao, P. et al. A theory of multineuronal dimensionality, dynamics and measurement. bioRxiv 214262 (2017) doi:10.1101/214262.

63. Zheng, J. & Meister, M. The unbearable slowness of being: Why do we live at 10 bits/s? Neuron 113, 192–204 (2025).

64. Zeraati, R., Levina, A., Macke, J. H. & Gao, R. Neural timescales from a computational perspective. (2024).

65. Advani, M., Lahiri, S. & Ganguli, S. Statistical mechanics of complex neural systems and high dimensional data. J. Stat. Mech. 2013, P03014 (2013).

66. Yang, G. R., Joglekar, M. R., Song, H. F., Newsome, W. T. & Wang, X.-J. Task representations in neural networks trained to perform many cognitive tasks. Nature Neuroscience 22, 297–306 (2019).

67. Abbott, L. F. & Dayan, P. The effect of correlated variability on the accuracy of a population code. Neural Comput 11, 91–101 (1999).

68. Masse, N. Y., Yang, G. R., Song, H. F.Wang, X.-J. & Freedman, D. J. Circuit mechanisms for the maintenance and manipulation of information in working memory. Nat Neurosci 22, 1159–1167 (2019).

69. Ayed, I., de Bézenac, E., Pajot, A., Brajard, J. & Gallinari, P. Learning Dynamical Systems from Partial Observations. (2019).

70. Das, A. & Fiete, I. R. Systematic errors in connectivity inferred from activity in strongly recurrent networks. Nat Neurosci 23, 1286–1296 (2020).

71. Qian, W., Zavatone-Veth, J. A., Ruben, B. S. & Pehlevan, C. Partial observation can induce mechanistic mismatches in data-constrained models of neural dynamics. in The Thirty-eighth Annual Conference on Neural Information Processing Systems (2024).

72. Rumyantsev, O. I. et al. Fundamental bounds on the fidelity of sensory cortical coding. Nature 580, 100–105 (2020).

73. Hazon, O. et al. Noise correlations in neural ensemble activity limit the accuracy of hippocampal spatial representations. Nat Commun 13, 4276 (2022).

74. Moreno-Bote, R. et al. Information-limiting correlations. Nature Neuroscience 17, 1410–1417 (2014).

75. Kira, S., Safaai, H., Morcos, A. S., Panzeri, S. & Harvey, C. D. A distributed and efficient population code of mixed selectivity neurons for flexible navigation decisions. Nat Commun 14, 2121 (2023).

76. Gao, P. & Ganguli, S. On simplicity and complexity in the brave new world of large-scale neuroscience. Curr Opin Neurobiol 32, 148–155 (2015).

77. Clark, D. G., Marschall, O., van Meegen, A. & Litwin-Kumar, A. Connectivity Structure and Dynamics of Nonlinear Recurrent Neural Networks. Phys Rev X 15, (2025).

78. Gosztolai, A., Peach, R. L., Arnaudon, A., Barahona, M. & Vandergheynst, P. MARBLE: interpretable representations of neural population dynamics using geometric deep learning. Nature Methods 22, 612–620 (2025).

79. Kim, T. D. et al. Flow-field inference from neural data using deep recurrent networks. bioRxiv (2023) doi:10.1101/2023.11.14.567136.

80. Ziv, Y. et al. Long-term dynamics of CA1 hippocampal place codes. Nat Neurosci 16, 264–266 (2013).

81. Eliasmith, C. & Anderson, C. H. Neural Engineering: Computation, Representation, and Dynamics in Neurobiological Systems. (MIT Press (MA), 2003).

82. Dorkenwald, S. et al. Neuronal wiring diagram of an adult brain. Nature 634, 124–138 (2024).

83. Beiran, M. & Litwin-Kumar, A. Prediction of neural activity in connectome-constrained recurrent networks. Nature Neuroscience 28, 2561–2574 (2025).

84. Perich, M. G. et al. Inferring brain-wide interactions using data-constrained recurrent neural network models. bioRxiv 2020.12.18.423348 (2021) doi:10.1101/2020.12.18.423348.

85. Perich, M. G. & Rajan, K. Rethinking brain-wide interactions through multi-region ‘network of networks’ models. Curr Opin Neurobiol 65, 146–151 (2020).

86. Hopfield, J. J. Neural networks and physical systems with emergent collective computational abilities. Proceedings of the National Academy of Sciences 79, 2554–2558 (1982).

87. Burak, Y. & Fiete, I. R. Fundamental limits on persistent activity in networks of noisy neurons. Proceedings of the National Academy of Sciences 109, 17645–17650 (2012).

88. Rajan, K., Harvey, C. D. & Tank, D. W. Recurrent Network Models of Sequence Generation and Memory. Neuron 90, 128–142 (2016).

89. Hermansen, E., Klindt, D. A. & Dunn, B. A. Uncovering 2-D toroidal representations in grid cell ensemble activity during 1-D behavior. Nat Commun 15, 5429 (2024).

90. Sompolinsky, H., Crisanti, A. & Sommers, H. J. Chaos in random neural networks. Phys Rev Lett 61, 259–262 (1988).

91. Sadeh, S. & Clopath, C. Contribution of behavioural variability to representational drift. Elife 11, (2022).

92. Micou, C. & O’Leary, T. Representational drift as a window into neural and behavioural plasticity. Curr Opin Neurobiol 81, 102746 (2023).

93. Schuessler, F., Mastrogiuseppe, F., Dubreuil, A., Ostojic, S. & Barak, O. The interplay between randomness and structure during learning in RNNs. Advances in Neural Information Processing Systems 33, 13352–13362 (2020).

94. Maheswaranathan, N., Williams, A. H., Golub, M. D., Ganguli, S. & Sussillo, D. Universality and individuality in neural dynamics across large populations of recurrent networks. Adv Neural Inf Process Syst 2019, 15629–15641 (2019).

95. Podlaski, W. F. & Machens, C. K. Approximating nonlinear functions with latent boundaries in low-rank excitatory-inhibitory spiking networks. Neural Computation 36.5, 803–857 (2024).

96. Pezon, L., Schmutz, V. & Gerstner, W. Linking neural manifolds to circuit structure in recurrent networks. Neuron 114, 1682–1694.e21 (2026).

97. Safaie, M. et al. Preserved neural dynamics across animals performing similar behaviour. Nature 623, 765–771 (2023).

98. Jackson, A. & Hall, T. M. Decoding Local Field Potentials for Neural Interfaces. https://ieeexplore.ieee.org/document/7742994/.

99. Werbos, P. J. Backpropagation through time: what it does and how to do it. Proc. IEEE Inst. Electr. Electron. Eng. 78, 1550–1560 (1990).

100. Karniol-Tambour, O. et al. Modeling state-dependent communication between brain regions with switching nonlinear dynamical systems. in The Twelfth International Conference on Learning Representations (2023).

101. Kandel, E. R., Schwartz, J. H. & Jessell, T. M. Principles of Neural Science. (1991).

102. Dinc, F., Shai, A., Schnitzer, M. & Tanaka, H. CORNN: Convex optimization of recurrent neural networks for rapid inference of neural dynamics. (2023).

103. Mukamel, E. A., Nimmerjahn, A. & Schnitzer, M. J. Automated analysis of cellular signals from large-scale calcium imaging data. Neuron 63, 747–760 (2009).

104. Garreta, R. & Moncecchi, G. Learning Scikit-Learn: Machine Learning in Python. (Packt Pub Limited, 2013).

105. Lee, J. M. Introduction to Smooth Manifolds. (Springer New York).

106. Acosta, F., Sanborn, S., Duc, K. D., Madhav, M. & Miolane, N. Quantifying Extrinsic Curvature in Neural Manifolds. (2022).

107. Montúfar, G., Pascanu, R., Cho, K. & Bengio, Y. On the Number of Linear Regions of Deep Neural Networks. (2014).

108. Pascanu, R., Montufar, G. & Bengio, Y. On the number of response regions of deep feed forward networks with piece-wise linear activations. (2013).

109. Huber, P. J. Robust Statistics. (John Wiley & Sons, 2004).

## References

1. Boerlin, M., Machens, C. K. & Denève, S. Predictive coding of dynamical variables in balanced spiking networks. PLoS Comput Biol 9, e1003258 (2013).

2. Eliasmith, C. A unified approach to building and controlling spiking attractor networks. Neural Comput 17, 1276–1314 (2005).

3. Eliasmith, C. & Anderson, C. H. Neural Engineering: Computation, Representation, and Dynamics in Neurobiological Systems. (MIT Press (MA), 2003).

4. Schneider, S., Lee, J. H. & Mathis, M. W. Learnable latent embeddings for joint behavioural and neural analysis. Nature 617, 360–368 (2023).

5. Butts, D. A. & Goldman, M. S. Tuning curves, neuronal variability, and sensory coding. PLoS Biol 4, e92 (2006).

6. Independent component analysis: algorithms and applications. Neural Networks 13, 411–430 (2000).

7. Hyvärinen, A. & Pajunen, P. Nonlinear independent component analysis: Existence and uniqueness results. Neural Netw 12, 429–439 (1999).

8. Zhou, D. & Wei, X.-X. Learning identifiable and interpretable latent models of high-dimensional neural activity using pi-VAE. Advances in Neural Information Processing Systems 33, 7234–7247 (2020).

9. Mastrogiuseppe, F. & Ostojic, S. Linking Connectivity, Dynamics, and Computations in Low-Rank Recurrent Neural Networks. Neuron 99, 609–623.e29 (2018).

10. Valente, A., Pillow, J. W. & Ostojic, S. Extracting computational mechanisms from neural data using low-rank RNNs. Advances in Neural Information Processing Systems 35, 24072–24086 (2022).

11. Dubreuil, A., Valente, A., Beiran, M., Mastrogiuseppe, F. & Ostojic, S. The role of population structure in computations through neural dynamics. Nat Neurosci 25, 783–794 (2022).

12. Beiran, M., Dubreuil, A., Valente, A., Mastrogiuseppe, F. & Ostojic, S. Shaping Dynamics With Multiple Populations in Low-Rank Recurrent Networks. Neural Comput 33, 1572–1615 (2021).

13. Glaser, J., Whiteway, M., Cunningham, J. P., Paninski, L. & Linderman, S. Recurrent Switching Dynamical Systems Models for Multiple Interacting Neural Populations. Advances in Neural Information Processing Systems 33, 14867–14878 (2020).

14. Nair, A. et al. An approximate line attractor in the hypothalamus encodes an aggressive state. Cell 186, 178–193.e15 (2023).

15. Linderman, S. et al. Bayesian Learning and Inference in Recurrent Switching Linear Dynamical Systems. in Artificial Intelligence and Statistics 914–922 (PMLR, 2017).

16. Vinograd, A., Nair, A., Kim, J. H., Linderman, S. W. & Anderson, D. J. Causal evidence of a line attractor encoding an affective state. Nature 634, 910–918 (2024).

17. Hu, A. et al. Modeling Latent Neural Dynamics with Gaussian Process Switching Linear Dynamical Systems. ArXiv (2025).

18. Liu, M., Nair, A., Coria, N., Linderman, S. W. & Anderson, D. J. Encoding of female mating dynamics by a hypothalamic line attractor. Nature 634, 901–909 (2024).

19. Abbaspourazad, H., Erturk, E., Pesaran, B. & Shanechi, M. M. Dynamical flexible inference of nonlinear latent factors and structures in neural population activity. Nat Biomed Eng 8, 85–108 (2024).

20. Stringer, C., Pachitariu, M., Steinmetz, N., Carandini, M. & Harris, K. D. High-dimensional geometry of population responses in visual cortex. Nature 571, 361–365 (2019).

21. Manley, J. et al. Simultaneous, cortex-wide dynamics of up to 1 million neurons reveal unbounded scaling of dimensionality with neuron number. Neuron 112, 1694–1709.e5 (2024).

## References

1. Qian, W., Zavatone-Veth, J. A., Ruben, B. S. & Pehlevan, C. Partial observation can induce mechanistic mismatches in data-constrained models of neural dynamics. in The Thirty-eighth Annual Conference on Neural Information Processing Systems (2024).

2. Ayed, I., de Bézenac, E., Pajot, A., Brajard, J. & Gallinari, P. Learning Dynamical Systems from Partial Observations. (2019).

3. Dinc, F., Shai, A., Schnitzer, M. & Tanaka, H. CORNN: Convex optimization of recurrent neural networks for rapid inference of neural dynamics. (2023).

4. Perich, M. G. & Rajan, K. Rethinking brain-wide interactions through multi-region ‘network of networks’ models. Curr Opin Neurobiol 65, 146–151 (2020).

5. Perich, M. G. et al. Inferring brain-wide interactions using data-constrained recurrent neural network models. bioRxiv 2020.12.18.423348 (2021) doi:10.1101/2020.12.18.423348.

6. Manley, J. et al. Simultaneous, cortex-wide dynamics of up to 1 million neurons reveal unbounded scaling of dimensionality with neuron number. Neuron 112, 1694–1709.e5 (2024).

7. Beiran, M., Dubreuil, A., Valente, A., Mastrogiuseppe, F. & Ostojic, S. Shaping Dynamics With Multiple Populations in Low-Rank Recurrent Networks. Neural Comput 33, 1572–1615 (2021).

8. Strogatz, S. H. Nonlinear Dynamics and Chaos: With Applications to Physics, Biology, Chemistry, and Engineering. (CRC Press, 2018).

9. Dowling, M. & Savin, C. Nonlinear multiregion neural dynamics with parametric impulse response communication channels. in The Thirteenth International Conference on Learning Representations (2024).

